# Decoding the pituitary gonadotrope regulatory architecture governing the preovulatory surge *in vivo*

**DOI:** 10.1101/2025.11.18.689023

**Authors:** C. Le Ciclé, G. Garrel, S. Chauvin, F. Giton, O. Taboureau, F. Miralles, J. Cohen Tannoudji, D. L’Hôte

## Abstract

Despite the global burden of female infertility, the molecular control of ovulation remains unclear. Pituitary gonadotropes tune hormonal output to regulate ovarian function, yet the molecular mechanisms that control their function during the sexual cycle remain poorly understood. Using state-of-the-art single-nucleus multiomic profiling of the female rat pituitary across the estrus cycle, we uncover unexpected cyclic epigenetic and transcriptional remodelling in gonadotrope cells, particularly during the preovulatory surge. This plasticity is driven by a biphasic gene regulatory network switch that orchestrates large-scale reorganisation of the secretory machinery. We further demonstrate that, contrary to the prevailing model, the LH surge does not depend on *Lhb* transcriptional upregulation. Instead, the FSH surge arises from dynamic *Fshb* transcription mediated by a complex combination of newly identified enhancers and key transcription factors. Overall, our findings define a new molecular framework for gonadotropin surge regulation and provide a foundational resource for understanding female reproductive disorders.

## Introduction

Rhythmicity is an essential property of living systems, shaping processes from circadian oscillations to seasonal reproductive cycles. These temporal programs synchronise cellular physiology with environmental cues, and their disruption has major physiological consequences. In mammals, one of the most tightly regulated rhythms is the female sexual cycle, called the estrus cycle in most species and menstrual cycle in women. Menstrual cycle disorders affect up to 35% of women worldwide and are a leading cause of anovulation and infertility^1^. Despite their prevalence, underlying mechanisms remain poorly defined. Current evidence implicates a convergence of genetic susceptibility, environmental and metabolic stressors, and socio-economic disparities, factors which increasing incidence in post-industrial societies^1^. Understanding the molecular basis of these disorders is essential for a better understanding of reproductive biology and for improving women’s health.

The female sexual cycle coordinates multi-organ interactions through notably neuroendocrine feedback to ensure fertility. It is governed by two pituitary hormones, luteinising hormone (LH) and follicle-stimulating hormone (FSH), secreted by gonadotrope cells under hypothalamic gonadotropin-releasing hormone (GnRH) control^2^. Gonadotropes display marked plasticity across the reproductive cycle, culminating in the preovulatory LH and FSH surge that triggers ovulation^3^. Although early studies reported transcriptional induction of the two specific subunit gonadotropin genes (*Lhb* and *Fshb*) during this surge^4^, later analyses showed that many *in vitro* identified regulators fail to explain this behaviour *in vivo*^5–7^. The pituitary’s structural heterogeneity and the scarcity of gonadotropes have further impeded the elucidation of gonadotrope regulatory mechanisms.

Dynamic biological processes rely on temporally coordinated gene modules governed by transcription factors (TFs) and distal regulatory elements such as enhancers. Together, these interactions form gene regulatory networks (GRNs) that determine cell-type identity and state. Single-nucleus multiomic approaches now enable simultaneous measurement of chromatin accessibility and transcription, allowing *in vivo* reconstruction of GRNs in complex tissues^8^. We applied this integrative strategy to map the gonadotrope gene regulatory mechanisms driving the preovulatory surge *in vivo*.

Our study reveals an unexpected cyclic epigenetic and transcriptional plasticity in gonadotrope cells, culminating during the gonadotropin surge. This plasticity supports a reorganisation of the secretory machinery and is mediated by a structured, dynamic GRN switch, controlled by previously unrecognised transcriptional regulators. Contrary to the prevailing model, *Lhb* is not transcriptionally upregulated during the LH surge and, accordingly, is not a major GRN target. Instead, *Fshb* is regulated through a complex interplay between multiple newly identified enhancers and key GRN TFs, defining a new framework for gonadotrope gene regulation across the estrus cycle. These findings provide the first *in vivo* characterisation of the regulatory networks controlling the preovulatory surge in rat gonadotropes and uncover new pathways potentially essential for human fertility.

## Results

### Hormonal Dynamics during the Estrus Cycle in Rats

In rats, the female reproductive cycle spans four days and includes Diestrus 1 and 2 (D1 and D2), Proestrus (PE), and Estrus (E). Under our breeding conditions, 95% of Wistar female rats exhibited a regular estrus cycle, as determined by cytological analysis of vaginal smears (Supplementary Figure S1).

#### Pituitary hormones

LH displayed a clear preovulatory surge, peaking one hour before the light-dark transition (at PE 6 pm), consistent with a unimodal pattern (ANOVA, p = 0.035; Figure 1A). In contrast, FSH exhibited a trend toward bimodality (p = 0.08), with two peaks estimated at PE 11 pm and E 4 am (Figure 1B), in agreement with previous reports^9^. This bimodal profile suggests that distinct regulatory mechanisms govern FSH synthesis and secretion during the PE-E night. Notably, no FSH peak coincided with the LH surge, although FSH levels were already significantly elevated at this time relative to PE morning (2.8-fold increase, p = 0.004).

**Figure 1:**
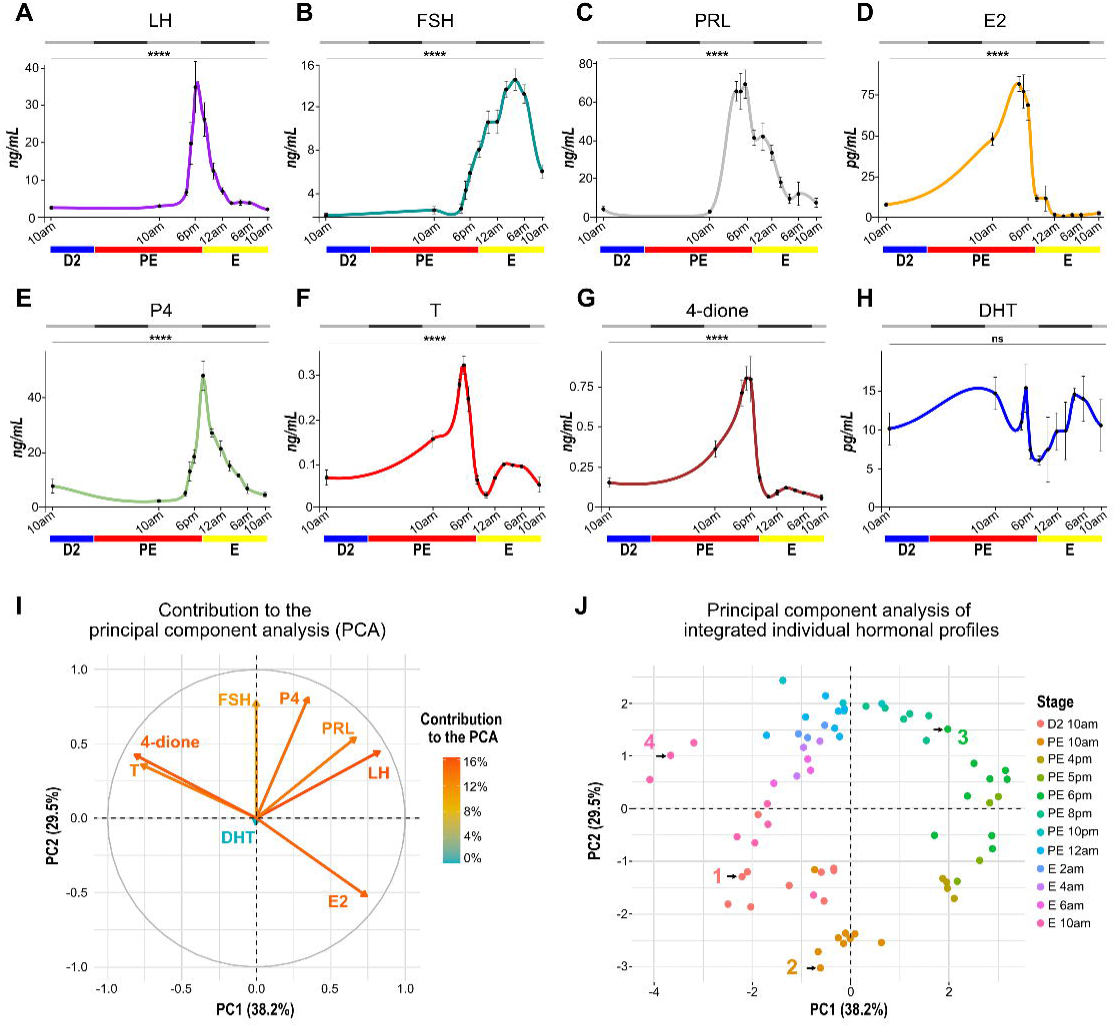
Characterisation and modelling of hormonal profiles during the estrus cycle in female rats. **(A-H)** Circulating levels of the pituitary hormones LH (**A**), FSH (**B**) and PRL (**C**), as well as the sex steroids E2 (**D**), P4 (**E**), T (**F**), 4-dione (**G**) and DHT (**H**). Results are presented as means ± SEM, and a fitted curve was generated using local polynomial regression (LOESS). Analyses were conducted on 69 individuals, with at least three per time point. Sampling times were: D2 10 am; PE 10 am, 4 pm, 5 pm, 6 pm, 8 pm, 10 pm, 12 am; and E 2 am, 4 am, 6 am, 10 am. Significant variation throughout the estrus cycle was confirmed using a non-parametric Kruskal-Wallis test followed by Dunn post-hoc tests with Benjamini-Hochberg correction applied to the global p-values. **** p < 0.0001. **(I)** Relative contribution of each variable (hormone levels) to the principal component analysis (PCA); vector colour and length indicate the magnitude of each variable contribution to the total variance. PC1 and PC2 together explain approximately 67% of the biological variance. **(J)** Coordinates of each individual on PC1 and PC2; colours indicate the sampling stage, and individuals used for snMultiome analysis are indicated by a black arrow and numbered.

In addition, we detected a prolactin (PRL) surge during the PE afternoon (Figure 1C), as previously reported in rats^10^. However, this surge began at least two hours before the LH surge, with an onset around PE 4 pm, consistent with recent findings in mice^11^.

#### Sex steroid hormones

Estradiol (E2) levels increased steadily from D2 and reached a sharp peak at PE 6 pm before rapidly declining (Figure 1D). In contrast, progesterone (P4) rose later, reaching its maximum at PE 8 pm (Figure 1E), consistent with previous studies^10,12^.

Among androgens, both androstenedione (4-dione) and testosterone (T) exhibited a pronounced PE afternoon surge (5-6 pm), paralleling E2 dynamics (Figure 1F-G). A second, smaller peak was detected at E 2-4 am, without a corresponding increase in E2. By contrast, dihydrotestosterone (DHT) levels remained low and stable throughout the cycle (Figure 1H). These findings not only confirm previous reports^12^ but also reveal a previously unrecognised second androgen peak during the PE-E night, temporally uncoupled from E2, suggesting distinct regulatory mechanisms of androgen and estrogen production.

#### Integrated hormonal profiles

We performed a principal component analysis (PCA) on all endocrine profiles. LH, FSH, E2, P4, 4-dione, and T emerged as the main contributors to estrus cyclicity, whereas DHT had negligible influence (Figure 1I). PC1 (38.2% of variance) was dominated by LH, PRL, E2, T and 4-dione, capturing proximity to the preovulatory surge, while PC2 (20% of variance) was driven by FSH and P4, likely reflecting the secondary FSH surge (Figure 1I).

Projection of individuals on the PC1/PC2 plane revealed a continuous circular trajectory, faithfully recapitulating the cyclical nature of the estrus cycle (Figure 1J). Guided by their positions, we selected four representative animals for further single nucleus Multiome (snMultiome) profiling: rat n°1 (negative PC1/PC2; corresponding to D2), rat n°2 (neutral PC1, negative PC2; corresponding to PE morning), rat n°3 (positive PC1/PC2; corresponding to late PE), and rat n°4 (negative PC1, positive PC2; corresponding to E phase).

### Single-Cell Multiome Characterisation of Gene Expression and Chromatin Accessibility in Pituitary Cell Populations

Nuclei purified from frozen hemi-pituitaries collected in the four above-mentioned females were processed using the Single Cell Multiome ATAC + Gene Expression platform (10x Genomics), followed by high-throughput sequencing. Transcriptomic and epigenomic analyses were conducted using the latest *Rattus norvegicus g*enome assembly mRatBN7.2 (rn7).

#### Re-annotation of the rat genome

Single-cell RNA-seq quantification relies heavily on accurate annotation of 3′ untranslated regions (3′UTRs), especially with 10x Genomics chemistry. However, existing rat RefSeq annotations often contain truncated or missing 3′UTRs in pituitary-expressed genes, leading to underestimated transcript counts^13,14^. We observed this quantification bias, as exemplified in Figure 2A for two representative key pituitary genes.

**Figure 2:**
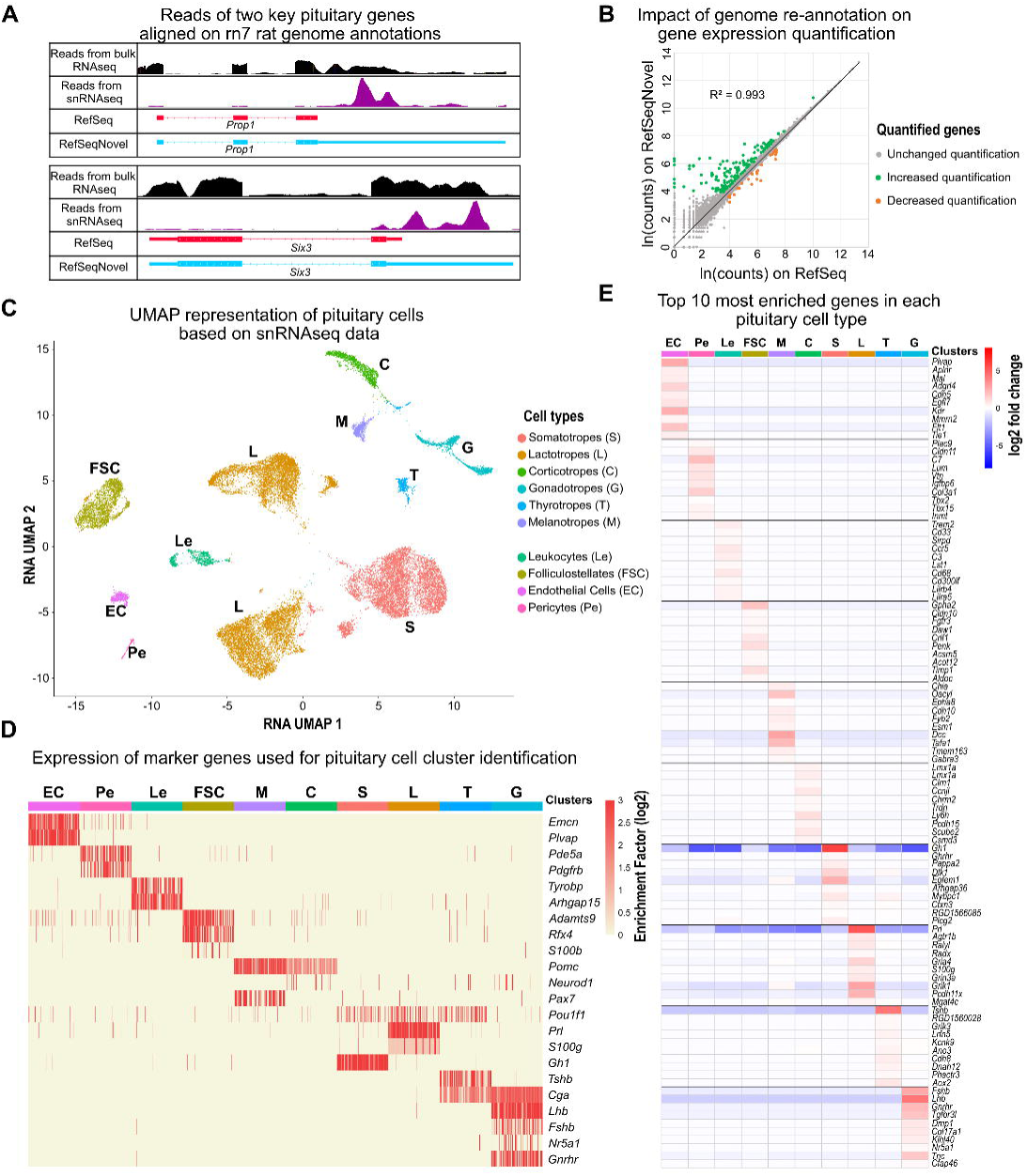
snRNAseq analysis of female rat pituitary glands. **(A)** Visualisation of mRNA alignment on the rat genome, derived from published bulk RNAseq data (black) or from our snRNAseq data (purple), for the genes *Prop1* (top panel) and *Six3* (bottom panel). Bulk RNAseq data originate from a collection of pituitary glands from male and female rats from neonatal to pubertal stages. The rat reference genome available in public databases (RefSeq) is shown in red, and the refined annotation of the rat genome (RefSeqNovel) is represented in blue. **(B)** Linear regression of log-transformed gene expression aligned on both RefSeq and RefSeqNovel. Genes with increased expression quantification are indicated in green, whereas genes with decreased quantification are indicated in orange. Genes with unchanged quantification are indicated in grey. **(C)** UMAP embedding of pituitary cell types based on snRNAseq data. Cells are coloured according to their identity. **(D)** Expression of specific markers used for cell type classification across a random sample of 100 cells per pituitary cell type. Values are represented as the enrichment factor (Log2) compared with all other cell types. **(E)** Heatmap of the top 10 most enriched genes in each pituitary cell type. Expression enrichment of each gene is represented as the Log2 fold change compared with all other pituitary cell types.

To overcome this limitation, we re-annotated the rn7 genome using bulk RNA-seq data from male and female rat pituitaries (accession numbers in Supplementary Table 1). Tissue-specific 3′UTRs were detected with Derfinder^15^ and incorporated into an updated reference, hereafter termed RefSeqNovel. The scRNA-seq and scATAC-seq libraries were then processed with 10x Genomics CellRanger-Arc against both RefSeq and RefSeqNovel, enabling a direct comparison of gene quantification performance.

Overall, re-annotation did not substantially alter global gene expression profiles (linear regression of log-transformed values: slope = 0.99, rZ = 0.993; Figure 2B). However, RefSeqNovel markedly improved transcript quantification for 148 genes that were poorly detected in the original annotation (< 50 reads), with expression levels increasing from to 1.5 to 38.5-fold. Importantly, these included critical pituitary regulators such as *Prop1* and *Six3*. Only 31 weakly expressed genes exhibited a moderate decrease in quantification (1.3 to 3-fold).

#### Single-nucleus RNA-seq pituitary cell clustering

SnRNA-seq reads from each hemi-pituitary were aligned to the RefSeqNovel genome and processed with Seurat V5^16^. After normalisation, scaling, and integration, dimensionality reduction by PCA and UMAP identified 15 clusters (Figure 2C).

Cell-type identities were assigned based on canonical marker expression (Figure 2D, Supplementary Figure S2A-B). Strikingly, lactotropes segregated into three disconnected clusters, while gonadotropes split into two closely related clusters (Figure 2C). This contrasts with previous scRNA-seq studies, which reported a single-cluster organisation of these two lineages in male or randomly cycling female pituitaries of both mice^14^ and rats^13^. Our findings therefore strongly suggest that lactotropes and gonadotropes exhibit marked transcriptional plasticity across the estrus cycle, whereas the other cell types, each represented by a single cluster, likely undergo more limited transcriptional remodelling. The robustness of this clustering pattern is supported by the absence of batch effects, as all other pituitary lineages displayed unambiguous single clusters (Supplementary Figure S2A).

We next identified cluster-specific marker genes (Figure 2E, Supplementary Table 2). Approximately 45% of the cell-specific genes reported by Fletcher *et al.*^13^ were validated, while we additionally uncovered novel lineage-predominant markers. Notably, *Tnc* and *Col17a1* were found to be restricted to gonadotropes, with expression further confined to specific estrus stages (Supplementary Table 2).

#### Single-nucleus ATAC-seq pituitary cell clustering

Chromatin accessibility was analysed using the Signac package^17^. Dimensionality reduction of integrated snATAC-seq data revealed 12 clusters (Figure 3A). Overall, chromatin activity was strongly correlated with transcript levels (rZ = 0.66, p < 0.001), and further confirmed cluster identity (Figure 3B, Supplementary Figure S2C-D), indicating that epigenomic data reliably captured transcriptional output.

**Figure 3:**
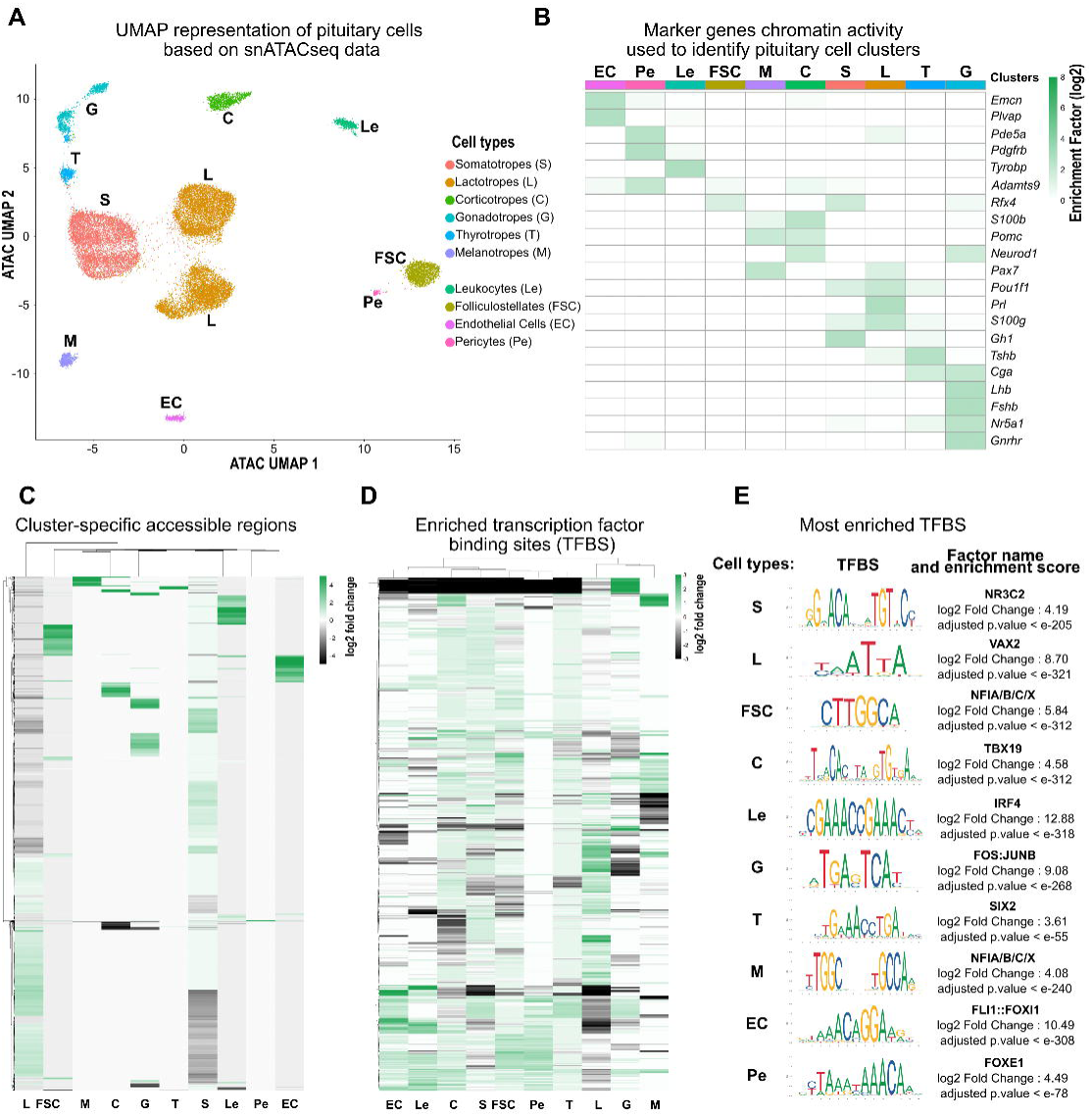
snATACseq analysis of female rat pituitary glands. **(A)** UMAP embedding of pituitary cell types based on snATACseq data. Cells are coloured according to their identity. **(B)** Chromatin activity of the specific marker genes used to classify each pituitary cell type. Chromatin activity is calculated as an estimate of chromatin accessibility around each gene locus, integrating accessibility across the gene body and promoter regions to approximate transcriptional activity (as implemented in the Signac package). Values are represented as the enrichment factor (Log2) compared with all other cell types. **(C)** Heatmap of cell type-specific accessible chromatin regions in the female pituitary gland. The colour scale represents the Log2 fold change of accessibility compared with all other cell types. **(D)** Heatmap of all enriched TFBS in cell type-specific accessible chromatin regions in female pituitary gland. The colour scale represents the Log2 fold change of accessibility compared with all other cell types. **(E)** Most enriched TFBS in each pituitary cell type. TFBS motifs are represented as indicated in the JASPAR database. The Log2 fold change of TFBS enrichment and the associated statistical significance are also indicated.

Most pituitary lineages segregated into single clusters based on their epigenomic profiles. However, both gonadotropes and lactotropes resolved into two distinct clusters, in agreement with snRNA-seq results, suggesting that these two cell types undergo marked epigenomic plasticity across the estrus cycle.

We next identified cluster-specific accessible chromatin regions (Figure 3C) and performed transcription factor binding site (TFBS) enrichment analyses which revealed lineage-specific regulatory signatures (Figure 3D-E, Supplementary Figure S2E). For instance, corticotropes showed a strong enrichment for TBX motifs, including binding site for TBX19, a master regulator of corticotrope fate and function^18^, while gonadotropes displayed enrichment for AP-1 composite motifs (FOS:JUNB)^19^ and nuclear receptor C4 zinc finger motifs, notably NR5A1^20^, both key regulators of gonadotrope identity and activity. Additionally, leukocytes were enriched for immune-related motifs, including IRF4^21^, reflecting their lineage-specific regulatory programs. Remarkably, gonadotropes are the only cells that display an exclusive cell-specific cluster of TFBS (Figure 3D), suggesting the existence of a distinct epigenetic program that is dynamically activated during the estrus cycle.

#### Integrated pituitary cell clustering

snRNA-seq and snATAC-seq datasets were integrated using multimodal dimensional reduction (Figure 4A). This integration revealed cell-type-specific associations between gene expression and local chromatin accessibility. Chromatin-to-gene linkage analysis identified regulatory regions whose accessibility correlated with the expression of proximal genes. Restricting the analysis to cell-type-specific gene-regulatory element pairs within the same topologically associating domain (TAD)^22^ revealed hundreds of putative *cis*-regulatory elements driving lineage-specific transcriptional programs (Figure 4B, Supplementary Table 3).

**Figure 4:**
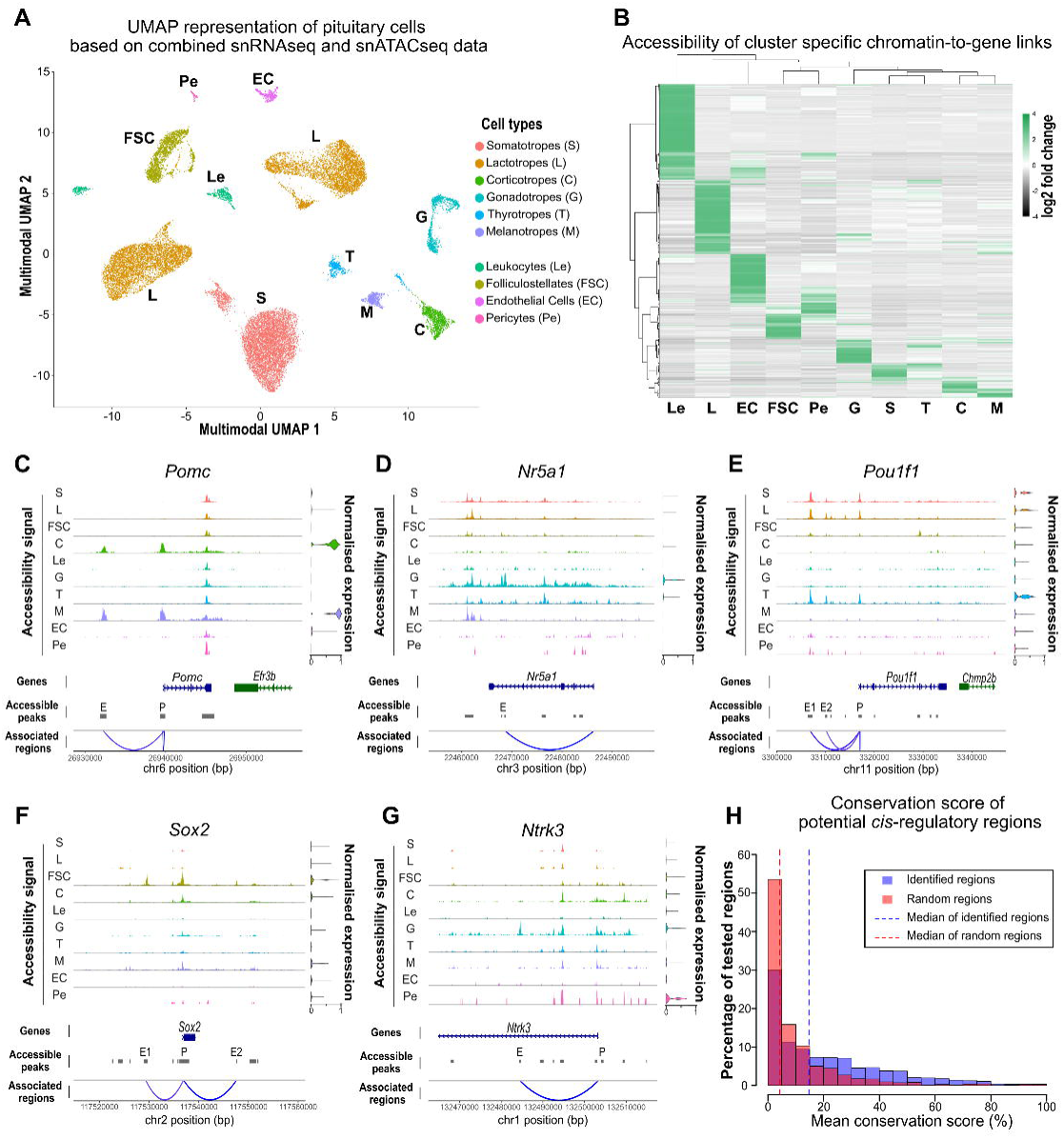
Integrated analysis of snMultiome data from female rat pituitary glands. **(A)** UMAP embedding of pituitary cell types based on combined snRNAseq and snATACseq data. Cells are coloured according to their identity. **(B)** Heatmap of cell type-specific accessible chromatin regions found to be associated with cell type-specific genes. The colour scale represents the Log2 fold change of accessibility compared with all other cell types. **(C-G)** Coverage plots of *Pomc* (**C**), *Nr5a1* (**D**), *Pou1f1* (**E**), *Sox2* (**F**) and *Ntrk3* (**G**) expression and associated chromatin accessibility across each pituitary cell type. Genomic regions are shown at the bottom of each plot, while the chromatin accessibility of these regions in each cell type is displayed at the top. Genes of interest are shown in blue, neighbouring genes in green. Accessible peaks are indicated by grey boxes, and their statistical association as potential regulatory regions of the studied gene is represented by blue links in the bottom panel. Each associated peak is labelled either as promoter (P) or as potential enhancer (E). Genome coordinates (bp) are from the rn7 assembly of the rat genome. Gene expression level in each cell type is displayed on the right of each corresponding plot. (**H**) Distribution of percentage of genomic regions according to conservation scores across 100 vertebrate species. Potential *cis*-regulatory regions are represented in blue and randomly chosen genomic regions with similar characteristics in red.

This approach successfully recovered several well-characterised pituitary enhancers, including the distal *Pomc* enhancer in corticotropes/melanotropes^23^, the late *Nr5a1* enhancer in gonadotropes^24^, and both early and late *Pou1f1* enhancers in thyrotropes, somatotropes and lactotropes^25^ (Figure 4C-E). Of note, the early *Nr5a1* enhancer, previously shown to initiate expression during embryogenesis^26^, is also active in adult female gonadotropes. Remarkably, this same enhancer was also primed in thyrotropes, raising the possibility that thyrotropes may retain a latent plasticity toward gonadotrope identity in adulthood, an idea consistent with recent report of thyrotrope-to-gonadotrope transdifferentiation in juvenile mice^27^.

Beyond these known enhancers, our analysis identified novel candidate regulatory regions for several key pituitary genes (Figure 4F-G), including, for instance, *Sox2* in folliculostellate cells^28^ and *Ntrk3* in gonadotropes^29^.

Importantly, identified potential *cis*-regulatory regions exhibited a higher evolutionary conservation than random genomic regions across 100 vertebrate species including Human (3.75 -fold increase of mean conservation, p < 0.001; Figure 4H), underscoring their putative functional importance. Conservation metrics and corresponding human orthologous regions are provided in Supplementary Table 4.

Collectively, these results highlight the critical role of conserved *cis*-regulatory elements in shaping pituitary cell identity and function, making them good candidates for investigating the genetic basis of idiopathic human hypopituitarism.

### Molecular Plasticity of Female Pituitary Cells During the Estrus Cycle

#### Differential gene expression analysis reveals cell-type-specific transcriptomic plasticity

Pairwise differential expression analysis across successive cycle stages (D2 10 am vs. PE 10 am, PE 10 am vs. PE 6 pm, PE 6 pm vs. E 10 am and E 10 am vs. D2 10 am) revealed a marked transcriptomic remodelling in lactotropes, gonadotropes, and corticotropes, with 389, 260, and 267 differentially expressed genes (DEGs), respectively (fold-change ≥ 2, adjusted p-value < 10^-5^; Figure 5A). By contrast, all other cell types displayed fewer than 50 DEGs, suggesting substantially lower responsiveness to cyclic hormonal fluctuations.

**Figure 5:**
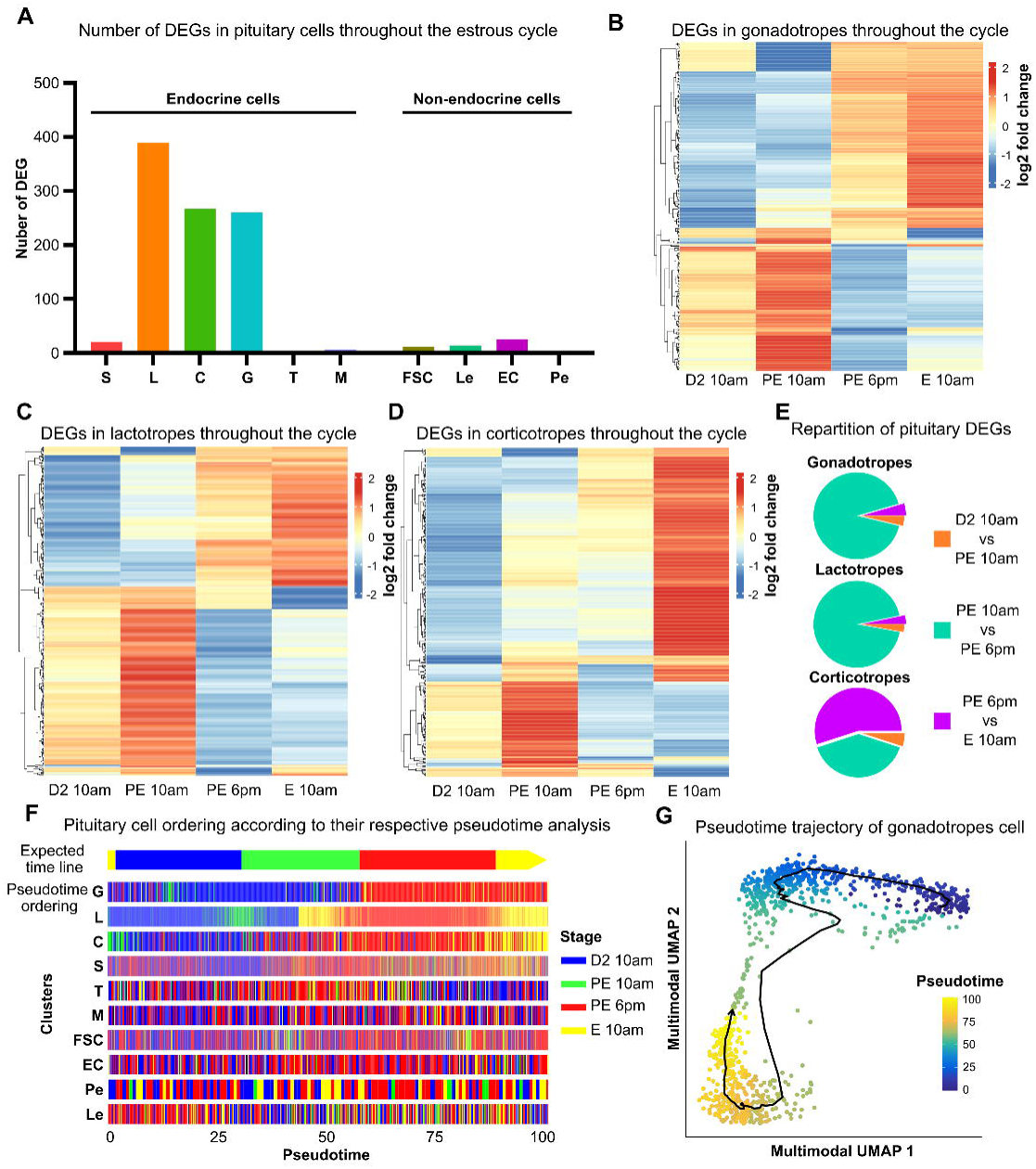
Transcriptional plasticity of pituitary cells during the estrus cycle. **(A)** Total number of DEGs in each pituitary cell type between all successive stages of the estrus cycle. Differential expression was considered significant for genes showing at least a twofold change between two successive stages with an adjusted p-value lower than 10^-^⁵. **(B-D)** Heatmap of DEG expression levels in gonadotrope (B), lactotrope (C) and corticotrope (D) cells at the four stages of the estrus cycle. The colour scale represents the Log2 fold change of gene expression relative to the average expression in the other stages. **(E)** Proportion of DEGs varying significantly between two successive stages of the cycle in gonadotrope, lactotrope and corticotrope cells. **(F)** Ordering of pituitary cells according to pseudotime analysis. The top panel shows the expected distribution according to the stage of sample collection, with colours corresponding to sampling stages. Pseudotime infers the ordering of cells according to their respective snMultiome molecular profiles. Pseudotime orderings for every pituitary cell types are illustrated; each vertical line corresponds to an individual cell which is coloured according to its sampling stage. **(G)** Multimodal UMAP embedding of gonadotrope cells and their pseudotime trajectory. The trajectory illustrates the temporal dynamics of the gonadotrope transcriptome within the UMAP space. Each point corresponds to an individual cell, coloured according to its progression along the trajectory from the initial state (dark blue, pseudotime = 0) to the most advanced state (yellow, pseudotime = 100).

These results were validated by RT-qPCR profiling of 40 cluster-specific candidate genes, selected for high cell-type specificity (> 3-fold higher expression than average across clusters), strong differential regulation (fold-change ≥ 2), and stringent significance (adjusted p-value < 10 ^-5^) (Supplementary Figure S3.1-3). Temporal dynamics revealed distinct windows of high plasticity across cell types. In lactotropes and gonadotropes, the vast majority of transcriptional changes (94% and 92% of DEGs, respectively) occurred between PE morning and evening, coinciding with the preovulatory surge (Figure 5B-C, E). In contrast, corticotropes exhibited the highest changes between PE evening and early estrus (55% of DEGs; Figure 5D-E).

To further investigate gonadotrope plasticity, we compared our rat dataset with the mouse dataset of Qiao *et al.*^30^, who performed bulk RNA-seq on FACS-isolated gonadotropes (D2 vs. PE 6 pm). Among the 547 DEGs detected in rat gonadotropes (D2 vs. PE 6 pm; fold-change ≥ 2, adjusted p-value < 10^-^⁵; Supplementary Figure S4A), only eight were shared with the mouse dataset (Supplementary Table 5). This poor concordance may reflect methodological differences: unlike Qiao *et al.*, who enzymatically dissociated cells at 37 °C followed by FACS sorting, procedures known to alter transcriptomic signatures^31^, we snap-froze pituitaries prior to nuclear isolation, thereby better preserving the *in vivo* transcriptional state.

#### Pseudotime analysis uncovers coordinated transcriptional regulation in gonadotropes

Because the estrus cycle is inherently dynamic, stepwise pairwise comparisons may inadequately capture the kinetics of the transcriptional regulatory program within the pituitary, throughout the cycle. To overcome this, we reconstructed cell type-specific trajectories using pseudotime analysis (Figure 5F). In gonadotrope and lactotrope, and to a lesser extent corticotrope cells, pseudotime ordering recapitulated the expected biological progression, as reflected by the proportion of pseudotime variance explained by original cell origin (ηZ_Gonadotrope = 0.56, ηZ_Lactotrope = 0.75, ηZ_Corticotrope = 0.58). By contrast, the pseudotime trajectories of other pituitary lineages appeared mostly random (ηZ_Somatotrope = 0.18, ηZ_Melanotrope = 0.04, ηZ_Thyrotrope = 0.02, ηZ_Folliculostellate = 0.07, ηZ_Endothelial = 0.03, ηZ_Pericyte = 0.04, ηZ_Leukocyte = 0.05).

Taken together, these results underscore that only gonadotrope, lactotrope, and corticotrope populations exhibit coordinated transcriptional reprogramming across the estrus cycle. Given their central role in regulating the reproductive function, we focused subsequent analyses on gonadotrope cells.

Gonadotrope cells and their inferred pseudotime ordering were projected onto the UMAP plan (Figure 5G). The trajectory followed a single, continuous path without branching, connecting the two gonadotrope subclusters. This pattern indicates a progressive, unidirectional shift in cellular states across the estrus cycle, consistent with gradual and coordinated transcriptional reprogramming in response to physiological cues. Pseudotime projections for the other pituitary cell types are provided in Supplementary Figure S4B.

#### Gonadotrope gene regulation and expression during the estrus cycle

To capture dynamic regulation in gonadotrope cells, we integrated chromatin accessibility, gene expression, and gene velocity across the estrus cycle. Gene velocity, defined as the balance between unspliced and spliced mRNA^32^, provides an additional measure of transcriptional dynamics. We inferred chromatin accessibility and gene expression changes along pseudotime, whereas velocity was inferred along latent time^32^. Because pseudotime and latent time order cells differently, we established a unified temporal axis, hereafter referred as “regulatory-time”, by combining pseudotime (azimuthal angle θ) and latent time (polar angle ϕ) into a cyclic model. Pseudotime and latent-time ordering were found highly correlated (RZ = 0.85), and the integrated regulatory-time framework explained more variance in cell ordering than either pseudotime and latent-time alone (ηZ_regulatory-time = 0.67 vs. 0.57 for pseudotime and 0.60 for latent-time) (Figure 6A-B).

**Figure 6:**
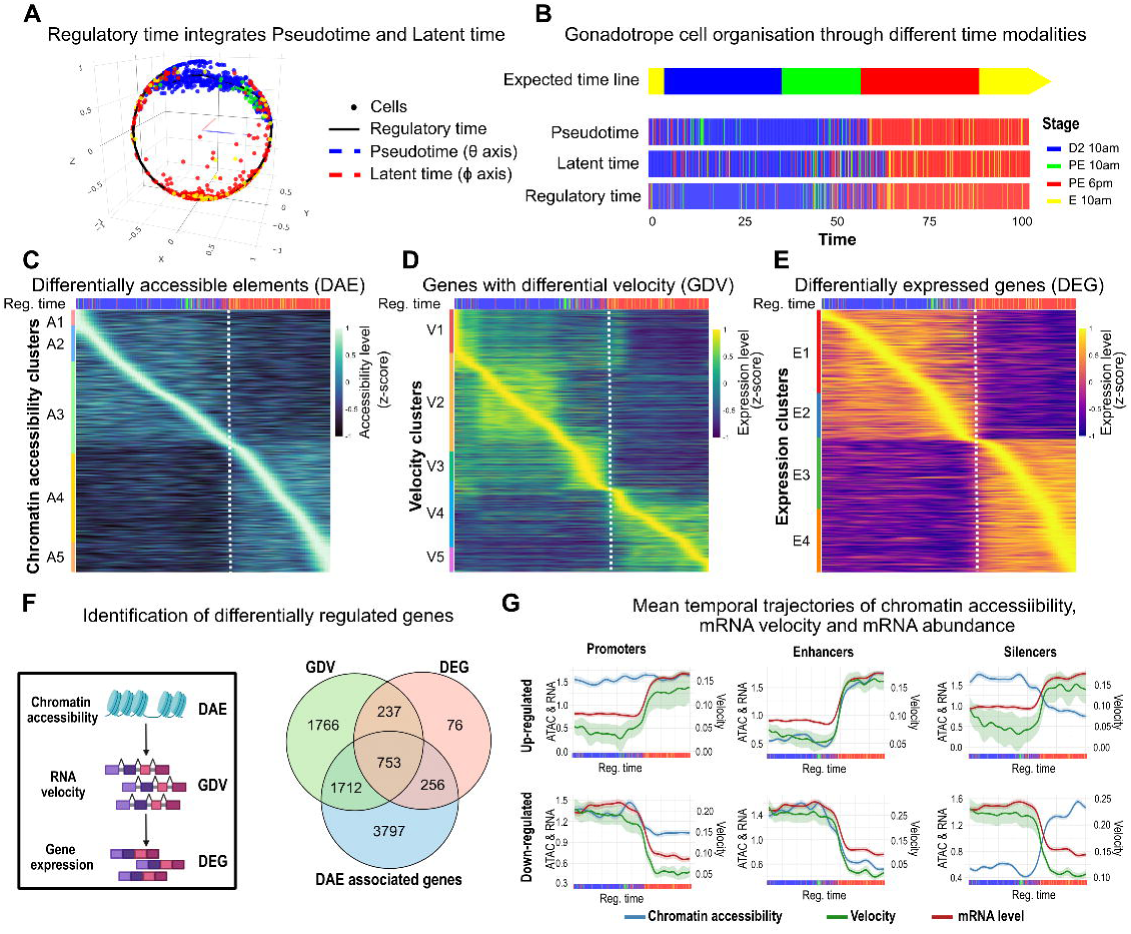
Molecular plasticity of pituitary gonadotrope cells during the estrus cycle. **(A)** Spherical representation of gonadotrope cell organisation, with pseudotime (θ axis, blue dashed line) and latent time (ϕ axis, red dashed line). Each gonadotrope cell’s pseudotime and latent time values were projected onto a circular space, and smoothed models were used to combine them into a single trajectory (solid black line). Individual cells are shown as coloured points by sampling stage: D2 10 am (blue), PE 10 am (green), PE 6 pm (red), and E 10 am (yellow). The projection of cells onto this periodic model defines a new time value, the “regulatory time,” capturing the cyclical progression of cellular states across the estrus cycle. **(B)** Ordering of pituitary gonadotrope cells according to pseudotime, latent time or regulatory time, with colours corresponding to the sampling stage. The upper panel shows the expected distribution according to the stage of sample collection. **(C-E)** Heatmaps of DAEs (**C**), GDVs (**D**) and DEGs (**E**) in gonadotrope cells across regulatory time. The upper panel shows gonadotrope cell distribution according to regulatory time. Signals are shown for each region or gene (lines) across the regulatory time (columns). The heatmap colour scale represents the relative signal (Z-score) in gonadotrope cells across the regulatory time. DAEs, GDVs and DEGs are clustered based on their respective accessibility (Cluster A1 to A5), velocity (Cluster V1 to V5) or expression profiles (Cluster E1 to E4). A dotted line indicates the transition between early to late PE, framing the preovulatory surge. **(F)** Identification of differentially regulated genes in gonadotrope cells across the estrus cycle. A gene was considered differentially regulated if associated with a DAE and displaying differential velocity and expression across regulatory time. The Venn diagram represents the overlap between GDV, DEG and DAE-associated genes. **(G)** Mean temporal trajectories of chromatin accessibility, mRNA velocity and mRNA abundance for each identified pair of gene and DAE (n = 1816 pairs). Mean ± SEM signals are plotted across the regulatory time. ATAC and RNA data are rescaled to a common amplitude (left y-axis) for comparison, while velocity is shown on its native scale (right y-axis).

Using this framework, we identified 12,022 differentially accessible elements (DAEs) (Figure 6C), which grouped into five temporal clusters. The most pronounced chromatin remodelling event occurred during the early-to-late PE transition. RNA velocity analysis revealed 4,468 genes with differential velocity (GDVs), segregating into five clusters (Figure 6D), while expression analysis identified 1,322 DEGs across four clusters (Figure 6E). Notably, GDV and DEG profiles across regulatory time highlighted the early-to-late PE transition as the stage of maximal remodelling, concordant with chromatin accessibility changes. This concordance across three regulatory layers underscores a coordinated epigenetic and transcriptional reprogramming, establishing the early-to-late PE transition as a critical regulatory point.

We integrated DAEs with genes located within TADs and identified 6,518 genes associated with at least one DAE. Approximately 40% of these (n = 1,712 + 753) also showed significant variation in RNA velocity or expression across the estrus cycle (Figure 6F). The remaining DAEs likely correspond to primed or non-regulatory genomic regions. Notably, 75% of DEGs (n = 753 + 237) also exhibited RNA velocity changes, whereas only 22% of GDVs showed alterations in expression. This indicates that RNA velocity frequently captures early or transient regulatory activity preceding detectable shifts in steady-state mRNA levels. We therefore focused on the 753 high-confidence genes showing concordant chromatin, velocity, and expression dynamics.

For these genes, we analysed the temporal relationship between chromatin accessibility, RNA velocity, and gene expression. Positively correlated *cis*-regulatory regions were classified as promoters or potential enhancers, while negatively correlated regions were considered potential silencers. Averaged profiles revealed that promoter accessibility exhibited only subtle changes across regulatory time with weak association to transcription. In contrast, enhancer accessibility closely tracked transcriptional activation while silencer accessibility mirrored repression and, again, the early-to-late PE transition emerged as a major regulatory shift (Figure 6G).

#### Functional states of gonadotropes across the estrus cycle

We performed Gene Ontology (GO) analysis on gonadotrope DEGs and identified 4 clusters (E1 to E4) related to key stages of the estrus cycle (Figure 7A and Supplementary Table 6).

**Figure 7:**
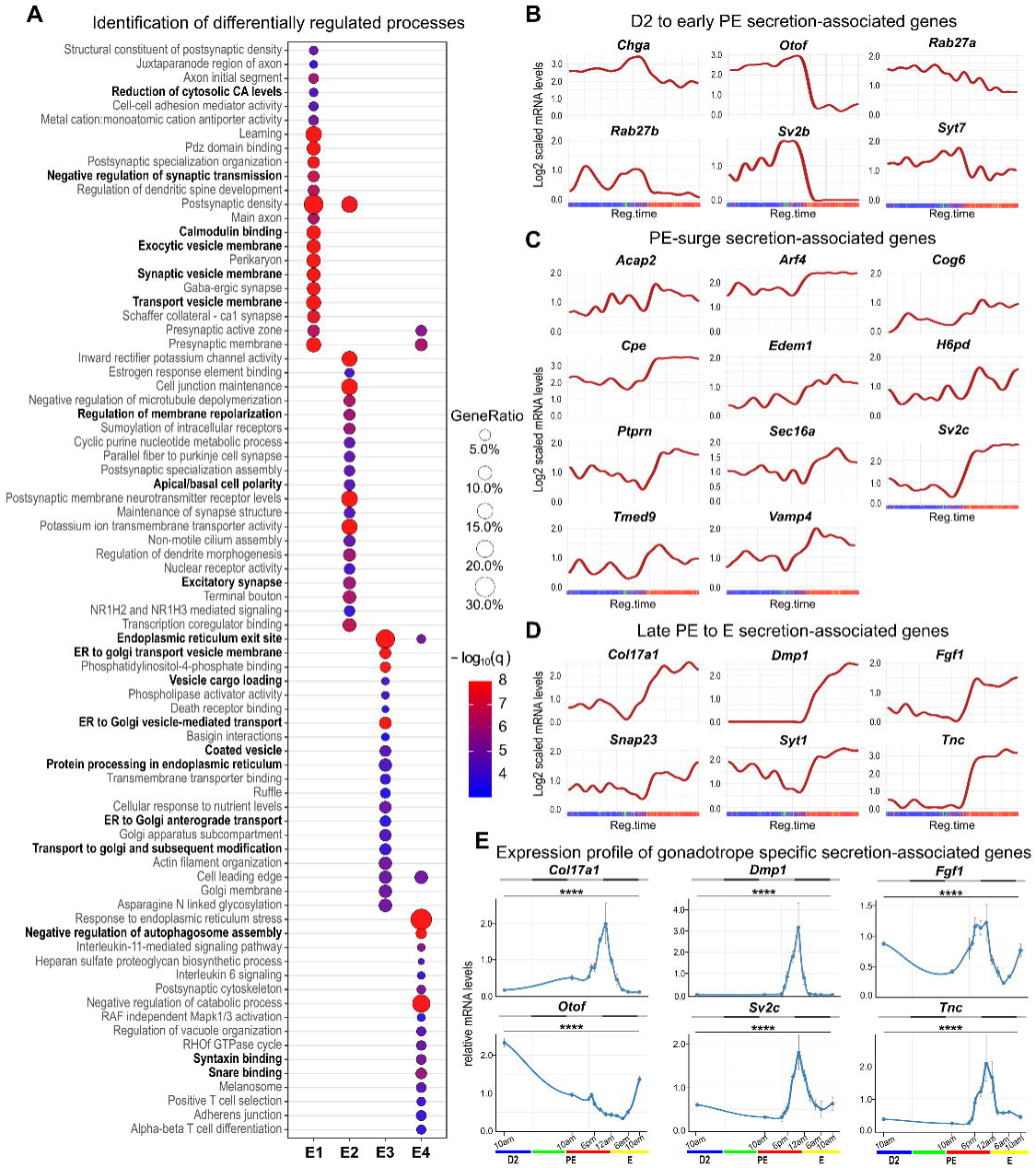
Identification of biological processes regulated in gonadotrope cells throughout the estrus cycle. **(A)** Differentially regulated biological processes in gonadotrope cells throughout the estrus cycle. The analysis highlights the top twenty processes enriched within each DEG cluster. Analyses were conducted using both rat and human ontologies across the complementary KEGG, Reactome and Gene Ontology databases. Redundant or semantically overlapping ontologies were automatically merged, and statistical enrichment was recalculated. Processes potentially related to secretory pathways are indicated in bold. Sphere size corresponds to the proportion of genes involved in a given process, and colour indicates the adjusted q-value reflecting the significance of enrichment. **(B-E)** Log2-normalised expression patterns of secretory pathway-related genes across the regulatory time. For genes expressed predominantly in gonadotropes cells, the expression pattern was also measured by RT-qPCR on hemi-pituitaries from a total of 69 individuals, with at least three per time point (**E**). Sampling times were: D2 10 am; PE 10 am, 4 pm, 5 pm, 6 pm, 8 pm, 10 pm, 12 am; and E 2 am, 4 am, 6 am, 10 am. Data are presented as relative mRNA levels normalised to two housekeeping genes, *Ppia* and *Ywhae*. Significant variation throughout the estrus cycle was confirmed using a non-parametric Kruskal-Wallis test followed by Dunn post-hoc tests with Benjamini-Hochberg correction applied to the global p-values. **** p < 0.0001.

##### D2 (Cluster E1)

Upregulated genes in gonadotrope cells are associated with cytosolic Ca^2+^ level regulation, calmodulin binding and several trafficking -related ontologies, including synaptic-related GO, which may support basal pulsatile neuronal-like LH secretion^33^.

##### Early PE (Cluster E2)

We observed enrichment of cell polarisation-related GO terms (regulation of membrane repolarisation, apical/basal cell polarity) alongside potassium channel-related GO terms (inward rectifier potassium channel activity, potassium ion transmembrane transporter activity), consistent with functional priming of gonadotropes in anticipation of the upcoming PE surge^34^.

##### PE surge (Cluster E3)

GO terms were enriched for Golgi-related and trafficking processes (*e.g.*, Endoplasmic Reticulum (ER) to Golgi transport vesicle membrane, coated vesicle, ER-to-Golgi anterograde transport), suggesting increased glycoprotein synthesis and secretion, consistent with the massive gonadotropin release during the PE surge. In parallel, enrichment of motility-related GO terms (ruffle formation, actin filament organisation, cell leading edge) may reflect reorganisation of gonadotrope cell networks during the PE-E night^35^.

##### Late PE to PE-E transition (Cluster E4)

During late PE night, gonadotrope cells upregulate genes associated with the ER stress response and negative regulation of autophagosome assembly, suggesting a previously unrecognised phase of cellular stress adaptation. Interestingly, gonadotropes also activate inflammatory signalling pathways involving IL1 and IL6, the latter previously shown to enhance gonadotropin secretion^36^ which may thus contribute to the late FSH surge.

Overall, these results highlight the dynamic functional reprogramming of gonadotrope cells across the estrus cycle, with the early-to-late PE transition representing a period of maximal cellular plasticity. This reprogramming likely coordinates electrical, secretory and structural adaptations to ensure the gonadotropin surge.

#### Stepwise maturation of the secretory machinery in gonadotropes during the estrus cycle

GO-based clustering revealed coordinated changes in cellular architecture, including enrichment of trafficking- and vesicle-related processes, indicative of dynamic modulation of secretory function. As gonadotropin release is the hallmark of gonadotrope activity, we next examined the temporal expression of secretion-related genes across the estrus cycle (Figure 7B-E, Supplementary Figure S6) and validated expression dynamics by RT-qPCR in whole pituitary when possible (Figure 7E, Supplementary Figure S5).

##### D2 to early PE phase (E1 and E2 clusters)

Upregulation of RAB27A/B expression indicate readiness for secretion^37^, while those of SYT7^38^, STX4^39^ and SV2B^40^ are consistent with a CaZ⁺-dependant lysosomal/exocytic pathway (Figure 7B & Supplementary Figure S6). High level of CHGA expression likely promotes granule biogenesis but may simultaneously impose inhibition on final granule maturation^41^, reinforced by STXBP5L upregulation (Supplementary Figure S6)^42^. Secretion at this stage appears to rely on non-canonical CaZ⁺ sensors such as OTOF^43^ and SYT7, with SV2B potentially modulating vesicle CaZ⁺ sensitivity (Figure 7B & E). Overall, these results indicate a gonadotrope cells priming with dense-core granules and an exocytosis-competent SNARE set, while vesicular fusion seems restrained.

##### PE surge (E3)

Upregulation of CPE^44^ and PTPRN^45^ may stabilise and process hormone cargo, while VAMP4^46^ and SV2C upregulations^47^ are consistent with enhanced vesicle trafficking (Figure 7C & E). In parallel, the expression of SEC16A^48^, KDELR1^49^, and TMED9^50^ suggests robust ER exit and selective cargo sorting, whereas COG6^51^ and ARF4^52^ expression increase supports Golgi maturation and vesicle export. Increased Rab GTPase activity, through enhanced RAB5A^53^, ARL5B^54^ and ACAP2^55^ expression, likely reflects increased endosomal membrane trafficking (Figure 7C & Supplementary Figure S6). Expression of genes related to protein quality control (SEL1L^56^, EDEM1^57^, PDIA5^58^, H6PD^59^) and autophagy (BECN1^60^) indicates mechanisms safeguarding protein folding and organelle homeostasis. Collectively, these features suggest a transition from immature storage vesicles to fully competent granules, alongside with expansion of the secretory machinery.

##### Late PE to E (E4)

The observed increase in SYT1 expression likely marks a switch from earlier CaZ⁺ sensors (Koh and Bellen, 2003) to mechanisms supporting precise, high-fidelity vesicle fusion. The co-expression of SNAP23^61^ and STX3^62^ (Figure 7D, Supplementary Figure S6) suggests the establishment of a t-SNARE platform for regulated, polarised secretion. Together, these features suggest the acquisition of competence for rapid, CaZ⁺-synchronous exocytosis.

In addition, gonadotropes also overexpress several secreted factors, including extracellular matrix (ECM) components, such as FGF1^63^, TNC^64^, COL17A1^65^, and DMP1^66^, implicated in tissular integrity, vascular permeability or remodelling (Figure 7D & E), consistent with a potential tissue reorganisation during PE night.

Altogether, GO analysis supports a model in which secretory capacity is progressively built through an integrated transcriptomic program: (i) granule priming and restraint, (ii) ER-Golgi reorganisation with vesicle maturation and quality control, and (iii) acquisition of rapid, polarised exocytosis. This dynamic remodelling of the secretory pathway underpins the PE surge, revealing tight coupling between organelle architecture and regulated hormone release. Moreover, the concurrent secretion of signalling molecules and ECM components points toward an unsuspected cyclic reorganisation of pituitary tissue across the estrus cycle.

### Gonadotropin beta Subunit Genes and *Gnrhr* Expression and Regulation During the Estrus Cycle

As transcriptional regulation of gonadotropin-coding genes has been suggested to play a key role during the PE surge, we leveraged chromatin accessibility, RNA velocity, and mRNA expression data to further examine the expression and regulation of *Lhb* and *Fshb*, together with that of the gonadotropin-releasing hormone receptor gene *Gnrhr*.

#### *Lhb* gene expression is not upregulated during the preovulatory surge

Seminal study from the late 1980s and early 1990 postulated that *Lhb* expression increases just before the PE surge in female rat gonadotrope cells^67,68^. Since then, a model in which *Lhb* transcription is required for the preovulatory LH surge has been the prevailing assumption.

To reassess this model, we analysed *Lhb* expression across the estrus cycle using single-cell and RT-qPCR analyses (Figure 8A). Single-cell analysis revealed a high level of nuclear *Lhb* mRNA throughout the regulatory time, with only a moderate and transient decrease during the early-to-late PE transition (Figure 8A Top panel). RT-qPCR quantification of mature cytoplasmic transcripts confirmed this decrease, which appeared more sustained (Figure 8A Bottom panel). The discrepancy between nuclear and cytoplasmic profiles likely reflects enhanced mRNA degradation during PE-E night. RNA velocity analysis indicated persistent negative values across the cycle, consistent with ongoing transcriptional repression (Figure 8A Top panel). Spliced/unspliced transcript profile further supported this conclusion as most gonadotrope cells show a deficit of newly synthesised *Lhb* mRNA associated with an accumulation of mature transcripts (Figure 8A Middle panel).

**Figure 8:**
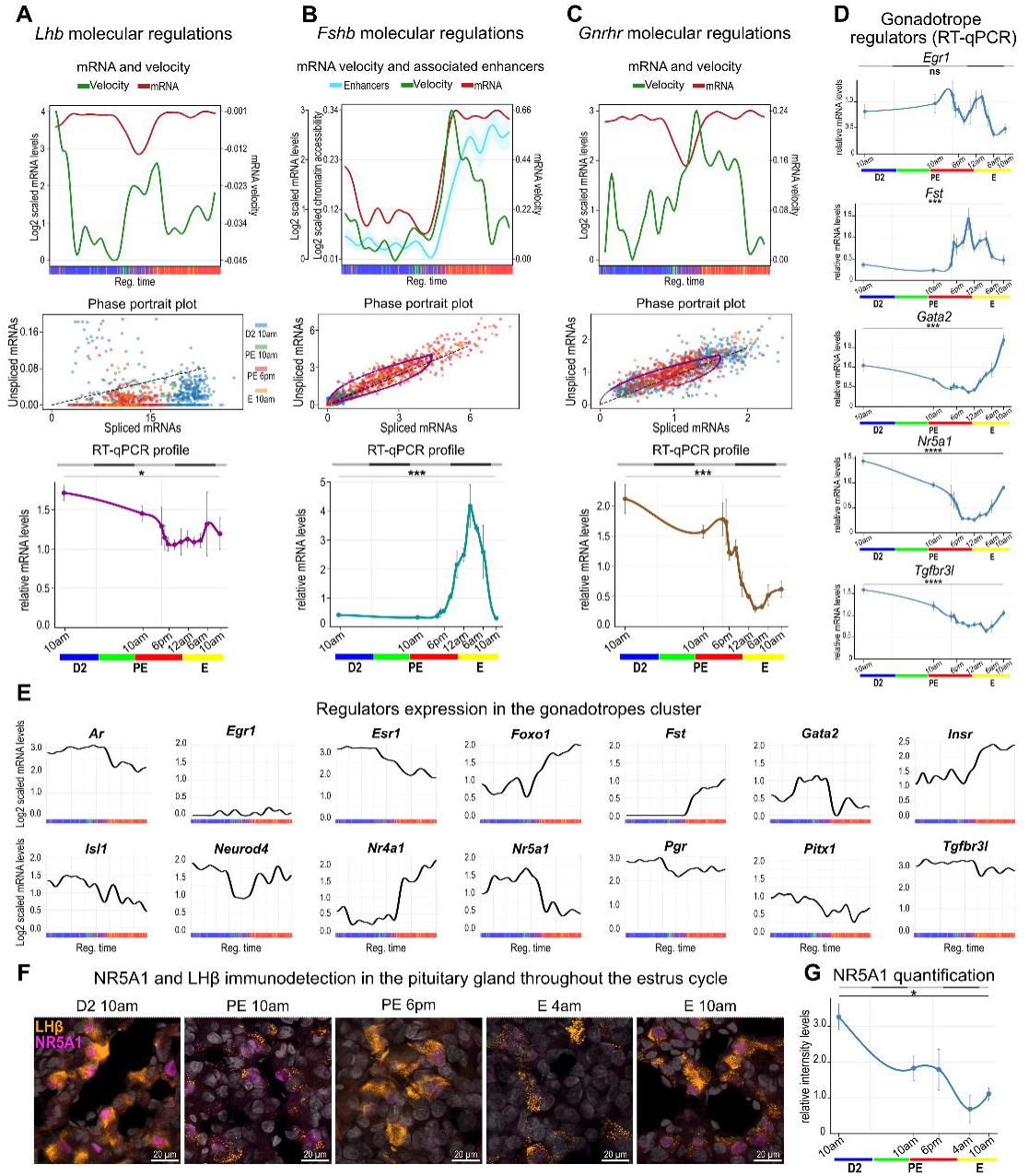
Molecular regulation of key gonadotrope genes throughout the estrus cycle. **(A-C)** Molecular regulation of *Lhb* (**A**), *Fshb* (**B**) and *Gnrhr* (**C**) in gonadotrope cells throughout the estrus cycle. Top panels show the variation of Log2-scaled mRNA levels (left y-axis), and native mRNA velocity (right y-axis) across the regulatory time. Middle panels show the gene transcriptional dynamics in all gonadotrope cells. Gonadotrope cells are coloured according to their sampling time and plotted relative to their spliced and unspliced mRNA ratio. Steady-state expression is indicated by a black dotted line and, when possible, modelling of transcriptional dynamics within the cell population is shown by a purple curve. Bottom panels show gene expression pattern measured by RT-qPCR on hemi-pituitaries from a total of 69 individuals, with at least three per time point for each stage. For the *Fshb* gene only (**B**), the variation of mean ± SEM Log_2_-scaled chromatin accessibility of all potentially associated enhancers is also shown (second left y-axis). **(D-E)** Expression patterns of some regulators of gonadotrope activity measured by RT-qPCR (69 individuals) (**D**) and by snRNAseq throughout the regulatory time (**E**). For RT-qPCR analyses, sampling times were: D2 10 am; PE 10 am, 4 pm, 5 pm, 6 pm, 8 pm, 10 pm, 12 am; and E 2 am, 4 am, 6 am, 10 am. Data are presented as relative mRNA levels normalised to two housekeeping genes, *Ppia* and *Ywhae*. Significant variation throughout the estrus cycle was confirmed using a non-parametric Kruskal-Wallis test followed by Dunn post-hoc tests with Benjamini-Hochberg correction applied to the global p-values. ns non significant, * p < 0.05, *** p < 0.001 and **** p < 0.0001. **(F-G)** Representative images (**F**) and quantification (**G**) of NR5A1 protein levels in gonadotrope cells throughout the estrus cycle. NR5A1 (magenta) and LHβ (orange) were immunolocalised in fixed pituitary sections and counterstained with DAPI (grey). NR5A1 signal in gonadotrope cell nuclei was quantified in three individuals per time point and on three images per individual. Background signal was systematically subtracted. For immunodetection analyses, sampling times were: D2 10 am; PE 10 am, 6 pm; E 4 am, 10 am. Significant variation throughout the estrus cycle was confirmed using a non-parametric Kruskal-Wallis test followed by Dunn post-hoc tests with Benjamini-Hochberg correction applied to the global p-values. * p < 0.05.

To investigate the regulatory basis of *Lhb* repression, we examined chromatin accessibility at associated *cis*-regulatory regions (Supplementary Figure S7.1A-C). The region overlapping with a recently described enhancer in mice^69^ displayed an accessibility profile highly correlated with *Lhb* expression (r = 0.78), in contrast with the proximal promoter. Together, these observations suggest that a coordinated regulation of both promoter and enhancer elements underlies the repression of *Lhb* during the PE surge.

We next analysed expression of TFs known to regulate *Lhb*, including the activators EGR1, NR5A1, and PITX1, to test whether their temporal profiles tracked *Lhb* transcription. None showed strong correlation with *Lhb* profil (r = 0.36, -0.46, and 0.10, respectively), indicating that their dynamics alone do not account for *Lhb* variation from D2 to E. Furthermore, *Egr1*, *Nr5a1*, and *Pitx1* did not display expected expression induction during PE, consistent with the observed lack of *Lhb* upregulation (Figure 8D-E). Instead, *Pitx1* and *Nr5a1* expression declined from early to late PE, and this was confirmed at the protein level for NR5A1 by immunodetection in female pituitaries (Figure 8F-G). This loss of activators likely contributes to *Lhb* repression during the surge. In parallel, the FOXO1 repressor^70^ increased during PE, potentially reinforcing this repressive state (Figure 8E).

Together, these findings indicate that high *Lhb* mRNA levels are maintained throughout the estrus cycle, ensuring sustained LH production capacity. *Lhb* transcription, initiated during or before D2, is subsequently repressed during PE through a combination of reduced enhancer and promoter accessibility and accelerated mRNA decay. Thus, the LH surge cannot be explained by *Lhb* transcriptional upregulation but must instead rely primarily on the mobilisation of pre-synthesised hormone through regulated secretory mechanisms.

#### *Fshb* is transcriptionally upregulated during the PE night

Unlike LH, whose secretion is primarily induced by GnRH pulses, FSH secretion occurs predominantly through a constitutive pathway^71^. Consequently, FSH secretion has been suggested to be tightly correlated with *Fshb* transcription^72^. In addition to GnRH, *Fshb* transcription is also regulated by activin and ovarian sex steroids including testosterone and progesterone. These additional inputs are believed to contribute to the second FSH peak occurring during the PE night (reviewed in^73^).

To further characterise *Fshb* regulation, we examined the dynamics of its expression using single-cell and RT-qPCR analyses (Figure 8B). Both approaches revealed strong transcriptional dynamics across the estrus cycle as *Fshb* transcription was found repressed during D2 and then peaked during the PE-E night. Interestingly, the subsequent decline in *Fshb* mRNA was only observed by RT-qPCR, suggesting cytoplasmic mRNA degradation at this phase. Spliced/unspliced mRNA ratios and mRNA velocity further confirmed this pattern of rapid transcriptional regulation.

To probe the regulatory basis of *Fshb* upregulation, we assessed chromatin accessibility across candidate *cis*-regulatory regions. Within the *Fshb* TAD, we identified 13 potential enhancers with significant accessibility variation during the estrus cycle (Supplementary Figure S7.2). Mean chromatin accessibility across all enhancers correlated strongly with *Fshb* expression (Figure 8B, top panel), indicative of a tight epigenetic control. These enhancers were found to be highly conserved across mammals, and most displayed gonadotrope-specific or enriched accessibility (Supplementary Figure S7.2), reinforcing their role as *Fshb* regulators. Notably, we identified a previously described *Fshb* enhancer (Enh1) whose mutation has been linked to impaired female fertility in Humans^74^.

We examined expression of previously described *Fshb* regulators. Among them, *Junb* (Figure 9F), *Fos* (Figure 9F), and *Nr4a1* (Figure 8E) showed strong positive correlations with *Fshb* (r= 0,92; 0,98 and 0,96 respectively), peaking alongside maximal RNA velocity, supporting their role in *Fshb* transcriptional activation during the PE night. *Atf2*, *Nfya/b/c*, *Smad2/3/4*, and *Foxl2* (Supplementary Figure S6) showed only weak or negative correlations, consistent with predominant post-translational regulation^75,76^. *Pgr* and *Ar* transcripts were modestly repressed (fold change <2; Figure 8E), yet their long protein half-lives (∼6-7 h)^77,78^ may explain functional contribution.

**Figure 9:**
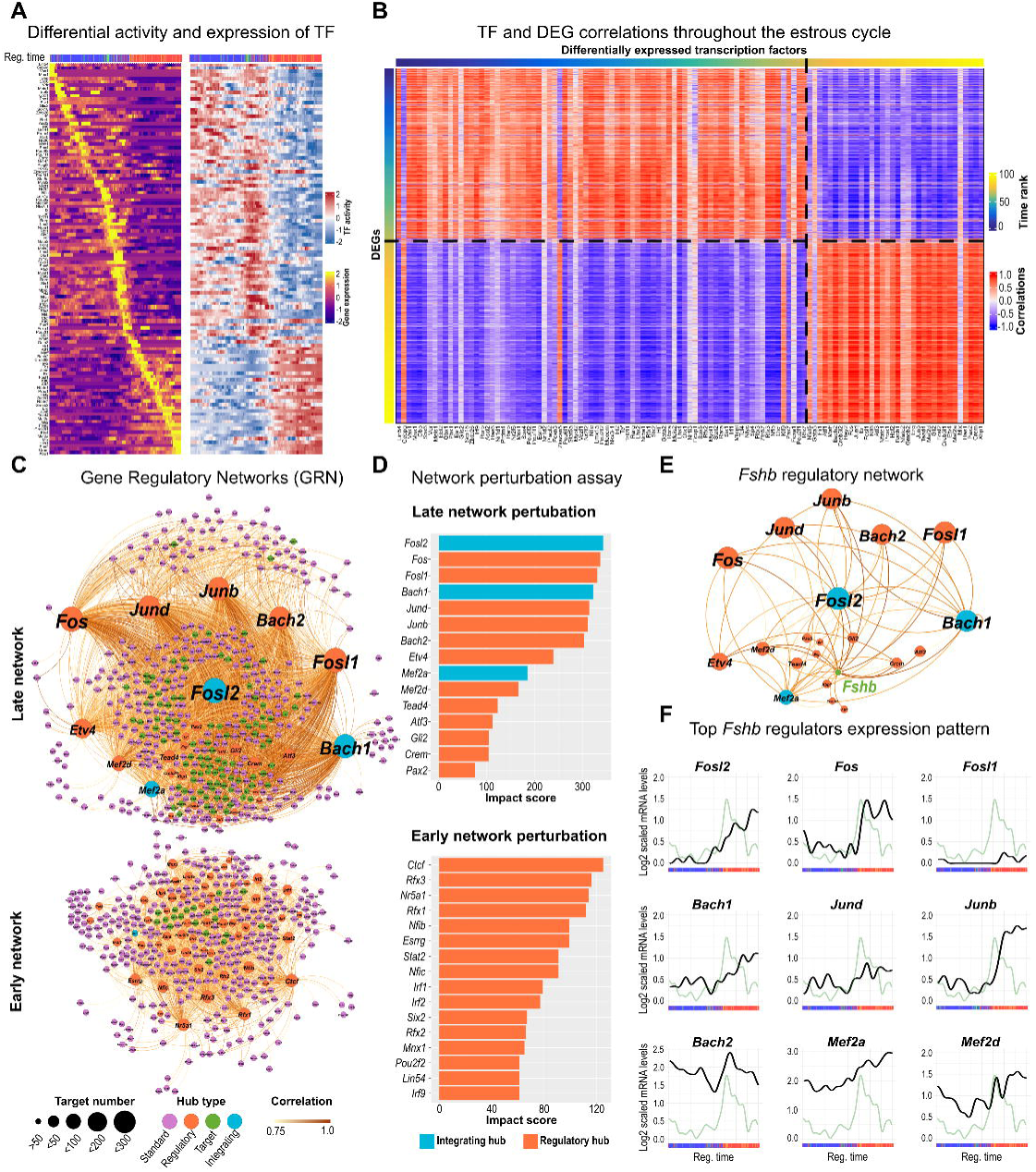
Characterisation of the gene regulatory networks mobilised in gonadotrope cells throughout the estrus cycle. **(A)** Expression and activity profiles of candidate TFs across regulatory time. RNA expression (left) and TF activity (right) are shown for each differentially expressed TF. Values are row-wise z-scores, and TFs are ordered according to their expression rank across regulatory time. **(B)** Correlation heatmap between differentially expressed TF and their differentially expressed target genes across regulatory time (columns = TFs, rows = target DEGs). Columns and rows are ordered by their respective time ranks across regulatory time. Correlation colour scale: -1 (blue) to +1 (red). **(C)** GRN reconstructed from all TFs and their target DEGs. The Early/Late classification was computed automatically from the heatmap matrix. Node size reflects the number of targets. Edge colour encodes correlation strength. Node colour denotes hub type, defined using degree enrichment: nodes with a high degree of incoming and outgoing edges were classified as “integrating”; nodes with a high degree of incoming edges as “targets”; nodes with a high degree of outgoing edges as “regulatory”; and the remaining nodes as “standard”. **(D)** Network perturbation assay. Each TF was removed *in silico* from the network and an impact score corresponding to the number of edges removed was assigned. Bars show the 15 most impactful TFs in the Late (top) and Early (bottom) networks; bar colour indicates hub type. **(E)** Restricted regulatory subnetwork focused on *Fshb*. Nodes are TFs directly connected to *Fshb* (in green). Arrows point from TFs to targets; edge colour reflects correlation strength, while node colour indicates hub type. **(F)** Regulatory time expression profiles (log2-normalised RNA) of the top *Fshb* regulators highlighted in (**E**), compared with the *Fshb* velocity pattern (in green) across regulatory time.

Unexpectedly, the FOXO1 repressor expression^70,79^ correlated positively with *Fshb* (r = 0.95; Figure 8E), peaking as *Fshb* RNA velocity declined. As FOXO1 activity can be suppressed by GnRH and insulin^80^, the peak of *Insr* expression at mid-PE (Figure 8E) may thus enable *Fshb* induction.

Interestingly, FST, that counteract activin induction of *Fshb*^9^, peaked at 10 pm between the two FSH maxima (Figure 8E). *Fst* expression positively correlated with repressors *Tgif1* (r = 0.77) and *Skil* profiles^81^ (r = 0.87; Supplementary Figure S6), indicating combined mechanisms for transient *Fshb* downregulation. Finally, decreased *Tgfbr3l* mRNA levels during PE (Figure 8D,E) may relieve the repression exerted by ovarian-derived inhibin B^82^.

Collectively, our results demonstrate that the FSH surge relies on dynamic *Fshb* transcriptional regulation. This regulation appears to involve coordinated epigenetic control at enhancers and combinatorial action of multiple TFs integrating GnRH-, activin- and insulin-mediated signalling.

#### G*nrhr* is transiently transcriptionally downregulated during the PE night

Data on *Gnrhr* expression in the female pituitary during the estrus cycle have long remained inconsistent. Early RT-PCR and autoradiography studies in the 1990s reported an upregulation of *Gnrhr* during the PE surge in rats^83^. Consistently, experiments in the LβT2 gonadotrope cell line and primary pituitary cultures showed that continuous GnRH stimulation, an experimental paradigm mimicking the PE surge, transiently increased *Gnrhr* expression^84,85^. However, later *in vivo* studies, using quantitative real-time RT-PCR, did not confirm this peak^86^.

To clarify these discrepancies, we re-examined *Gnrhr* expression in gonadotropes across the cycle using single-cell and RT-qPCR analyses (Figure 8C). Nuclear *Gnrhr* transcript levels were mostly stable during the cycle, displaying only a moderate transient decrease during the PE night. However, RT-qPCR indicated a sharp decline of cytoplasmic *Gnrhr* mRNA starting from PE 5 pm and reaching a minimum at E 2 am. These findings are consistent with those of Schirman-Hildesheim and collaborators^86^ and align with reports of a post-transcriptional regulation of *Gnrhr* transcripts^87,88^. Both RNA velocity and spliced/unspliced profile analysis further revealed alternating phases of transcriptional activation and repression, with velocity peaks coinciding with transcript reactivation, corroborating this dynamic regulation (Figure 8C).

We then assessed chromatin accessibility at the proximal promoter only, as no associated enhancers could be detected. Accessibility showed no correlation with *Gnrhr* transcript levels (Supplementary Figure S7.1D-E), suggesting that chromatin state changes are unlikely to drive *Gnrhr* expression variation across the estrus cycle. Decades of research have established that *Gnrhr* is regulated by multiple TFs acting on the proximal promoter during gonadotrope differentiation (reviewed in ^89,90^), with NR5A1, ERα, ISL1, LHX3, and GATA2 playing central roles. Moreover, we recently identified NEUROD4 as a novel additional regulator of *Gnrhr* expression in embryonic gonadotrope cells^29^. In the present study, we found that all these TFs, except LHX3 (Supplementary Figure S6), exhibit cyclic variations in gonadotrope cells throughout the estrus cycle. *Nr5a1*, *Isl1*, *Gata2*, and *Esr1* (coding ERα) displayed broadly similar expression profiles (Figure 8E), peaking during late D2-early PE and subsequently declining, but these dynamics did not correlate with *Gnrhr* regulation. In contrast, *Neurod4* expression (but not *Neurod1,* Supplementary Figure S6) closely correlated with *Gnrhr* profile (Figure 8E, r = 0.86). These results suggest that NEUROD4 is the key activator of *Gnrhr* during the estrus cycle. NR5A1, ISL1, and ERα may contribute to *Gnrhr* expression specifically during D2, while GATA2, a known repressor of *Gnrhr* expression^91^, likely mediates transient repression.

Overall, our results suggest that previously described mechanisms are insufficient to account for the dynamic expression of *Lhb*, *Fshb*, and *Gnrhr* during the PE surge, motivating a comprehensive reinvestigation of transcriptomic regulation in gonadotropes across the estrus cycle.

### Dynamic Reorganisation of Gonadotrope Gene Regulatory Networks Across the Estrus Cycle

To dissect the regulatory programs shaping gonadotrope activity during the estrus cycle, we inferred GRNs from snMultiome data using the scMega package^92^ adapted to the *Rattus norvegicus* rn7 genome. Candidate TFs and their putative target genes were identified in gonadotrope cells. TF binding activity, estimated from chromatin accessibility, correlated strongly with TF expression (Figure 9A). Similarly, the dynamic of TF expression and of their differentially expressed target genes were tightly correlated across regulatory time (Figure 9B). These results confirm that our dataset captured biologically relevant regulations in gonadotropes. Strikingly, this analysis also revealed a major regulatory transition from early-to-late PE (Figure 9A-B and Supplementary Figure S8A), further supporting the idea that a large-scale reprogramming of the gonadotrope transcriptional landscape occurs at this moment.

Network inference identified two distinct subnetworks (Figure 9C): a smaller Early-GRN, comprising genes expressed before the surge, and a larger Late-GRN, encompassing genes induced during and after the surge. Thus, gonadotrope regulation is organised in a biphasic manner, with distinct regulatory architectures operating before and after ovulatory priming.

The Early-GRN appears to be decentralised, *i.e.* composed of multiple interconnected TFs without a single dominant regulator. Among the most highly connected nodes are ERα, CTCF, NR5A1, and RFX1/3; however, only ERα, displaying high connectivity, might function as a network integrator (Figure 9C; Supplementary Figure S8B-C&F). This result is consistent with its well-established role in mediating estrogen regulation of gonadotrope physiology^93^. *In silico* perturbation assays (Figure 9D) further indicated that removal of these highly connected nodes would cause the strongest disruptions in the Early-GRN.

These predictions identify already established gonadotrope regulators such as NR5A1^94^ but also highlight less expected factors. For example, the involvement of CTCF implicates upstream chromatin remodelling as a potential prerequisite for the gonadotropin surge, and the identification of RFX1/3, known for roles in ciliogenesis, raises the possibility of previously unrecognised regulatory functions of these factors in pituitary adult gonadotrope cells.

That the Early-GRN relies on multiple interconnected TFs may confer robustness, while its decentralised topology could enable an adaptative program capable of integrating diverse upstream signals before the gonadotropin surge.

By contrast, the Late-GRN is hierarchical and hub-driven, composed of a handful of strongly connected TFs, including JUNB/D, FOS, FOSL1/2, BACH1/2, and MEF2A/D (Figure 9C-D; Supplementary Figure S8B-F). Among these, FOSL2, BACH1, and MEF2A emerge as key integrating hubs, reflecting a shift toward consolidated transcriptional control during and after the surge. Unexpectedly, BACH1 and BACH2, whose roles in gonadotropes have not been reported, also emerged as important hubs. Notably, a recent preprint reported that BACH1/2, together with FOSL1/2, JUNB, and JUND, coordinate transcriptional programs in T cells^95^, suggesting potential conserved regulatory functions across cell types. Importantly, all these hub TFs are predicted regulators of *Fshb* (Figure 9E), with expression profiles strongly correlated with *Fshb* velocity (Figure 9F), suggesting that *Fshb* transcription is orchestrated by a coordinated Late-GRN module to ensure its robust activation during the preovulatory surge.

#### Regulatory architecture of *Fshb* gene under GnRH

To validate GRN predictions, we used LβT2 gonadotrope cells under a sustained GnRH stimulation paradigm mimicking the 8-10h PE exposure (Figure 10A). Treatment with the GnRH superagonist Triptorelin (GnRHa) led to an early and transient induction of *Lhb* and *Gnrhr*, followed by prolonged repression, mirroring *in vivo* dynamics. *Fshb* displayed a striking biphasic response, with a brief early rise and a delayed, robust peak after 8h of GnRH treatment. Stimulation of LβT2 cells thus reproduces physiological gonadotropin gene dynamics, establishing this system as a relevant model to dissect the molecular mechanisms regulating *Fshb*.

**Figure 10:**
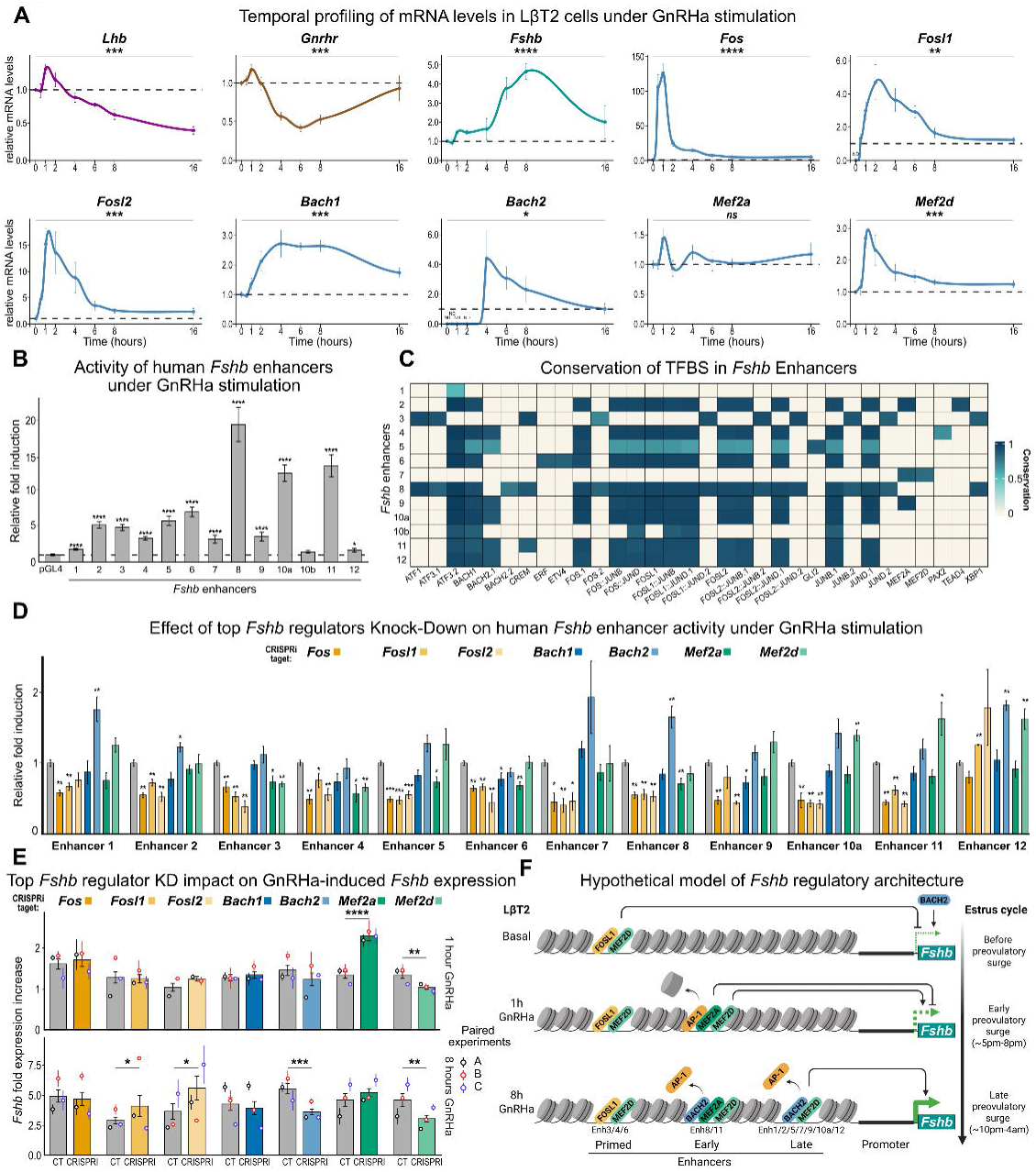
Deciphering the GnRH dependent *Fshb* regulatory architecture. **(A)** Temporal profiling of mRNA levels in LβT2 cells under GnRHa stimulation for *Lhb, Gnrhr, Fshb* and some TF predicted to regulate *Fshb* expression throughout the estrus cycle. Transcript levels were measured by RT-qPCR in LβT2 cells either untreated or treated with 10nM GnRHa for indicated times. Data were normalised to two housekeeping genes, *Ppia* and *Rplp0* and are presented as mean +/- SEM mRNA level relative to either the untreated cells or the lowest expression point when the transcript was not detected in untreated cells (ND). Expression changes over time were evaluated using a non-parametric Kruskal-Wallis test on normalised log-transformed values (log10), with Benjamini-Hochberg correction applied to the global p-values. *p < 0.05, **p < 0.01, ***p < 0.001 and **** p < 0.0001 (N>3 per time point). **(B)** Analysis of the activity of human *Fshb* enhancers under 8h of GnRHa stimulation. LβT2 cells were transiently transfected with a GL4 luciferase reporter cassette under the control of each enhancer or with pGL4 empty backbone used as control. Data represent the mean +/- SEM of treated over untreated fold induction, normalised to pGL4. Differences were evaluated using a linear mixed-effects model on log10-transformed values, including experiment identity to account for experiment-to-experiment variability, followed by a Dunnett post-hoc test against pGL4 with Benjamini–Hochberg correction. *p < 0.05 and **** p < 0.0001 (N>5). **(C)** Cross-species conservation of predicted TFBS across the twelve *Fshb* enhancers. Each column denotes a TF motif identified by TOMTOM against the JASPAR vertebrate database, and each row corresponds to an individual enhancer. For each TFBS detected within an enhancer, a composite conservation score was calculated by combining (i) the evolutionary conservation of the aligned sequence across 26 mammalian species including Human and (ii) its similarity to the JASPAR consensus position weight matrix. Scores were normalised between 0 and 1. The colour scale reflects both conservation and motif–sequence agreement. **(D)** Impact of the knock-down of top *Fshb* regulators on the GnRHa-induced activity of the human *Fshb* enhancers. LβT2 cells were transiently transfected with a GL4 luciferase reporter cassette under the control of each enhancer, along with CRISPRi targeting each of the top *Fshb* regulators: *Fos*, *Fosl1*, *Fosl2*, *Bach1*, *Bach2*, *Mef2a* or *Mef2d*. Irrelevant gRNA was used as untargeting control. Enhancer activity was assessed after 8h of stimulation with GnRHa. Data represent mean +/- SEM of fold induction, normalised to both pGL4 and untargeting gRNA. Differences relative to control were evaluated using Wilcoxon rank-sum tests, with Benjamini-Hochberg correction applied within each enhancer. *p < 0.05, **p < 0.01 and ***p < 0.001 (N>5). **(E)** Impact of the knock-down of the top *Fshb* regulators on GnRHa-induced *Fshb* expression. LβT2 cells were transiently transfected with CRISPRi targeting each of the top *Fshb* regulators: *Fos*, *Fosl1*, *Fosl2*, *Bach1*, *Bach2*, *Mef2a* or *Mef2d*. Irrelevant gRNA was used as untargeting control. Data are shown as mean +/- SEM of mRNA induction. Paired CT-CRISPRi experiments are plotted using different colours (experiments A, B and C). Statistical analysis used a linear mixed-effects model on log10-transformed induction values, including experiment identity to account for experiment-to-experiment variability, followed by a Dunnett post-hoc test versus control with Benjamini-Hochberg correction. *p < 0.05, **p < 0.01, ***p < 0.001 and **** p < 0.001 (N=3, n=9). **(F)** Hypothetical model of *Fshb* transcriptional regulation during the estrus cycle. Simplified representation of chromatin accessibility at the *Fshb* locus in the basal (unstimulated) state, and after early (1h) and late (8h) GnRH stimulation. Enhancers active at basal state are defined as primed. Enhancers activated after 1h or 8h of GnRH stimulation are defined as early or late enhancers, respectively. The effects of enhancer-TF interactions on *Fshb* expression are indicated by black lines (blunt arrow for inhibition, sharp arrow for activation). The resulting *Fshb* expression output is shown by the green arrow.

To investigate these mechanisms and their conservation in Humans, we cloned the human orthologs of *Fshb* conserved enhancers and assessed their activity in LβT2 cells using luciferase reporters. Basal activity was suppressed for most enhancers, whereas Enh3, Enh4, and Enh6 exhibited clear basal activation. Upon GnRH stimulation, only Enh8 and 11 were activated as early as 1h post-treatment, suggesting a role in early *Fshb* regulation. Finally, most enhancers were strongly induced after 8h of GnRHa stimulation (Figure 10B, Supplementary Figure S9A). In contrast, Enh10b was unresponsive to GnRH, likely ruling out involvement in *Fshb* expression control.

Interestingly, amongst the other expressed genes within the *Fshb* TAD (*Kcna4*, *Dcdc5*, *Kif18a,* Supplementary Figure S9B), only *Kcna4* exhibited an expression profile mirroring *Fshb*, suggesting potential co-regulation by the same enhancers. Together, these results demonstrate that the identified enhancers exhibit activation dynamics consistent with a functional role in *Fshb* regulation, highlighting conservation of regulatory mechanisms between rats and Humans.

Analysis of motif conservation across 26 mammalian species revealed the presence of highly conserved AP-1 motifs in all *Fshb* enhancers, BACH1/2 motifs in most enhancers (10 over 13), and MEF2 motifs in enh2, 7, and 9 only (Figure 10C, Supplementary Table 7). These data support a model in which these GRN-predicted TFs orchestrate *Fshb* expression *via* enhancer modulation. Expression profiling in LβT2 cells after GnRHa stimulation revealed that a fast increase in *Fos*, *Fosl1*, *Fosl2*, and *Mef2d* expression levels, peaking 1-2h post-treatment, while *Bach1* and *Bach2* expression peaked later (∼6h), and *Mef2a* levels remained stable (Figure 10A). These dynamics mirror the *in vivo* induction during PE night (Figure 9) and suggest distinct temporal contribution in *Fshb* regulation. We then measured the effect of TF knock-down on GnRHa-stimulated enhancer activity (Figure 10D). TF knockdown efficiency by CRISPRi was confirmed by RT-qPCR (60-80% expression decreases) (Supplementary Figure S9C). Results showed that FOS, FOSL1, or FOSL2 contribute to GnRH regulation of most enhancers. BACH1 increased GnRHa activity induction of Enh6 and 9 while BACH2 reduced it on Enh1, 2, 8 and 12. MEF2A increased GnRHa induction of Enh3, 4, 5, 6 and 8, whereas the effect of MEF2D was sequence-dependent, increasing induction of Enh3 and 4, while decreasing GnRHa induction of enh10a, 11 and 12. These results demonstrate that the GRN analysis successfully identified functionally relevant enhancer-TF interactions.

Finally, we examined the involvement of these TFs in GnRH regulation of endogenous *Fshb* expression by CRISPRi TF knock-down. Basal *Fshb* gene expression was increased by BACH2 and decreased by both FOSL1 and MEF2D (Supplementary Figure S9D).

GnRHa-induced *Fshb* expression was transiently reduced by MEF2A after 1h, whereas MEF2D enhanced expression at both 1 and 8h of stimulation. After 8h, FOSL1/2 decreased *Fshb* induction, while BACH2 and MEF2D further promoted its expression (Figure 10E). Knockdown of these TFs did not alter GnRHa-induced expression of other genes within the *Fshb* TAD, except for FOS-mediated activation and MEF2A-dependent repression of *Kcna4* at 8h post-treatment (Supplementary Figure S9E-G), further suggesting that *Kcna4* and *Fshb* share common regulatory mechanisms.

In summary, we uncover a novel regulatory architecture controlling *Fshb* expression during the PE night, involving newly identified enhancers and TFs, and provide strong experimental support for the GRN model inferred from snMultiome data.

## Discussion

Irregular menstrual cycles and dysovulation affect millions of women worldwide, yet the molecular mechanisms that orchestrate ovulation timing remain poorly understood. Central to this regulation are pituitary gonadotropes, which dynamically adjust hormonal output to coordinate ovarian function. Despite their pivotal role, the plasticity of these cells across the female cycle has remained largely unexplored. Here, we combine high-resolution experimental profiling with computational modelling to reveal how gonadotropes remodel their regulatory networks, uncovering core mechanisms that govern reproductive cyclicity. This dataset captured epigenetic and transcriptional dynamics at single-nucleus resolution across the cycle, enabling time-resolved modeling of gene expression. The resulting data further supported the reconstruction of gene regulatory networks, which were then experimentally validated. This provides a robust predictive framework for dissecting the cyclic control of female reproduction by gonadotrope cells.

### Unexpected Epigenetic Plasticity in Adult Female Gonadotrope Cells

Epigenetic plasticity, allowing cells to remodel their chromatin landscape, underlies lineage specification through stable reprogramming of *cis*-regulatory architecture. In fully differentiated cells, this landscape is thought to be fixed, maintaining identity and function. When disrupted, it can drive pathological reprogramming, including oncogenic transformation.

Evidence of active epigenetic remodelling in adult somatic cells are scarce. Dynamic chromatin changes have been reported in neurons during learning^96^, in macrophages upon immune activation^97^, in hepatocytes during fasting^98^, and in skeletal muscle during exercise^99^, typically affecting about 6,000 ± 1,000 *cis*-regulatory elements. Yet the significance of these remodelling in differentiated cells remain unclear. Our data reveal extensive epigenomic reorganisation in adult female gonadotropes in a physiological context. Indeed, across the estrus cycle, ∼12,000 out of ∼100,000 *cis*-regulatory regions showed cyclic variations in chromatin accessibility, which represent over 10% of the regulatory landscape. The gonadotrope epigenome thus oscillates between at least two distinct accessibility states within each four-day cycle, indicative of a highly reversible reprogramming. This large-scale, periodic remodelling defines an “epigenomic elasticity,” a property not previously recognised in adult somatic cells. Ours findings demonstrate that terminally differentiated cells can undergo broad, cyclical reorganisation of their chromatin architecture, revealing that adult epigenetic states are inherently dynamic.

### Biphasic GRN Architecture as a Feature of Gonadotrope Plasticity

We propose that a transition from an Early-to Late-GRN drives the switch in gonadotrope activity across the estrus cycle. The pre-surge Early-GRN operates as a decentralised network, with distributed regulatory control over many TFs to maintain responsiveness to external cues. In contrast, the hierarchical Late-GRN enables coordinated yet transient activation of broad transcriptional programs during PE. This reorganisation is mediated by a small set of TFs acting as central nodes, including AP-1 family members, MEF2, and BACH factors. Late-GRN regulatory interactions converge on few central hubs, forming an architecture that likely drives the sharp, reversible transcriptional shift underlying the gonadotropin surge. Comparable network transitions occur during differentiation, where highly connected TFs stabilise lineage commitment^100,101^. The hub-driven architecture of the Late-GRN may likewise commit gonadotropes to a surge-compatible transcriptional program.

GRN phase transitions of this kind are rarely observed outside developmental contexts. Yet the pronounced epigenetic plasticity observed in some adult tissues parallels our findings, suggesting that GRN phase transitions may represent a general yet overlooked strategy by which differentiated cells adapt to changing physiological demands.

### Redefining the Molecular Basis of the Gonadotropin Surge

The gonadotropin surge has traditionally been ascribed to transcriptional activation of the β-subunit genes *Lhb* and *Fshb* under hypothalamic GnRH control. Transcriptional regulation of *Lhb* and *Fshb* was first characterised *in vivo* in the late 1980s^67,68^. In our study, *Fshb* upregulation preceded FSH secretion, whereas *Lhb* showed no corresponding induction before the LH surge, as confirmed by single-cell and RT-qPCR analyses. Several independent studies likewise failed to detect *Lhb* induction during PE in intact^102^ or in ovariectomised, estradiol-supplemented female rats^103^ as well as in ewes^104,105^. The foundational report of *Lhb* upregulation thus likely reflected methodological artifacts. Our findings demonstrate that LH and FSH surges rely on distinct mechanisms: the LH surge is primarily secretory and independent of new β-subunit synthesis, whereas the FSH surge is transcriptionally driven.

Previous studies using LβT2 cells showed that low-frequency GnRH pulses (one every two hours) favour *Fshb* expression, while high-frequency pulses (every 30 minutes) enhance *Lhb*^106^, suggesting frequency decoding by gonadotropes. However, our *in vivo* data challenge the relevance of this mechanism during PE. Recently reported recordings from GnRH neurons in female mice^107^ showed sustained firing throughout the PE-E night, reaching ∼20 Hz, far exceeding the pulse range required for frequency decoding. Such a near continuous activity argues against frequency-based control at this stage, though it may operate outside the PE-E night during the cycle or during perinatal or pubertal periods.

If frequency decoding does not operate during PE, other mechanisms must explain the temporal 8-10h lag between LH and FSH release. Yet the molecular basis of this dissociation remains unknown. The GnRH receptor couples to both Gαq/11-CaZ⁺-PKC and Gαs-cAMP-PKA pathways^108–111^, each contributing to gonadotropin synthesis and release under strong GnRH stimulation. Consistent with this dual coupling, our data showed induction of Gαq/11-CaZ⁺-PKC-dependent genes (*Fos*, *Junb*, *Jund*) and Gαs-cAMP-PKA-dependent genes (*Fst*, *Crem*) during the PE evening. The CaZ⁺-dependent pathway desensitises rapidly^112,113^, whereas cAMP signalling persists throughout the PE night^108,114^. Temporal separation of LH and FSH release may therefore result from differential pathway activation or distinct desensitisation kinetics.

A further regulatory layer involves the TGFβ-SMAD3 pathway, which modulates *Fshb* synthesis *via* activin or myostatin^115^, under negative control of both FST and inhibin B. During PE, reduced expression of *Tgfbr3l,* needed for inhibin B-mediated repression of activin^116^, combined with *Fst* induction, may transiently delay *Fshb* transcription onset. Because *Fst* remains elevated through the night, additional factors likely drive the late-phase activation of *Fshb* synthesis.

### New Model of Gonadotropin Gene Regulation During Proestrus

Chromatin accessibility and GRN analyses reveal a new regulatory framework for gonadotropin gene control across the estrus cycle. We observed minimal *Lhb* upregulation, consistent with reduced promoter and enhancer accessibility during the LH surge and absence of *Lhb* as a major GRN target. In contrast, *Fshb* occupies a central position in the late-phase GRN, associated with thirteen evolutionarily conserved candidate enhancers, suggesting that its regulation depends on coordinated enhancer interactions rather than a single dominant element. Twelve human orthologous enhancers were shown to be activated by GnRH in LβT2 cells, highlighting the conservation of *Fshb* regulatory architecture between rodents and Humans.

One enhancer, enh1, maps to a human locus linked to female fertility and polycystic ovary syndrome (PCOS)^117^ and exhibits dynamic activity across the estrus cycle, demonstrating that single-nucleus multiomic profiling in the rat pituitary can provide relevant information on Human reproductive physiology. Mutation of enh1 in mice only mildly impairs female fertility^118^, supporting functional redundancy among enhancers and a model in which *Fshb* transcription is shaped by a complex molecular network.

Most *Fshb* enhancers recapitulate transcriptional kinetics of *Fshb*: weak basal state activity, followed by strong but delayed activation in response to GnRH. In addition to being responsive to GnRH, two enhancers also contain activin-response motifs, enh1 with a conserved SMAD site^117^ and enh9 with both FOXL2 and FOXO3 motifs^119^. This suggestes that GnRH and activin signalling might converge at these enhancers to fine-tune *Fshb* activation during the PE phase.

GRN analysis and enhancer conservation predicted that late-GRN hub factors, including FOS, FOSL1/2, BACH1/2, and MEF2A/D, converge on *Fshb* enhancers. CRISPR interference in GnRH-treated LβT2 cells confirmed that all hubs modulate basal or GnRH-stimulated *Fshb* expression, validating the GRN model. Interestingly, AP-1 factors activated enhancers while repressing endogenous *Fshb* expression. Previous promoter-reporter studies identifying FOS as a potential activator^120^ however likely underappreciated the complex enhancer architecture revealed here. Beyond direct transcriptional effects, AP-1 factors have also been shown to promote enhancer accessibility^121,122^. By analogy, AP-1 TFs may similarly prime gonadotrope enhancers early in PE while transiently restraining *Fshb* transcription. Conversely, BACH2 expression peaks before to *Fshb* activation, and as a known inhibitor of AP-1^123^, it may relieve AP-1-mediated repression, thereby enabling *Fshb* expression. Finally, MEF2A and MEF2D also emerge as key modulators of *Fshb* enhancers. Although MEF2D has been reported to regulate *Fshb* in embryonic gonadotropes, this regulation did not implicate *Fshb* promoter^124^. Here we showed that MEF2D, and to a lesser extent MEF2A, potently regulate *Fshb* enhancers, potentially orchestrating epigenetic remodelling *via* both activator and repressor complexes^125^.

The paradoxical observation that some TFs activate enhancers while repressing *Fshb* is consistent with an incoherent feed-forward loop^126^, in which a regulator both promotes and restrains target expression. Such a regulatory switch, arising from competitive interactions between TFs and enhancers, provides a mechanistic explanation for the delayed induction of *Fshb* during the PE-E transition and the subsequent temporal uncoupling of LH and FSH surges. Collectively, these results support a model in which *Fshb* regulation under GnRH is governed by a dynamic transcriptional network (Figure 10F). At the basal state, most enhancers are repressed, with only a few ones already primed to maintain low *Fshb* expression under the antagonistic influence of both MEF2 and BACH TFs. GnRH may then recruit early enhancers *via* AP-1 TFs, yet transcription remains low due to MEF2A/D-mediated repression. Full *Fshb* expression is subsequently achieved through late BACH2 expression, which likely relieves this inhibition. This competitive interplay between TFs and enhancers likely enables precise temporal control of *Fshb* expression during the PE night, a hypothesis that deserves further experimental investigation.

### Cellular and Tissular Pituitary Plasticity During the Proestrus Night

Subcellular and cellular plasticity of gonadotropes has been reported in castrated male rats, where intense GnRH stimulation triggers extensive remodelling of the ER-Golgi network, vesicle trafficking, and secretory machinery, enabling polarised hormone release^127,128^. By analogy, our gene ontology analysis suggests that female gonadotropes undergo similar plasticity across the estrus cycle. We observed that gonadotropes also secrete numerous angiogenic-related factors, including TNC, DMP1, FGF1, and VEGFA^129^, likely supporting gonadotrope-driven modulation of pituitary architecture, including potential vascular remodelling. Together, these observations highlight a coordinated interplay between transcriptomic programs and cellular remodelling, indicating that female gonadotropes actively drive their own cellular plasticity during PE. Although compelling, these interpretations remain inferential and require further experimental validation.

Beyond gonadotropes, we also demonstrated that lactotropes and corticotropes exhibit transcriptomic plasticity across the cycle. In female rats, lactotropes drive the prolactin surge during the afternoon of PE, triggered by the release of dopamine inhibition and estradiol priming ^130^. Estradiol further regulates lactotrope proliferation^131^. The observed transcriptomic changes likely support these well-established physiological responses. We also observed unexpected corticotrope plasticity, predominantly during estrus, a phenomenon not previously described in females. Interestingly, a recent work indicates that FSH signalling in corticotropes restrains ACTH, and hence corticosterone production, thereby protecting against female-specific hepatic steatosis^132^. The physiological significance of lactotrope and corticotrope remodelling remains unclear, but it highlights a potentially overlooked paracrine coordination within the pituitary, potentially impacting female fertility and broader metabolic homeostasis.

In summary, our results demonstrate that adult female gonadotropes undergo reversible, cycle-dependent epigenetic and transcriptional reprogramming, coordinated by conserved hub factors and enhancers that drive the preovulatory surge. The conservation of these regulatory elements in Humans, including loci linked to fertility and PCOS, provides a mechanistic framework and candidate loci for better understanding female reproductive physiology and infertility.

## Material and Methods

### Animal care

Seventy-four female Wistar rats (8-weeks old) were obtained from Janvier Labs (Le Genest-Saint-Isle, France) and housed under a 12h-light:12h-dark cycle (lights on 7 am to 7 pm) ventilated cabinet open cages. After a 2-week acclimatisation period, estrus cyclicity was monitored for a further 2 weeks by analysis of daily vaginal smear cytology stained with May-Grünwald-Giemsa. Rats showing irregular cycles and later exhibiting anatomical abnormalities at necropsy (e.g., unilateral development of the reproductive tract or tumours) were excluded from analysis. Females with regular cycles were euthanised at defined stages of the estrus cycle: diestrus 2 (D2, 10 am), proestrus morning (PE, 10 am), then every 1-2 h from 4 pm to 6 am across the PE-E transition, and finally at E (10 am). Pituitaries were collected, halved, and snap-frozen for molecular analyses. Blood was collected for endpoint hormone profiling.

The experiments conducted on animal were conform to the best practice and the 3Rs rule and followed strict regulations under the control of *Ministère de l’enseignement superieur et de la recherche*. All procedures were made in accordance with current French and European laws and policies and have been examined and authorised (Approbation number APAFIS #43635-2023032416202556).

### Hormonal assays

Serum LH, FSH and PRL concentrations were simultaneously assayed with the Luminex technology using the rat pituitary magnetic bead panel Milliplex Map kit (RPTMAG-86K-Milliplex, Merck-Millipore, Nottingham, UK) following the manufacturer’s instructions.

Steroids (estradiol [E2], progesterone [P4], 4-androstenedione [4-dione], dihydrotestosterone [DHT], and testosterone [T]) were assayed by gas chromatography-mass spectrometry (GC-MS) at the Mondor Institut Mass Spectrometry plateform as previously described^133^.

### Nuclei preparation for Chromium Single-nucleus Multiome ATAC + Gene Expression (10x Genomics)

Frozen half-pituitaries were minced in lysis buffer (10 mM Tris-HCl, 10 mM NaCl, 3 mM MgCl_2_, 0.1% NP-40 in nuclease-free water) for 1 min and incubated for 9 min at 4 °C with gentle shaking. Lysates were homogenised on ice using a 2-mL Dounce homogeniser and diluted to 3 mL with wash buffer [1× PBS (Gibco), 2% BSA, 0.2 U/μL RNase inhibitor (Merck)]. Suspensions were filtered through 100-μm strainers (VWR), centrifuged (500 × g, 10 min, 4 °C), and resuspended in wash buffer.

Nuclei were incubated in 250 μL wash buffer with Alexa Fluor® 647 anti-Nuclear Pore Complex antibody (Mab414, BioLegend; 2 μg/μL, 15 min, 4 °C), washed, filtered through 35-μm strainers (Falcon 332235, Dominique Dutscher), and sorted on a BD FACSAria III (BD Biosciences).

For each pituitary, 10,000 nuclei were used for tagmentation and loaded onto the 10x Genomics Chromium Controller. ATAC-seq and RNA-seq libraries were prepared according to the manufacturer’s protocol (https://www.10xgenomics.com/support/single-cell-multiome-atac-plus-gene-expression) and sequenced on an Illumina NextSeq 500 (paired-end, 26 bp read 1, 10 bp dual index, 90 bp read 2) to a depth of ≥20,000 reads per nucleus.

### Reannotation of rn7 reference genome and RNA-seq and ATAC-seq processing

(https://github.com/Dalhte/snMultiome-Pituitary/snMultiome-processing)

To maximise alignment accuracy and annotation consistency in the rat, we built a custom reference for single-cell multiome analyses. Public bulk paired-end RNA-seq datasets (ENA accessions PRJEB39130, PRJNA183967, PRJCA004225) were aligned to the Rattus norvegicus mRatBN7.2 (rn7) assembly using STAR (v2.7+). The STAR genome index was generated from the matching FASTA and GTF (Rattus_norvegicus.mRatBN7.2.110). Alignments were processed with samtools (v1.12+) in the following order: name sort, fixmate, coordinate sort, duplicate removal. Strand-specific BAMs were derived using standard SAM flags; for downstream reference support, the strand-separated BAMs were then merged. Gene annotations were restricted to canonical chromosomes (chr1-chr20, chrX, chrY and chrM for mitochondrial DNA) and normalised with AGAT^134^ to produce a harmonised GTF. A Cell Ranger ARC reference genome (cellranger-arc v2.0.2) was then created using the improved annotation. The snRNA and snATAC libraries were then processed with cellranger-arc count to quantify gene expression and chromatin accessibility using the new reference genome.

### SnRNA-seq preprocessing and ambient RNA correction

(https://github.com/Dalhte/snMultiome-Pituitary/snMultiome-processing)

SnRNA-seq data were processed using the 10x Genomics Cell Ranger output as input for downstream analyses. For each sample, raw and filtered gene expression matrices were imported in R (v4.3.0) using Seurat (v5.0.1)^16^ and DropletUtils (v1.20.0). To account for ambient RNA contamination, we applied SoupX (v1.6.2)^135^. For each dataset, a SoupChannel object was created by combining raw and filtered matrices. Cluster assignments and UMAP embeddings were transferred to SoupX using setClusters and setDR. Contamination fractions were estimated with autoEstCont and manually adjusted to optimise removal of ambient transcripts. Corrected expression matrices were generated with adjustCounts and exported in 10x-compatible format using DropletUtils.

### SnATAC-seq preprocessing and peak set unification

(https://github.com/Dalhte/snMultiome-Pituitary/snMultiome-processing)

Accessible chromatin profiling was performed using the 10x Genomics ATAC platform. Peak calls for each dataset were obtained from Cell Ranger ATAC output files. Individual peak lists were converted to GenomicRanges objects using GenomicRanges (v1.52.0). To ensure consistent peak definitions across samples, we constructed a unified peak set by merging overlapping intervals using GenomicRanges. Peaks shorter than 20 bp or longer than 10 kb were excluded to remove technical artifacts and broad, non-informative regions. This unified peak set was used for downstream quantification of chromatin accessibility.

### SnMultiome data processing and integration

(https://github.com/Dalhte/snMultiome-Pituitary/snMultiome-processing)

Filtered single-cell RNA and ATAC data were processed using Seurat and Signac. SoupX-corrected gene expression matrices were incorporated as an independent assay alongside raw RNA counts. ATAC fragment files and barcode-level metadata were imported, and fragments were filtered to retain high-confidence cells. Chromatin accessibility was quantified using the unified peak set. A chromatin assay was then created with annotation from the Ensembl *Rattus norvegicus* genome (EnsDb.Rnorvegicus.v110).

Quality control included nucleosome signal estimation and TSS enrichment scoring, with cells failing mitochondrial content thresholds (>1%) or exhibiting <200 or >2500 detected RNA features were removed. RNA assays were normalised using SCTransform, regressing out mitochondrial percentage. ATAC assays were processed with TF-IDF normalisation, dimensionality reduction by latent semantic indexing (LSI), and UMAP embedding.

To enable cross-sample integration, Seurat objects from each sample were merged and cell barcodes uniquely prefixed by dataset of origin. Dataset-specific ATAC embeddings were aligned using reciprocal LSI (rLSI) integration with Seurat FindIntegrationAnchors and IntegrateEmbeddings. MACS3 was used to call peaks on the integrated object, and counts were re-quantified over this peak set to generate a harmonised chromatin accessibility assay.

Dimensionality reduction was performed independently on both RNA (PCA, UMAP) and ATAC (LSI, UMAP) modalities. Integrated embeddings were used for downstream visualisation and clustering.

To jointly analyse transcriptomic and chromatin accessibility profiles, we performed multi-modal integration using Seurat. A weighted nearest-neighbor (WNN) graph was constructed with FindMultiModalNeighbors, combining RNA and ATAC modalities with modality-specific weights.

Cells were embedded in a joint multimodal UMAP space, enabling integrated visualisation across datasets. Clustering was performed on the WNN graph (FindClusters, resolution = 0.4), yielding transcriptionally and epigenetically coherent cell groups. Clusters were visualised across RNA, ATAC, and multimodal embeddings for consistency.

Cluster identities were manually curated based on canonical transcriptomic and accessibility signatures, and annotations were assigned to pituitary cell cluster.

### Gene activity inference from chromatin accessibility

(https://github.com/Dalhte/snMultiome-Pituitary/snMultiome-processing)

To infer transcriptional potential directly from chromatin accessibility, we computed gene activity scores using the GeneActivity function in Signac. Accessibility fragments within promoter-proximal and gene body regions were aggregated per gene, generating a sparse gene-cell activity matrix. The resulting GeneActivity assay was log-normalised and scaled for downstream visualisation.

### Motif enrichment and transcription factor activity analysis

(https://github.com/Dalhte/snMultiome-Pituitary/snMultiome-processing)

To identify TFBS motifs enriched within accessible chromatin regions, we integrated JASPAR2022 core vertebrate position frequency matrices (PFMs) into the analysis. Motif instances were scanned across high-confidence peaks using motifmatchr. Motif deviation scores were computed with chromVAR (v1.22.0), correcting for GC content and sequencing biases.

To account for redundancy among closely related TFs, we further grouped motifs into JASPAR-defined motif families using publicly available clustering files. Root motifs were mapped to family-level consensus PWMs, which were rescanned across the unified peak set. ChromVAR deviation scores were then recalculated at the motif family level.

### Pairwise analysis of differential expression within cell types

(https://github.com/Dalhte/snMultiome-Pituitary/snMultiome-processing)

Within each cell type, differential expression across four time points (D2 10 am → PE 10 am → PE 6 pm → E 10 am) was assessed on SoupX-adjusted RNA. To mitigate imbalance, clusters were subsetted and randomly down-sampled per origin towards the gonadotrope population size. Pairwise contrasts were performed with FindMarkers (Wilcoxon rank-sum). Genes were considered differentially expressed if any comparison met |log_2_FC| > 1 and FDR < 1 × 10^-5^.

### Cross-species conservation analysis

(https://github.com/Dalhte/snMultiome-Pituitary/Conservation-analysis)

Rat peak coordinates, lifted from rn7 to hg38 using liftover^136^. Nearby regions were merged across gaps ≤50 bp, and the mean phastCons score per region was computed from PhyloP100-way conservation tracks. Each distribution was benchmarked against a chromosome, and width-matched shuffled background using Wilcoxon rank-sum and Kolmogorov-Smirnov tests. Statistical difference between the two distribution was assessed by one-sided permutation test (over 10,000 permutations).

### Regulatory time construction

(https://github.com/Dalhte/snMultiome-Pituitary/Regulatory-Time)

To assess the temporal progression of molecular regulation, cluster-specific trajectories were computed with the ArchR package on the multimodal (RNA + ATAC) UMAP, yielding a per-cell pseudotime.

RNA velocity was estimated with scVelo^32^ from velocyto-derived .loom files. Looms were concatenated into a single AnnData object with spliced/unspliced layers; Seurat exports were merged by exact barcode match. scVelo preprocessing used filter_and_normalise, followed by moments, and the dynamical model to infer latent time.

Pseudotime values were projected onto a circular coordinate system, where the azimuthal angle θ represented pseudotime and the polar angle φ represent the scaled latent time. To capture periodic regulatory dynamics, we fitted generalised additive models (GAMs) with cyclic cubic splines. Two models were fitted sequentially: i) a quality-control model to confirm cyclicity and smoothness of the projection, and ii) a regulatory model that aligned latent time with pseudotime. This procedure yielded a single, continuous measure of “regulatory time”, integrating both pseudotime ordering and RNA velocity-derived latent time.

To assess the proportion of variance in pseudotime, latent time, and regulatory time explained by cell origin, we performed an ANOVA on each trajectory using the sample of origin as the grouping factor. Eta-squared (ηZ) was calculated from each ANOVA model to quantify the effect size, representing the fraction of total variability in each trajectory attributable to differences between groups.

Trajectory modelling and feature selection across molecular layers (mRNA abundance, RNA velocity, and chromatin accessibility) were projected onto regulatory time and summarised as 100-bin trajectories. Features were variance-filtered using a robust rule (row s.d. > median + 1.5 × MAD), spline-smoothed (smooth.spline, spar = 0.45), and tested for temporal variation by one-way ANOVA across regulatory time with Benjamini-Hochberg control (q < 1 × 10^-3^). For timing-based stratification, rows were ordered by a plateau-based “mode-like” time index and segmented with piecewise linear models (segmented; BIC selection; minimum segment length 100 rows). The number of segments K was set by a relative-BIC elbow, choosing the smallest k for which adding another segment reduced BIC by <15%, to ensure parsimony.

### Peak-gene association and regulatory class

(https://github.com/Dalhte/snMultiome-Pituitary/Regulome)

Candidate peak-gene links were defined by 3D context and promoter proximity. TADs were defined per chromosome as the intervals between previously described Hi-C boundaries in rats ^22^. For each gene, ATAC and RNA profiles (100 bins) were spline-smoothed (spar = 0.45). Post-smoothing Pearson correlation coefficients (r) with two-sided P-values were computed for all region-gene pairs; links were retained at FDR < 0.01 (with Benjamini-Hochberg correction). Empirical correlation cut-offs were estimated by circular permutation: for up to 50,000 significant non-promoter pairs, the RNA series was randomly phase-shifted and re-correlated with the original ATAC series (over 1,000 permutations). The 95th and 5th percentiles of this random distribution defined the positive and negative thresholds, respectively. Final classes were defined as: promoters, non-promoter “enhancers” (r above the positive threshold), and “silencers” (r below the negative threshold).

### Genes Functional enrichment

(https://github.com/Dalhte/snMultiome-Pituitary/GO)

Functional enrichment was performed on differentially expressed genes for each expression cluster (E1-E4). For each cluster, gene lists as well as their Human orthologous were tested for over-representation against a background comprising all genes present in the expression matrix. Gene Ontology (all aspects), KEGG, MsigDB Hallmark and Reactome gene sets were queried. One-sided hypergeometric tests were run *via* compareCluster with a minimal gene-set size of 5; FDR was controlled using Benjamini-Hochberg and results retained at q ≤ 0.05. To reduce redundancy, GO results were simplified by semantic similarity (Wang metric; cut-off = 0.60), after which all sources were clustered by gene overlap (Jaccard; average linkage; cut height = 0.40) and collapsed within clusters. For each representative term, counts, rich factor, fold enrichment, z-scores, p-values, and segment-wise BH-adjusted Q-values were recomputed using the segment-specific universe.

### Gene regulatory network inference and analysis

(https://github.com/Dalhte/snMultiome-Pituitary/Regulome.)

Gene regulatory network was assembled using the scMEGA package^92^. Briefly, TF activity, chromatin accessibility, and RNA abundance along regulatory time were integrated. Potential TF regulators were identified by correlating each TF activity trajectory with its matched RNA trajectory and retaining TFs with BH-adjusted p < 0.01 and correlation r > 0.30. Peak-to-gene links were called on trajectory-level matrices within TADs with gene position defined by their transcription start site. For each peak-gene pair, the Pearson correlation across 100 regulatory time units between accessibility and expression was computed, and positively correlated pairs were retained at FDR with BH q < 1 × 10^-4^. Signac motif-by-peak matrix was intersected with retained peaks and TFs. Directed TF to gene links were created when at least one linked peak carried the TF motif and the expression of the TF and gene were correlated. Each link/edge received a “regulatory score” equal to the mean of three z-scored components: TF-gene correlation, log₁₀(n_peaks + 1) (n_peaks = number of supporting peaks), and -log₁₀(FDR). High-confidence analyses retained positively concordant TF to gene edges with r > 0.75. The GRN was formated as a tidygraph and characterised with centralities (in-, out-, and total degree; closeness; betweenness using 1 / abs(regulatory_score) as the edge weight; eigenvector; PageRank).

An *in silico* perturbation analysis was performed by removing sequentially all outgoing edges of each TF and measuring network-level changes.

Nodes and edges with time annotations were exported for Gephi visualisation^137^.

### *Fshb* associated enhancers alignments and motif analysis

(https://github.com/Dalhte/snMultiome-Pituitary/Conservation-analysis)

Orthologous enhancer sequences were first identified using BLAT searches against the UCSC Genome Browser^138^ across 26 mammal species: Rat (rn7), Human (hg38), Mouse, (mm9), Chimp (panTro6), Gorilla (gorGor6), Dog (canFam4), Cow (bosTau9), Horse (equCab3), Rabbit (oryCun2), Opossum (monDom5), Female Naked mole rat (hetGla2), Guinea Pig (cavPor3), Squirrel (speTri2), Gibbon (nomLeu3), Baboon (papAnu4), Tarsier (tarSyr2), Alpaca (vicPac2), Panda (ailMel1), Sloth (choHof1), Elephant (loxAfr3), Cat (felCat9), Marmoset (calJac4), Chinese pangolin (manPen1), Ferret (musFur1), Hedgehog (eriEur2), Pig (susScr11) and Mouse lemur (micMur2). Retrieved orthologous regions were aligned using MAFFT^139^. When necessary, the sequences automatically reverse-complemented, ensuring consistent orientation across species. These multiple sequence alignments served as the basis for downstream conservation analyses. Multiple sequence alignments of enhancer regions were processed to extract conserved blocks of potential TFBS. A block was considered conserved if at least 80% of bases were identical across species and the block extended over a minimum of 6 bp. Positions containing gaps in more than 25% of sequences were excluded. To tolerate moderate variability, up to 20% of columns and up to three consecutive poorly conserved positions were allowed within a block. Each block was further subdivided using a sliding window of 20 bp with a 5 bp step, ensuring that overlapping candidate motifs were captured across the entire region. The resulting motifs were exported in MEME format and compared against the JASPAR vertebrate database using TOMTOM tool of the MEME suite^140,141^, with internal matches allowed, a minimum overlap of 5 bp, and a significance threshold of 0.2. The TFBS conservation of predicted *Fshb* regulators on *Fshb* enhancers was then assessed by integrating the evolutionary conservation of each aligned motif with its similarity to the corresponding JASPAR consensus. For each TOMTOM-identified site, a single composite conservation score was computed by combining a motif-to-PWM (Position-Specific Weight Matrices) similarity score and a base evolutionary conservation score.

### RNA extraction and RT-qPCR assays

Total RNA from rat anterior pituitary tissue was isolated with the RNeasy Kit (Qiagen, France). Two micrograms of total RNA were reverse-transcribed with SuperScript II (Thermo Fisher Scientific) and random primers, according to the manufacturer’s instructions. Real-time PCR was performed using Takyon™ 2X MasterMix (Eurogentec) on a LightCycler 480 instrument. Changes in candidate gene mRNA were assessed by RT-qPCR only when total pituitary mRNA variation could be attributed to a specific cell type (*i.e*., mRNA level > 3-fold higher than the average across clusters, fold-change ≥ 2, and adjusted p < 10^-5^). Expression levels of genes of interest were normalised to *Ppia* and *Ywhae* (transcripts stably and homogeneously expressed in all pituitary cell types).

LβT2 gonadotrope cell line RNA was extracted using TRIzol™ Reagent, following the manufacturer’s protocol. Reverse transcription and mRNA quantification were conducted as above. Expression was normalised to *Ppia* and *Rplp0*.

All oligonucleotide primer sequences (listed in Supplementary Table 8) were designed to target all known transcripts of each gene according to the Ensembl database, with amplicons spanning at least one intron (≥ 500 bp) to prevent amplification of genomic DNA.

### Immunohistofluorescence

Hemi-pituitaries were fixed in PBS containing 4% PFA for up to 24h at 4°C, incubated in 30% sucrose, embedded in OCT compound (CellPath, Newtown, UK), and then frozen at -80 °C for cryosectioning (15 µm-thick sections). Cryosections were post-fixed in 4% PFA in PBS for 15 min, blocked in 10% goat serum in PBS for 1h, and then incubated overnight at 4 °C with anti-LHβ (gift of Dr J. F. Roser, University of California, Davis, USA) and anti-NR5A1 (KO611) primary antibodies. Alexa Fluor-conjugated secondary antibodies (Invitrogen) were used. Nuclei were counterstained with DAPI, and slides were mounted in ProLong Gold Antifade Mountant (Thermo Fisher Scientific) for imaging. Images were acquired on a Zeiss LSM 700 confocal microscope at the Functional and Adaptive Biology Unit microscopy facility. For each time point, at least 10 regions of interest from three different pituitary sections of three pituitaries were analysed. To reduce positional bias, images were captured at standardised anatomical positions. Images were acquired as z-stacks, and the maximal NR5A1 signal in each nucleus of LH-positive cells was measured. Background in LH-negative cells was also measured and subtracted.

### Cell culture and GnRH stimulation

The LβT2 mouse gonadotrope cell line (generously provided by P. Mellon, University of California, La Jolla, CA, USA) was maintained in high-glucose DMEM supplemented with 10% foetal bovine serum (FBS; Gibco) and 0.1% penicillin-streptomycin at 37 °C in 5% CO_2_. Cells were regularly tested for mycoplasma and for cell identity.

To examine gonadotrope cell responses to GnRH, LβT2 cells were seeded at 250,000 cells/cmZ in PBS-rinsed wells coated with growth factor-reduced (GFR) Matrigel® (2%). Fifty-six hours after seeding, cells were serum-starved (FBS-free) for 16 h and then treated with GnRH agonist triptorelin (GnRHa, 10 nM) or left untreated.

### CRISPRi gene silencing

For each targeted genes, the top 5 gRNAs targeting proximal promoters (as described in^142^ and Supplementary Table 8) were cloned into psgRNA (Addgene #114005), and pooled. Cells were co-transfected with pLV hUbC-Zim3KRAB-dCas9-T2A-GFP (Addgene #53191), expressing a catalytically inactive Cas9 (dCas9) fused to the KRAB repression domain of human ZIM3, together with the gRNA plasmids, as described in^143^. Approximately 5 × 10⁶ LβT2 cells were electroporated with the dCas9-Zim3 and sgRNA plasmids at a ratio of 1.5:8.5 µg, using a Neon® Transfection System (Invitrogen) following the manufacturer’s protocol (two pulses at 1500 mV for 15 ms). The empty pSPgRNA plasmid was used as a control.

### Luciferase assay

Putative *cis*-regulatory regions of *Fshb* were amplified from human genomic DNA using Platinum™ *SuperFi* II DNA Polymerase (Thermo Fisher Scientific) and cloned into the pGL4.12 vector (Promega).

For GnRH stimulation assay, approximatively 10,000 cells per 384 wells-plate (Revvity) were transfected with Lipofectamine 3000 (Thermo Fisher Scientific) using 30 ng per well of the pGL4 construct and 4 ng per well of the pRL-null Renilla plasmid (Promega) as an internal control for normalisation.

For CRISPRi silencing assays, approximatively 10,000 cells per 384 wells-plate were transfected with Lipofectamine 3000 using 15 ng per well of the pGL4 construct, 2 ng per well of the pRL-null Renilla plasmid, 6 ng per well of the pLV hUbC-Zim3KRAB-dCas9-T2A-GFP plasmid and 16 ng per well of gRNA targeting pool. The empty pSPgRNA plasmid was used as a control.

At 48 h post-transfection, cells were serum-deprived overnight and then treated with 10 nM triptorelin for the indicated durations. Firefly and Renilla luciferase activities were measured using the Dual-Luciferase Reporter Assay System (Promega). Experiments were performed independently three times, each in triplicate. Data were normalised to Renilla luciferase for each condition.

### Data and code availability

Raw single-nucleus data as well as the analysis pipelines are respectively available on EMBL-EBI arrayexpress (E-MTAB-16103) and at github.com/Dalhte/snMultiome-Pituitary/.

### Use of AI-assisted tools

We used ChatGPT v5 (OpenAI) for two purposes: (i) drafting some initial skeleton scripts in R and Python and suggesting optimisations, and (ii) enhancing the manuscript’s language quality, including clarity, grammar, and spelling, without contributing new scientific content. All AI-generated outputs were critically reviewed, edited, and validated by the authors.

---

Figure S1: **Cytological characterisation of estrus cycle progression.**

**(A)** Representative images of vaginal smears from Diestrus 1, Diestrus 2, Proestrus and Estrus female rats, stained with May-Grünwald-Giemsa.

**(B)** Fifteen-day monitoring of the sexual cycle in the four individuals analysed using snMultiome technology. Individuals were sacrificed on day 15 and were in Diestrus 2 (10 am, Rat n°1), early and late Proestrus (10 am, Rat n°2 and 6 pm, Rat n°3) and Estrus (10 am, Rat n°4).

Figure S2: **Pituitary cell type characterisation by snRNAseq and snATACseq.**

**(A)** snRNAseq UMAP embedding with cells coloured according to the sample origin.

**(B)** snRNAseq UMAP embedding showing the expression of marker genes used for cell type identification.

**(C)** snATACseq UMAP embedding with cells coloured according to the sample origin.

**(D)** snATACseq UMAP embedding showing chromatin activity of marker genes used for cell type identification.

**(E)** snATACseq UMAP embedding showing TFBS enrichment in accessible chromatin regions, highlighting the most enriched TFBS in each pituitary cell type.

Figure S3: **Expression patterns of differentially expressed genes throughout the estrus cycle.**

For each gene: multimodal UMAP embedding of gene expression is represented in the upper panel and mRNA levels measured by RT-qPCR in the lower panel.

RT-qPCR analyses were only conducted if the expected variation could be attributed to the mentioned cell type according to mRNA quantification in other cell clusters (mRNA level > 3-fold higher than the average across clusters, fold-change ≥ 2, and adjusted p < 10^-^⁵). RT-qPCR analyses were performed on 69 hemi-pituitaries, with at least three individuals per time point. Sampling times were: D2 10 am; PE 10 am, 4 pm, 5 pm, 6 pm, 8 pm, 10 pm, 12 am; and E 2 am, 4 am, 6 am, 10 am. Data are presented as relative mRNA levels normalised to two housekeeping genes, *Ppia* and *Ywhae*. Significant variation throughout the estrus cycle was confirmed using a non-parametric Kruskal-Wallis test followed by Dunn post-hoc tests with Benjamini-Hochberg correction applied to the global p-values. ns non significant, * p < 0.05, ** p < 0.01, *** p < 0.001 and **** p < 0.0001.

**(S3.1-2)** Differentially expressed genes in gonadotrope cells.

**(S3.3)** Differentially expressed genes in lactotrope (*Gal*, *Gria4*, *S100g*); corticotrope (*Ccnjl*, *Trim66*); somatotrope (*Ghrhr*, *Tmc2*); thyrotrope (*Thy1*); melanotrope (*Brinp3*); folliculostellate (*Adamts9*, *Kcna2*); and endothelial cells (*Akap12*, *Sh2d3c*, *Tgfbr2*).

Figure S4: **Pituitary transcriptional changes throughout the estrus cycle.**

**(A)** Transcriptome of gonadotrope cells at PE 6 pm compared with D2 10 am. Differential expression was considered significant for genes showing at least a twofold change (vertical black dotted lines) with an adjusted p-value lower than 10^-^⁵ (horizontal black dotted line). Upregulated genes at PE 6 pm are shown in red, down-regulated genes in blue, and non-significantly regulated genes in grey.

**(B)** UMAP projection and pseudotime trajectory of lactotrope, corticotrope, somatotrope, thyrotrope, melanotrope, folliculostellate, endothelial, pericyte and leukocyte cells. The pseudotime trajectory illustrates the transcriptome temporal dynamics within the multimodal UMAP space. Each point corresponds to an individual cell, coloured according to its progression along the trajectory from the initial state (dark blue, pseudotime = 0) to the most advanced state (yellow, pseudotime = 100).

Figure S5: **Expression pattern of selected genes in pituitary cells.**

Multimodal UMAP embedding of mRNA levels for genes analysed by RT-qPCR in Figures 7 and 8. All the genes are enriched in the gonadotrope cell cluster.

Figure S6: **snRNAseq analysis of gene expression patterns in gonadotrope cells.**

Log2-normalised mRNA levels variation across the regulatory time of differentially expressed genes. Top panels: secretion related genes. Bottom panels: previously described gonadotrope molecular regulators.

Figure S7: **Characterisation of *Lhb*, *Gnrhr* and *Fshb* associated regulatory elements.**

**(S7.1)** Coverage plot of *Lhb* (**A**) and *Gnrhr* (**D**) expression and associated chromatin accessibility in each pituitary cell type. The studied DNA region is shown at the bottom of the plot, while its accessibility in each cell type is shown at the top. Accessible chromatin regions are indicated by grey boxes. Genome coordinates (bp) are from the rn7 assembly of the rat genome. Gene expression levels in each cell type are shown on the right of the plot. Each accessible region statistically associated with the gene expression is marked by a vertical line. Characterisation of region associated with *Lhb* (**B-C**) or *Gnrhr* (**E**) expression. Top panel: pseudo-bulk genomic region accessibility in the gonadotrope cells of the four female rats. Middle panel: Region conservation score across 20 vertebrates according to the PhastCons database. Bottom panel: chromatin region accessibility in gonadotrope cells throughout regulatory time. Values are shown as Log2-scaled accessibility obtained with the Signac package.

**(S7.2)** Coverage plot of *Fshb* expression and associated chromatin accessibility in each pituitary cell type (**A**). The studied genomic region is shown at the bottom of the plot, while its accessibility in each cell type is shown at the top. Accessible regions are indicated by grey boxes. Genome coordinates (bp) are from the rn7 assembly of the rat genome. *Fshb* expression levels in each cell type is shown on the right of the plot. Each accessible region statistically associated with *Fshb* expression is marked by a vertical line and numbered. (**B-M**) Characterisation of each of the regions associated with *Fshb* expression. First panel: genomic region accessibility in each pituitary cell type. Second panel: pseudo-bulk in the gonadotrope cells of the four female rats. Third panel: Region conservation score across 20 vertebrates according to the PhastCons database. Fourth panel: chromatin region accessibility in gonadotrope cells throughout regulatory time. Values are shown as Log2-scaled accessibility obtained with the Signac package.

Figure S8: **Analysis of gene regulatory networks in gonadotrope cells.**

**(A)** Distribution of the number of DEG with peak expression according to the regulatory time. This distribution appears to be bimodal and the red line marks the separation between the two underlying distributions. The associated regulatory time value was used as a cutpoint to identify the “Early” and “Late” subnetworks in figure 9C.

**(B-F)** Rankings of TFs within the GRN, colored by subnetwork assignment (Early in pink; Late in blue). Each panel shows a different centrality metric: Betweenness **(B):** how often a TF lies on the shortest paths between other TFs, reflecting its role as a connecting bridge. Closeness **(C):** how close a TF is to all other TFs, indicating its central position in the network. Total degree **(D):** the number of direct connections a TF has, measuring its immediate influence. Eigenvector centrality **(E):** a measure of a TF’s influence based on the importance of its neighbors. PageRank **(F):** a variant of eigenvector centrality that ranks TFs by connectivity and network flow.

Data are shown in native values or log10 scales where indicated.

Figure S9: **GnRH dependent regulatory architecture of DEGs in the *Fshb* TAD.**

**(A)** Analysis of basal (untreated) or 1h GnRHa-stimulated activity of human *Fshb* enhancers. LβT2 cells were transiently transfected with a GL4 luciferase reporter cassette under the control of each enhancer or with pGL4 empty backbone used as control. Data represent the mean +/- SEM of basal (top panel) or treated over untreated fold induction (bottom panel), normalised to pGL4. Differences for untreated cells was evaluated using a linear mixed-effects model on log10-transformed values, including experiment identity to account for experiment-to-experiment variability, followed by a Dunnett post-hoc test against pGL4 with Benjamini-Hochberg correction (N>5). For 1h stimulated cells, differences were evaluated using a one-way ANOVA on log10-transformed values, followed by a Dunnett post-hoc test versus pGL4 with Benjamini-Hochberg correction (N=3). **p < 0.01, ***p < 0.001 and **** p < 0.0001

**(B)** Temporal profiling of mRNA levels in LβT2 cells under GnRHa stimulation for *Kcna4, Dcdc5* and *Kif18a*. Transcript levels were measured by RT-qPCR in LβT2 cells either untreated or treated with 10nM GnRHa for indicated times. Data were normalised to two housekeeping genes, *Ppia* and *Rplp0* and are presented as mean +/- SEM mRNA level relative to the untreated cells. Expression changes over time were evaluated using a non-parametric Kruskal-Wallis test on normalised log-transformed values (log10), with Benjamini-Hochberg correction applied to the global p-values. *p < 0.05, **p < 0.01, ***p < 0.001 and **** p < 0.0001 (N>3 per time point).

**(C)** Analysis of CRISPRi efficiency in LβT2 cells. LβT2 cells were transiently transfected with CRISPRi targeting each of the top *Fshb* regulators: *Fos*, *Fosl1*, *Fosl2*, *Bach1*, *Bach2*, *Mef2a* or *Mef2d*. Irrelevant gRNA was used as untargeting control. Changes in relative mRNA level for each TF was assessed by RTqPCR and differences were evaluated using a Mann-Whitney test against the control. **p < 0.01 and **** p < 0.0001 (N>6).

**(D-G)** Impact of the top regulators on basal (**D**) and GnRHa-regulated expression of *Kcna4* (**E**), *Dcdc5* (**F**) and *Kif18a* (**G**). LβT2 cells were transiently transfected with CRISPRi targeting each of the top *Fshb* regulators: *Fos*, *Fosl1*, *Fosl2*, *Bach1*, *Bach2*, *Mef2a* or *Mef2d*. Irrelevant gRNA was used as untargeting control. Data are shown as mean +/- SEM of basal mRNA level (E-G top panels) or mRNA fold change (E-G bottom panels). Paired CT-CRISPRi experiments are plotted using different colours (experiments A, B and C). Statistical analysis used a linear mixed-effects model on log10-transformed induction values, including experiment identity to account for experiment-to-experiment variability, followed by a Dunnett post-hoc test versus control with Benjamini-Hochberg correction. *p < 0.05, **p < 0.01 and **** p < 0.001 (N=3, n=9).

## Fundings

This work was supported by grants from Université Paris Cité, the Centre National de la Recherche Scientifique (CNRS), and the French National Research Agency (ANR) under project FEMCYCLE (ANR-25-CE14-7932). C.L.C. received a PhD fellowship from the French Ministry of Higher Education, Research and Space, a fourth-year PhD fellowship from the Fondation pour la Recherche Médicale (FDT202404018080), and post-doctoral fellowship from the Formula inIdEx project (Université Paris Cité), part of the “Initiative d’Excellence” IdEx program (ANR-18-IDEX-0001) of the France 2030 investment plan.

## Conflicts of interest

None

## Acknowledgements

This work benefited from equipment and services provided by the Cybio and Genom’IC core facilities, the Cochin Institute, and the Animal Facility of Institut Jacques Monod. We are particularly grateful to Brigitte Izak from the Genom’IC factility for her help with the snMultiome wet-lab experiments. We also acknowledge technical support from Eloïse Airaud and Raphaël Core. We warmly thank bioinformaticians such as Dr. Sam Morabito (https://smorabit.github.io/blog/2021/velocyto/) and Dr. Ryan Mulqueen (https://github.com/mulqueenr), to name only a few, who made their codes freely available, providing invaluable assistance. Finally, we thank Dr. Mariam Okhovat from the Oregon Health and Science University for generously providing the Rat TADs file.

## Notes

### Competing Interest Statement

The authors have declared no competing interest.

## References

1. Attia, G. M., Alharbi, O. A. & Aljohani, R. M. The Impact of Irregular Menstruation on Health: A Review of the Literature. Cureus 15, e49146.

2. Sarkar, D. K., Chiappa, S. A., Fink, G. & Sherwood, N. M. Gonadotropin-releasing hormone surge in pro-oestrous rats. Nature 264, 461–463 (1976).

3. Gay, V. L., Midgley, A. R. & Niswender, G. D. Patterns of gonadotrophin secretion associated with ovulation. Fed. Proc. 29, 1880–1887 (1970).

4. Marshall, J. C. et al. Gonadotropin-releasing hormone pulses: regulators of gonadotropin synthesis and ovulatory cycles. Recent Prog. Horm. Res. 47, 155–187; discussion 188-189 (1991).

5. Fortin, J. et al. NR5A2 regulates Lhb and Fshb transcription in gonadotrope-like cells in vitro, but is dispensable for gonadotropin synthesis and fertility in vivo. PloS One 8, e59058 (2013).

6. Schultz, H. et al. ZEB1 Inhibits LHβ Subunit Transcription When Overexpressed, but Is Dispensable for LH Synthesis in Mice. Endocrinology 165, bqae116 (2024).

7. Alonso, C. A. I. et al. Activating Transcription Factor 3 Stimulates Follicle-Stimulating Hormone-β Expression In Vitro But Is Dispensable for Follicle-Stimulating Hormone Production in Murine Gonadotropes In Vivo. Endocrinology 164, bqad050 (2023).

8. Badia-I-Mompel, P. et al. Gene regulatory network inference in the era of single-cell multi-omics. Nat. Rev. Genet. 24, 739–754 (2023).

9. Besecke, L. M. et al. Pituitary follistatin regulates activin-mediated production of follicle-stimulating hormone during the rat estrous cycle. Endocrinology 138, 2841–2848 (1997).

10. Butcher, R. L., Collins, W. E. & Fugo, N. W. Plasma concentration of LH, FSH, prolactin, progesterone and estradiol-17beta throughout the 4-day estrous cycle of the rat. Endocrinology 94, 1704–1708 (1974).

11. Wall, E. G. et al. Unexpected Plasma Gonadal Steroid and Prolactin Levels Across the Mouse Estrous Cycle. Endocrinology 164, bqad070 (2023).

12. Dupon, C. & Kim, M. H. Peripheral plasma levels of testosterone, androstenedione, and oestradiol during the rat oestrous cycle. J. Endocrinol. 59, 653–654 (1973).

13. Fletcher, P. A. et al. Cell Type- and Sex-Dependent Transcriptome Profiles of Rat Anterior Pituitary Cells. Front. Endocrinol. 10, 623 (2019).

14. Cheung, L. Y. M. et al. Single-Cell RNA Sequencing Reveals Novel Markers of Male Pituitary Stem Cells and Hormone-Producing Cell Types. Endocrinology 159, 3910–3924 (2018).

15. Collado-Torres, L. et al. Flexible expressed region analysis for RNA-seq with derfinder. Nucleic Acids Res. 45, e9 (2017).

16. Hao, Y. et al. Dictionary learning for integrative, multimodal and scalable single-cell analysis. Nat. Biotechnol. 42, 293–304 (2024).

17. Stuart, T., Srivastava, A., Madad, S., Lareau, C. A. & Satija, R. Single-cell chromatin state analysis with Signac. Nat. Methods 18, 1333–1341 (2021).

18. Pulichino, A.-M. et al. Tpit determines alternate fates during pituitary cell differentiation. Genes Dev. 17, 738–747 (2003).

19. Cesnjaj, M., Catt, K. J. & Stojilkovic, S. S. Coordinate actions of calcium and protein kinase-C in the expression of primary response genes in pituitary gonadotrophs. Endocrinology 135, 692–701 (1994).

20. Ingraham, H. A. et al. The nuclear receptor steroidogenic factor 1 acts at multiple levels of the reproductive axis. Genes Dev. 8, 2302–2312 (1994).

21. Tamura, T., Yanai, H., Savitsky, D. & Taniguchi, T. The IRF family transcription factors in immunity and oncogenesis. Annu. Rev. Immunol. 26, 535–584 (2008).

22. Okhovat, M. et al. TAD evolutionary and functional characterization reveals diversity in mammalian TAD boundary properties and function. Nat. Commun. 14, 8111 (2023).

23. Langlais, D., Couture, C., Sylvain-Drolet, G. & Drouin, J. A pituitary-specific enhancer of the POMC gene with preferential activity in corticotrope cells. Mol. Endocrinol. Baltim. Md 25, 348–359 (2011).

24. Shima, Y. et al. Pituitary homeobox 2 regulates adrenal4 binding protein/steroidogenic factor-1 gene transcription in the pituitary gonadotrope through interaction with the intronic enhancer. Mol. Endocrinol. Baltim. Md 22, 1633–1646 (2008).

25. DiMattia, G. E. et al. The Pit-1 gene is regulated by distinct early and late pituitary-specific enhancers. Dev. Biol. 182, 180–190 (1997).

26. Pacini, V. et al. Identification of a pituitary ERα-activated enhancer triggering the expression of Nr5a1, the earliest gonadotrope lineage-specific transcription factor. Epigenetics Chromatin 12, 48 (2019).

27. Cheung, L. Y. M. et al. Novel Candidate Regulators and Developmental Trajectory of Pituitary Thyrotropes. Endocrinology 164, bqad076 (2023).

28. Fletcher, P. A. et al. The astroglial and stem cell functions of adult rat folliculostellate cells. Glia 71, 205–228 (2023).

29. Le Ciclé, C. et al. The Neurod1/4-Ntrk3-Src pathway regulates gonadotrope cell adhesion and motility. Cell Death Discov. 9, 327 (2023).

30. Qiao, S. et al. Molecular Plasticity of Male and Female Murine Gonadotropes Revealed by mRNA Sequencing. Endocrinology 157, 1082–1093 (2016).

31. Mattei, D., et al. Enzymatic Dissociation Induces Transcriptional and Proteotype Bias in Brain Cell Populations. Int. J. Mol. Sci. 21, 7944 (2020).

32. La Manno, G. et al. RNA velocity of single cells. Nature 560, 494–498 (2018).

33. Gallo, R. V. Pulsatile LH release during periods of low level LH secretion in the rat estrous cycle. Biol. Reprod. 24, 771–777 (1981).

34. Götz, V. et al. Ovulation is triggered by a cyclical modulation of gonadotropes into a hyperexcitable state. Cell Rep. 42, 112543 (2023).

35. Santiago-Andres, Y., Golan, M. & Fiordelisio, T. Functional Pituitary Networks in Vertebrates. Front. Endocrinol. 11, 619352 (2020).

36. Yamaguchi, M., Matsuzaki, N., Hirota, K., Miyake, A. & Tanizawa, O. Interleukin 6 possibly induced by interleukin 1 beta in the pituitary gland stimulates the release of gonadotropins and prolactin. Acta Endocrinol. (Copenh.) 122, 201–205 (1990).

37. Tolmachova, T. et al. A General Role for Rab27a in Secretory Cells. Mol. Biol. Cell 15, 332– 344 (2004).

38. MacDougall, D. D. et al. The high-affinity calcium sensor synaptotagmin-7 serves multiple roles in regulated exocytosis. J. Gen. Physiol. 150, 783–807 (2018).

39. Kennedy, M. J., Davison, I. G., Robinson, C. G. & Ehlers, M. D. Syntaxin-4 Defines a Domain for Activity-Dependent Exocytosis in Dendritic Spines. Cell 141, 524–535 (2010).

40. Nowack, A., Yao, J., Custer, K. L. & Bajjalieh, S. M. SV2 regulates neurotransmitter release via multiple mechanisms. Am. J. Physiol. - Cell Physiol. 299, C960–C967 (2010).

41. D’amico, M. A., Ghinassi, B., Izzicupo, P., Manzoli, L. & Di Baldassarre, A. Biological function and clinical relevance of chromogranin A and derived peptides. Endocr. Connect. 3, R45–R54 (2014).

42. Geerts, C. J. et al. Tomosyn-2 is required for normal motor performance in mice and sustains neurotransmission at motor endplates. Brain Struct. Funct. 220, 1971–1982 (2015).

43. Michalski, N. et al. Otoferlin acts as a Ca2+ sensor for vesicle fusion and vesicle pool replenishment at auditory hair cell ribbon synapses. eLife 6, e31013.

44. Ji, L., Wu, H.-T., Qin, X.-Y. & Lan, R. Dissecting carboxypeptidase E: properties, functions and pathophysiological roles in disease. Endocr. Connect. 6, R18–R38 (2017).

45. Toledo, P. L. et al. Condensation of the β-cell secretory granule luminal cargoes pro/insulin and ICA512 RESP18 homology domain. Protein Sci. Publ. Protein Soc. 32, e4649 (2023).

46. Steegmaier, M., Klumperman, J., Foletti, D. L., Yoo, J.-S. & Scheller, R. H. Vesicle-associated Membrane Protein 4 is Implicated in Trans-Golgi Network Vesicle Trafficking. Mol. Biol. Cell 10, 1957–1972 (1999).

47. Stout, K. A., Dunn, A. R., Hoffman, C. & Miller, G. W. The Synaptic Vesicle Glycoprotein 2: Structure, Function, and Disease Relevance. ACS Chem. Neurosci. 10, 3927–3938 (2019).

48. Watson, P., Townley, A. K., Koka, P., Palmer, K. J. & Stephens, D. J. Sec16 defines endoplasmic reticulum exit sites and is required for secretory cargo export in mammalian cells. Traffic Cph. Den. 7, 1678–1687 (2006).

49. Yamamoto, K. et al. The KDEL receptor mediates a retrieval mechanism that contributes to quality control at the endoplasmic reticulum. EMBO J. 20, 3082–3091 (2001).

50. Roberts, B. S. & Satpute-Krishnan, P. The many hats of transmembrane emp24 domain protein TMED9 in secretory pathway homeostasis. Front. Cell Dev. Biol. 10, 1096899 (2023).

51. Bailey Blackburn, J., Pokrovskaya, I., Fisher, P., Ungar, D. & Lupashin, V. V.. COG Complex Complexities: Detailed Characterization of a Complete Set of HEK293T Cells Lacking Individual COG Subunits. Front. Cell Dev. Biol. 4, 23 (2016).

52. Dejgaard, S. Y. & Presley, J. F. Arfs on the Golgi: four conductors, one orchestra. Front. Mol. Biosci. 12, 1612531 (2025).

53. Yuan, W. & Song, C. The Emerging Role of Rab5 in Membrane Receptor Trafficking and Signaling Pathways. Biochem. Res. Int. 2020, 4186308 (2020).

54. Rosa-Ferreira, C., Christis, C., Torres, I. L. & Munro, S. The small G protein Arl5 contributes to endosome-to-Golgi traffic by aiding the recruitment of the GARP complex to the Golgi. Biol. Open 4, 474–481 (2015).

55. Jackson, T. R. et al. Acaps Are Arf6 Gtpase-Activating Proteins That Function in the Cell Periphery. J. Cell Biol. 151, 627–638 (2000).

56. Wang, H. H., Biunno, I., Sun, S. & Qi, L. SEL1L-HRD1-mediated ERAD in mammals. Nat. Cell Biol. 27, 1063–1073 (2025).

57. Roth, J. & Zuber, C. Quality control of glycoprotein folding and ERAD: the role of N-glycan handling, EDEM1 and OS-9. Histochem. Cell Biol. 147, 269–284 (2017).

58. Hayano, T. & Kikuchi, M. Molecular cloning of the cDNA encoding a novel protein disulfide isomerase-related protein (PDIR). FEBS Lett. 372, 210–214 (1995).

59. Senesi, S. et al. Hexose-6-phosphate dehydrogenase in the endoplasmic reticulum. Biol. Chem. 391, 1–8 (2010).

60. Kang, R., Zeh, H. J., Lotze, M. T. & Tang, D. The Beclin 1 network regulates autophagy and apoptosis. Cell Death Differ. 18, 571–580 (2011).

61. Castle, J. D., Guo, Z. & Liu, L. Function of the t-SNARE SNAP-23 and secretory carrier membrane proteins (SCAMPs) in exocytosis in mast cells. Mol. Immunol. 38, 1337–1340 (2002).

62. Soo Hoo, L., et al. The SNARE Protein Syntaxin 3 Confers Specificity for Polarized Axonal Trafficking in Neurons. PLoS ONE 11, e0163671 (2016).

63. Sukriti, S., Tauseef, M., Yazbeck, P. & Mehta, D. Mechanisms regulating endothelial permeability. Pulm. Circ. 4, 535–551 (2014).

64. Midwood, K. S., Hussenet, T., Langlois, B. & Orend, G. Advances in tenascin-C biology. Cell. Mol. Life Sci. CMLS 68, 3175–3199 (2011).

65. Namba, T. et al. Collagen 17A1 in the Urothelium Regulates Epithelial Cell Integrity and Local Immunologic Responses in Obstructive Uropathy. Am. J. Pathol. 194, 1550–1570 (2024).

66. Ganapathy, A., Chen, Y., Bakthavachalam, V. & George, A. DMP1-mediated transformation of DPSCs to CD31+/CD144+ cells demonstrate endothelial phenotype both in vitro and in vivo. Front. Cell Dev. Biol. 13, 1630129 (2025).

67. Zmeili, S. M. et al. Alpha and luteinizing hormone beta subunit messenger ribonucleic acids during the rat estrous cycle. Endocrinology 119, 1867–1869 (1986).

68. Shupnik, M. A., Gharib, S. D. & Chin, W. W. Divergent effects of estradiol on gonadotropin gene transcription in pituitary fragments. Mol. Endocrinol. Baltim. Md 3, 474–480 (1989).

69. Refael, T. et al. An i-motif-regulated enhancer, eRNA and adjacent lncRNA affect Lhb expression through distinct mechanisms in a sex-specific context. Cell. Mol. Life Sci. CMLS 81, 361 (2024).

70. Arriola, D. J., Mayo, S. L., Skarra, D. V., Benson, C. A. & Thackray, V. G. FOXO1 transcription factor inhibits luteinizing hormone β gene expression in pituitary gonadotrope cells. J. Biol. Chem. 287, 33424–33435 (2012).

71. McNeilly, A. S., Crawford, J. L., Taragnat, C., Nicol, L. & McNeilly, J. R. The differential secretion of FSH and LH: regulation through genes, feedback and packaging. Reprod. Camb. Engl. Suppl. 61, 463–476 (2003).

72. Haisenleder, D. J., Dalkin, A. C., Ortolano, G. A., Marshall, J. C. & Shupnik, M. A. A pulsatile gonadotropin-releasing hormone stimulus is required to increase transcription of the gonadotropin subunit genes: evidence for differential regulation of transcription by pulse frequency in vivo. Endocrinology 128, 509–517 (1991).

73. Burger, L. L., Haisenleder, D. J., Dalkin, A. C. & Marshall, J. C. Regulation of gonadotropin subunit gene transcription. J. Mol. Endocrinol. 33, 559–584 (2004).

74. Bohaczuk, S. C., Thackray, V. G., Shen, J., Skowronska-Krawczyk, D. & Mellon, P. L. FSHB Transcription is Regulated by a Novel 5’ Distal Enhancer With a Fertility-Associated Single Nucleotide Polymorphism. Endocrinology 162, bqaa181 (2021).

75. Benayoun, B. A., Auer, J., Caburet, S. & Veitia, R. A. The post-translational modification profile of the forkhead transcription factor FOXL2 suggests the existence of parallel processive/concerted modification pathways. Proteomics 8, 3118–3123 (2008).

76. Xu, P., Lin, X. & Feng, X.-H. Posttranslational Regulation of Smads. Cold Spring Harb. Perspect. Biol. 8, a022087 (2016).

77. Astapova, O., Seger, C. & Hammes, S. R. Ligand Binding Prolongs Androgen Receptor Protein Half-Life by Reducing its Degradation. J. Endocr. Soc. 5, bvab035 (2021).

78. Huang, P. et al. SOX4 facilitates PGR protein stability and FOXO1 expression conducive for human endometrial decidualization. eLife 11, e72073 (2022).

79. Park, C. H., Skarra, D. V., Rivera, A. J., Arriola, D. J. & Thackray, V. G. Constitutively active FOXO1 diminishes activin induction of Fshb transcription in immortalized gonadotropes. PloS One 9, e113839 (2014).

80. Skarra, D. V., Arriola, D. J., Benson, C. A. & Thackray, V. G. Forkhead box O1 is a repressor of basal and GnRH-induced Fshb transcription in gonadotropes. Mol. Endocrinol. Baltim. Md 27, 1825–1839 (2013).

81. Mistry, D. S. et al. Gonadotropin-releasing hormone pulse sensitivity of follicle-stimulating hormone-beta gene is mediated by differential expression of positive regulatory activator protein 1 factors and corepressors SKIL and TGIF1. Mol. Endocrinol. Baltim. Md 25, 1387– 1403 (2011).

82. Brûlé, E. et al. TGFBR3L is an inhibin B co-receptor that regulates female fertility. Sci. Adv. 7, eabl4391 (2021).

83. Bauer-Dantoin, A. C., Hollenberg, A. N. & Jameson, J. L. Dynamic regulation of gonadotropin-releasing hormone receptor mRNA levels in the anterior pituitary gland during the rat estrous cycle. Endocrinology 133, 1911–1914 (1993).

84. Janjic, M. M. et al. Divergent expression patterns of pituitary gonadotropin subunit and GnRH receptor genes to continuous GnRH in vitro and in vivo. Sci. Rep. 9, 20098 (2019).

85. Bjelobaba, I. et al. The relationship between basal and regulated Gnrhr expression in rodent pituitary gonadotrophs. Mol. Cell. Endocrinol. 437, 302–311 (2016).

86. Schirman-Hildesheim, T. D., Ben-Aroya, N. & Koch, Y. Daily GnRH and GnRH-receptor mRNA expression in the ovariectomized and intact rat. Mol. Cell. Endocrinol. 252, 120–125 (2006).

87. Odle, A. K., MacNicol, M. C., Childs, G. V. & MacNicol, A. M. Post-Transcriptional Regulation of Gnrhr: A Checkpoint for Metabolic Control of Female Reproduction. Int. J. Mol. Sci. 22, 3312 (2021).

88. Odle, A. K. et al. Association of Gnrhr mRNA With the Stem Cell Determinant Musashi: A Mechanism for Leptin-Mediated Modulation of GnRHR Expression. Endocrinology 159, 883– 894 (2018).

89. Janjic, M. M., Stojilkovic, S. S. & Bjelobaba, I. Intrinsic and Regulated Gonadotropin-Releasing Hormone Receptor Gene Transcription in Mammalian Pituitary Gonadotrophs. Front. Endocrinol. 8, 221 (2017).

90. Schang, A. Inside and outside the pituitary: comparative analysis of Gnrhr expression provides insight into the mechanisms underlying the evolution of gene expression. J. Neuroendocrinol. 27, 177–186 (2015).

91. Schang, A.-L. et al. GATA2-induced silencing and LIM-homeodomain protein-induced activation are mediated by a bi-functional response element in the rat GnRH receptor gene. Mol. Endocrinol. Baltim. Md 27, 74–91 (2013).

92. Li, Z., Nagai, J. S., Kuppe, C., Kramann, R. & Costa, I. G. scMEGA: single-cell multi-omic enhancer-based gene regulatory network inference. Bioinforma. Adv. 3, vbad003 (2023).

93. Gieske, M. C. et al. Pituitary gonadotroph estrogen receptor-alpha is necessary for fertility in females. Endocrinology 149, 20–27 (2008).

94. Zhao, L., Bakke, M. & Parker, K. L. Pituitary-specific knockout of steroidogenic factor 1. Mol. Cell. Endocrinol. 185, 27–32 (2001).

95. Tse, J. et al. Basic-Leucine-Zipper Transcription Factors Regulate Selective Molecular Phenotypes in Regulatory T Cells During IL-2-Induced Activation. 2025.02.25.638325 Preprint at 10.1101/2025.02.25.638325 (2025).

96. Borrelli, E., Nestler, E. J., Allis, C. D. & Sassone-Corsi, P. Decoding the epigenetic language of neuronal plasticity. Neuron 60, 961–974 (2008).

97. Ostuni, R. et al. Latent enhancers activated by stimulation in differentiated cells. Cell 152, 157–171 (2013).

98. Goldstein, I. et al. Transcription factor assisted loading and enhancer dynamics dictate the hepatic fasting response. Genome Res. 27, 427–439 (2017).

99. Williams, K. et al. Epigenetic rewiring of skeletal muscle enhancers after exercise training supports a role in whole-body function and human health. Mol. Metab. 53, 101290 (2021).

100. Ghaffarizadeh, A., Flann, N. S. & Podgorski, G. J. Multistable switches and their role in cellular differentiation networks. BMC Bioinformatics 15 **Suppl 7**, S7 (2014).

101. Okawa, S., Nicklas, S., Zickenrott, S., Schwamborn, J. C. & Del Sol, A. A Generalized Gene-Regulatory Network Model of Stem Cell Differentiation for Predicting Lineage Specifiers. Stem Cell Rep. 7, 307–315 (2016).

102. Terashima, R., Laoharatchatathanin, T., Kurusu, S. & Kawaminami, M. Sequential preovulatory expression of a gonadotropin-releasing hormone-inducible gene, Nr4a3, and its suppressor Anxa5 in the pituitary gland of female rats. J. Reprod. Dev. 67, 217–221 (2021).

103. Haisenleder, D. J. et al. LH subunit mRNA concentrations during LH surge in ovariectomized estradiol-replaced rats. Am. J. Physiol. 254, E99–103 (1988).

104. Brooks, J. et al. Cloning and sequencing of the sheep pituitary gonadotropin-releasing hormone receptor and changes in expression of its mRNA during the estrous cycle. Mol. Cell. Endocrinol. 94, R23–27 (1993).

105. Currie, R. J. & McNeilly, A. S. Mobilization of LH secretory granules in gonadotrophs in relation to gene expression, synthesis and secretion of LH during the preovulatory phase of the sheep oestrous cycle. J. Endocrinol. 147, 259–270 (1995).

106. Thompson, I. R. & Kaiser, U. B. GnRH pulse frequency-dependent differential regulation of LH and FSH gene expression. Mol. Cell. Endocrinol. 385, 28–35 (2014).

107. Liu, Y. et al. In-vivo properties and functional subtypes of gonadotropin-releasing hormone neurons. iScience 28, 112513 (2025).

108. Garrel, G. et al. Sustained gonadotropin-releasing hormone stimulation mobilizes the cAMP/PKA pathway to induce nitric oxide synthase type 1 expression in rat pituitary cells in vitro and in vivo at proestrus. Biol. Reprod. 82, 1170–1179 (2010).

109. Larivière, S. et al. Gonadotropin-releasing hormone couples to 3’,5’-cyclic adenosine-5’-monophosphate pathway through novel protein kinase Cdelta and -epsilon in LbetaT2 gonadotrope cells. Endocrinology 148, 1099–1107 (2007).

110. Lozach, A., Garrel, G., Lerrant, Y., Bérault, A. & Counis, R. GnRH-dependent up-regulation of nitric oxide synthase I level in pituitary gonadotrophs mediates cGMP elevation during rat proestrus. Mol. Cell. Endocrinol. 143, 43–51 (1998).

111. Stamatiades, G. A. et al. Deletion of Gαq/11 or Gαs Proteins in Gonadotropes Differentially Affects Gonadotropin Production and Secretion in Mice. Endocrinology 163, bqab247 (2022).

112. McArdle, C. A., Franklin, J., Green, L. & Hislop, J. N. Signalling, cycling and desensitisation of gonadotrophin-releasing hormone receptors. J. Endocrinol. 173, 1–11 (2002).

113. Tsutsumi, R., Mistry, D. & Webster, N. J. G. Signaling responses to pulsatile gonadotropin-releasing hormone in LbetaT2 gonadotrope cells. J. Biol. Chem. 285, 20262–20272 (2010).

114. Cohen-Tannoudji, J., Avet, C., Garrel, G., Counis, R. & Simon, V. Decoding high Gonadotropin-releasing hormone pulsatility: a role for GnRH receptor coupling to the cAMP pathway? Front. Endocrinol. 3, 107 (2012).

115. Ongaro, L. et al. Muscle-derived myostatin is a major endocrine driver of follicle-stimulating hormone synthesis. Science 387, 329–336 (2025).

116. Lin, Y.-F. et al. Loss of Inhibin Negative Feedback to Pituitary Gonadotropes Leads to Enhanced Ovulation but Pregnancy Failure in Mice. Endocrinology 166, bqaf142 (2025).

117. Bohaczuk, S. C., Cassin, J., Slaiwa, T. I., Thackray, V. G. & Mellon, P. L. Distal Enhancer Potentiates Activin- and GnRH-Induced Transcription of FSHB. Endocrinology 162, bqab069 (2021).

118. Bohaczuk, S. C. et al. A Point Mutation in an Otherwise Dispensable Upstream Fshb Enhancer Moderately Impairs Fertility in Female Mice. Endocrinology 166, bqaf073 (2025).

119. Wang, B.-B. et al. Activin A promotes bovine pituitary FSH synthesis and secretion through the cooperative action of the SMAD signaling pathway and FOXO3†. Biol. Reprod. ioaf179 (2025) doi:10.1093/biolre/ioaf179.

120. Wang, Y. et al. Activator protein-1 and smad proteins synergistically regulate human follicle-stimulating hormone beta-promoter activity. Endocrinology 149, 5577–5591 (2008).

121. Madrigal, P. & Alasoo, K. AP-1 Takes Centre Stage in Enhancer Chromatin Dynamics. Trends Cell Biol. 28, 509–511 (2018).

122. Zhang, H. et al. Fosl2 facilitates chromatin accessibility to determine developmental events during follicular maturation. Nat. Commun. 16, 8955 (2025).

123. Jang, E. et al. Bach2 represses the AP-1-driven induction of interleukin-2 gene transcription in CD4+ T cells. BMB Rep. 50, 472–477 (2017).

124. Lim, S. et al. Distinct mechanisms involving diverse histone deacetylases repress expression of the two gonadotropin beta-subunit genes in immature gonadotropes, and their actions are overcome by gonadotropin-releasing hormone. Mol. Cell. Biol. 27, 4105–4120 (2007).

125. Ichihara, M., Kamiya, T., Hara, H. & Adachi, T. The MEF2A and MEF2D function as scaffold proteins that interact with HDAC1 or p300 in SOD3 expression in THP-1 cells. Free Radic. Res. 52, 799–807 (2018).

126. Choi, S. G. et al. Growth differentiation factor 9 (GDF9) forms an incoherent feed-forward loop modulating follicle-stimulating hormone β-subunit (FSHβ) gene expression. J. Biol. Chem. 289, 16164–16175 (2014).

127. Bochimoto, H. et al. Sustained treatment with a GnRH agonist (leuprorelin) affects the ultrastructural characteristics of membranous organelles in male rat pituitary gonadotropes. Arch. Histol. Cytol. 74, 41–57 (2013).

128. Koga, D., Bochimoto, H., Kusumi, S., Ushiki, T. & Watanabe, T. Changes in the three-dimensional ultrastructure of membranous organelles in male rat pituitary gonadotropes after castration. Biomed. Res. Tokyo Jpn. 38, 1–18 (2017).

129. Choi, S. G. et al. Characterization of Gonadotrope Secretoproteome Identifies Neurosecretory Protein VGF-derived Peptide Suppression of Follicle-stimulating Hormone Gene Expression. J. Biol. Chem. 291, 21322–21334 (2016).

130. Yin, P. & Arita, J. Differential regulation of prolactin release and lactotrope proliferation during pregnancy, lactation and the estrous cycle. Neuroendocrinology 72, 72–79 (2000).

131. Hashi, A. et al. Estradiol-induced diurnal changes in lactotroph proliferation and their hypothalamic regulation in ovariectomized rats. Endocrinology 137, 3246–3252 (1996).

132. Qiao, S. et al. Intra-pituitary follicle-stimulating hormone signaling regulates hepatic lipid metabolism in mice. Nat. Commun. 14, 1098 (2023).

133. Giton, F. et al. Sex Steroids, Precursors, and Metabolite Deficiencies in Men With Isolated Hypogonadotropic Hypogonadism and Panhypopituitarism: A GCMS-Based Comparative Study. J. Clin. Endocrinol. Metab. 100, E292–E296 (2015).

134. Dainat, J., et al. NBISweden/AGAT: AGAT v1.5.1. Zenodo 10.5281/zenodo.16317950 (2025).

135. Young, M. D. & Behjati, S. SoupX removes ambient RNA contamination from droplet-based single-cell RNA sequencing data. GigaScience 9, giaa151 (2020).

136. Perez, G. et al. The UCSC Genome Browser database: 2025 update. Nucleic Acids Res. 53, D1243–D1249 (2025).

137. Bastian, M., Heymann, S. & Jacomy, M. Gephi: An Open Source Software for Exploring and Manipulating Networks. Proc. Int. AAAI Conf. Web Soc. Media 3, 361–362 (2009).

138. Kent, W. J. BLAT--the BLAST-like alignment tool. Genome Res. 12, 656–664 (2002).

139. Katoh, K., Misawa, K., Kuma, K. & Miyata, T. MAFFT: a novel method for rapid multiple sequence alignment based on fast Fourier transform. Nucleic Acids Res. 30, 3059–3066 (2002).

140. Bailey, T. L. et al. MEME SUITE: tools for motif discovery and searching. Nucleic Acids Res. 37, W202–208 (2009).

141. Gupta, S., Stamatoyannopoulos, J. A., Bailey, T. L. & Noble, W. S. Quantifying similarity between motifs. Genome Biol. 8, R24 (2007).

142. Horlbeck, M. A. et al. Compact and highly active next-generation libraries for CRISPR-mediated gene repression and activation. eLife 5, e19760 (2016).

