## Supplementary Figure S1 for "Decoding the pituitary gonadotrope regulatory architecture governing the preovulatory surge *in vivo*"

### Fig. S1

**A**

Representative images of vaginal smears used to characterise female sexual cycle stages

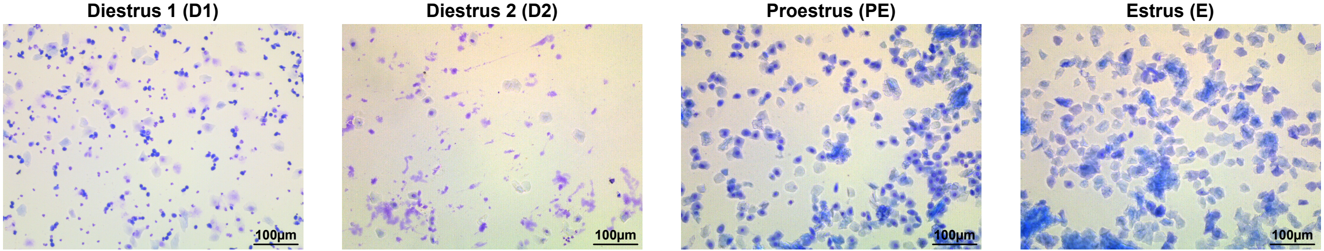


**B**

Sexual cycle progression over 15 days of the individuals selected for snMultiome analysis

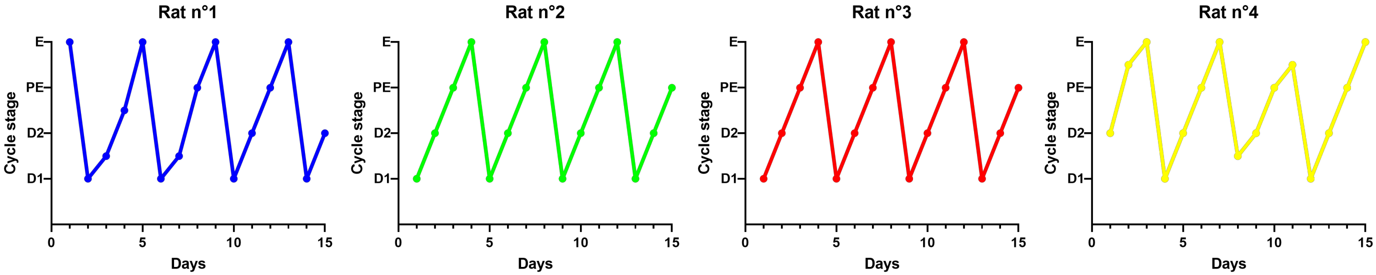
