## Supplementary figures and images for "Decoding the pituitary gonadotrope regulatory architecture governing the preovulatory surge *in vivo*"

### Supplementary Figure S2

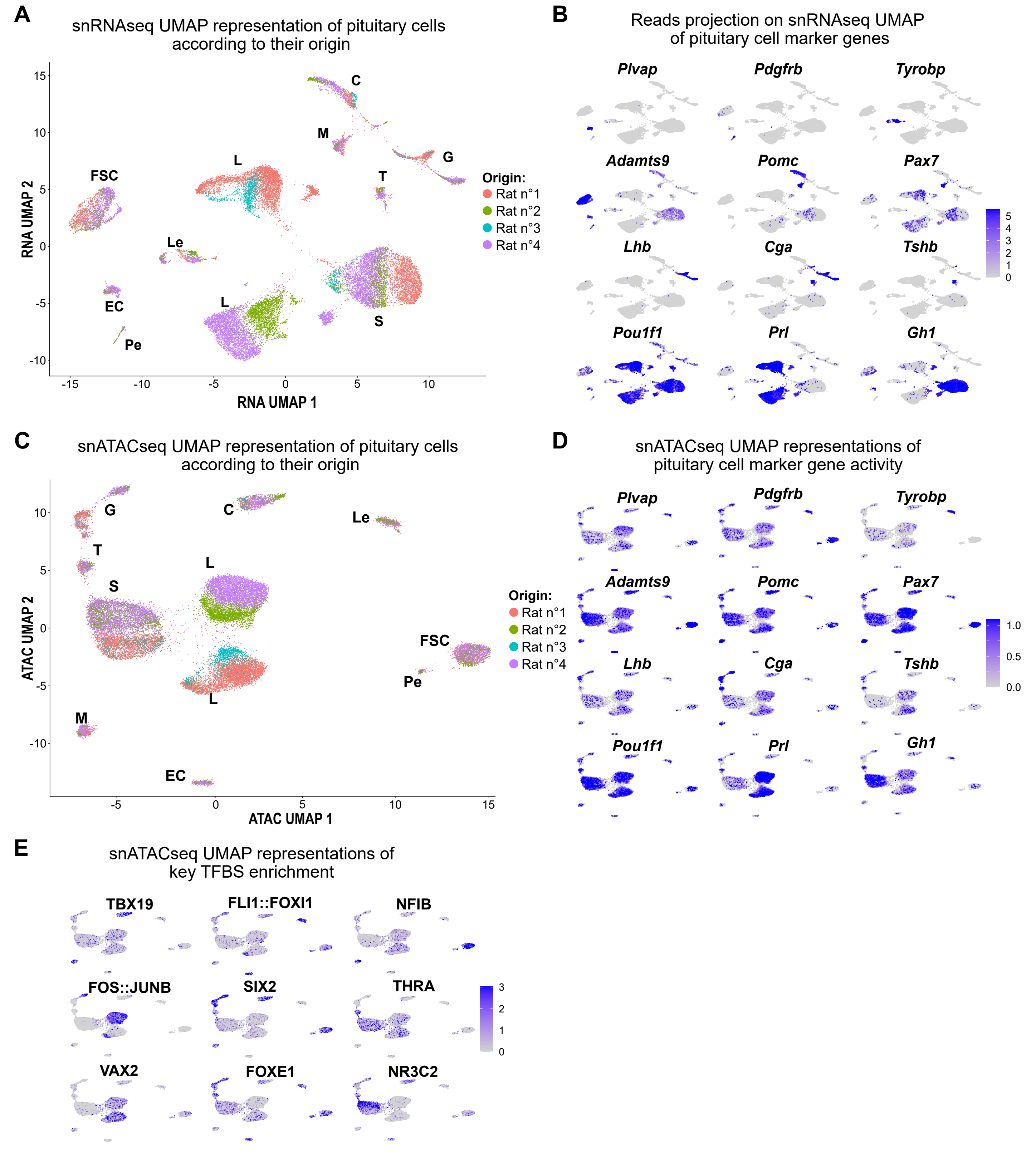

### Supplementary Figure S3.1

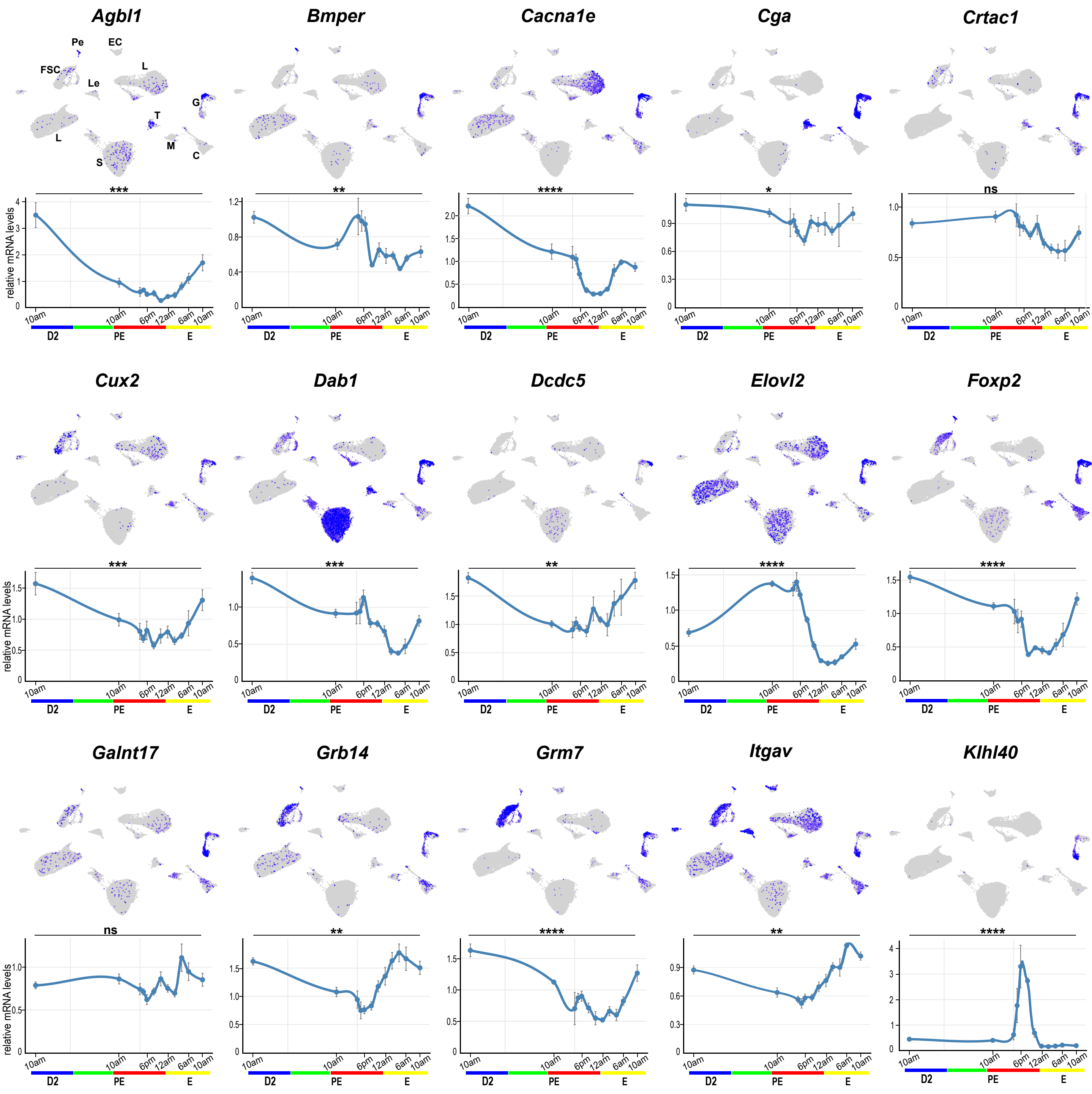

### Supplementary Figure S3.2

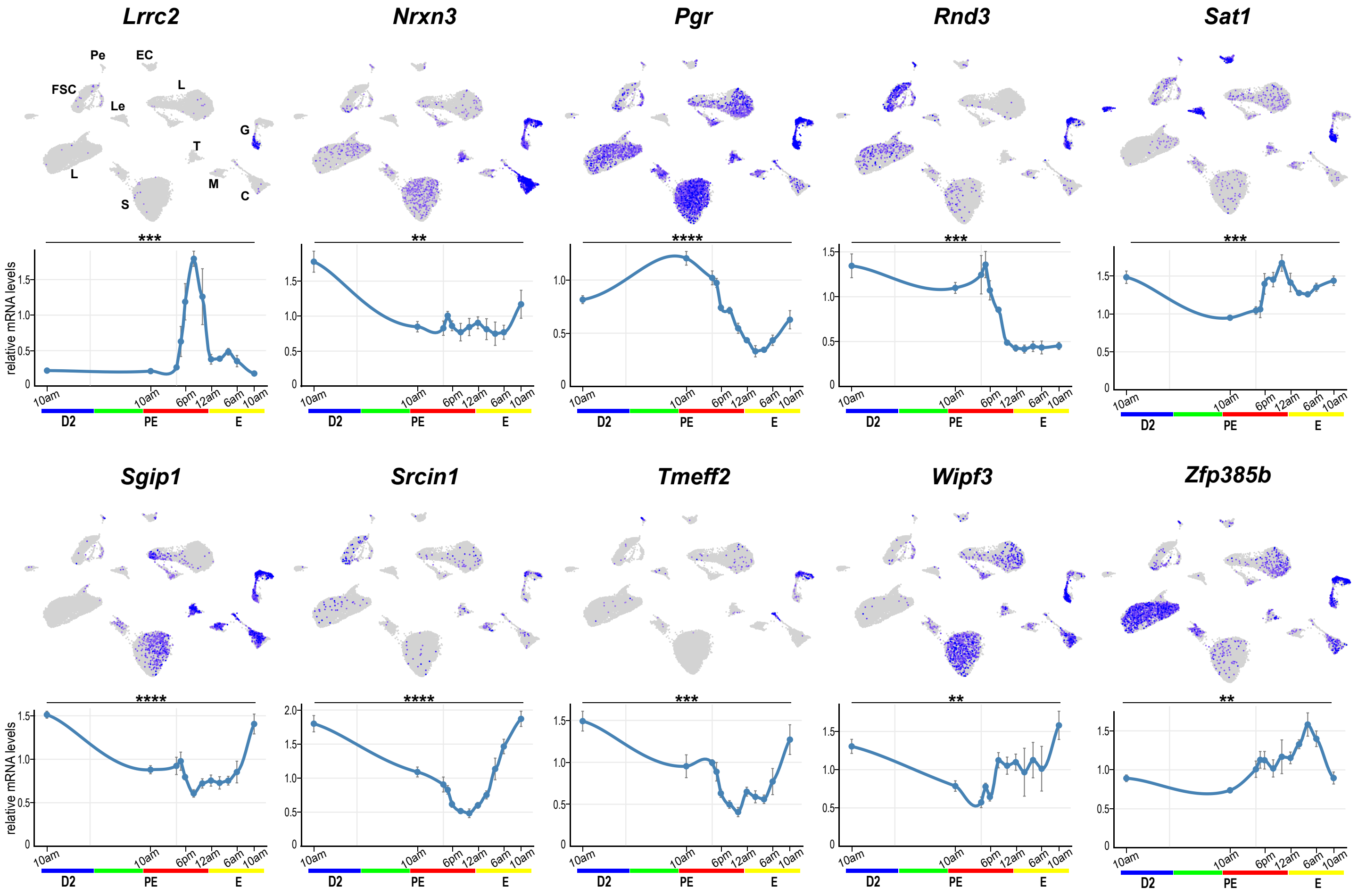

### Supplementary Figure S3.3

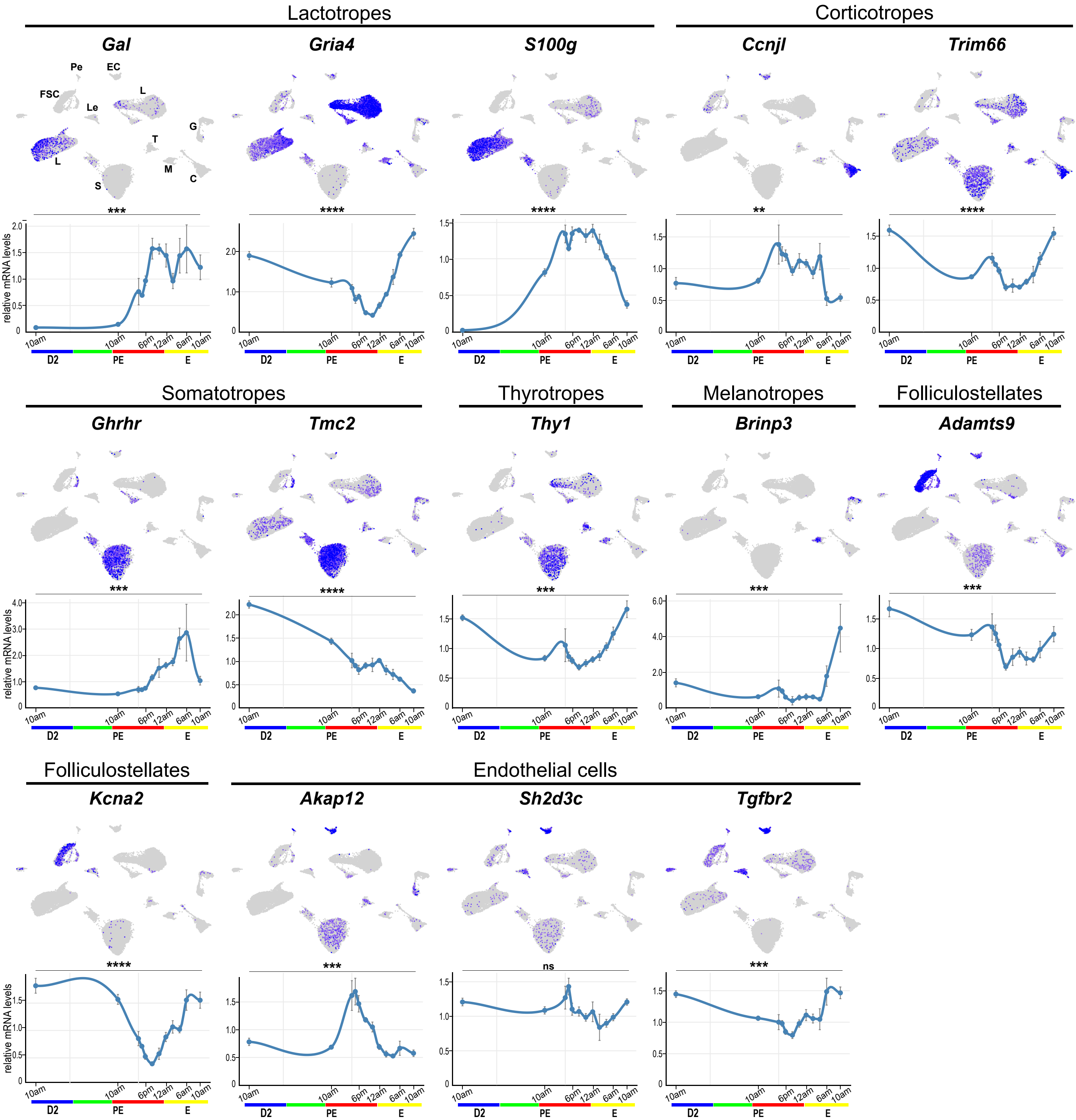

### Supplementary Figure S4

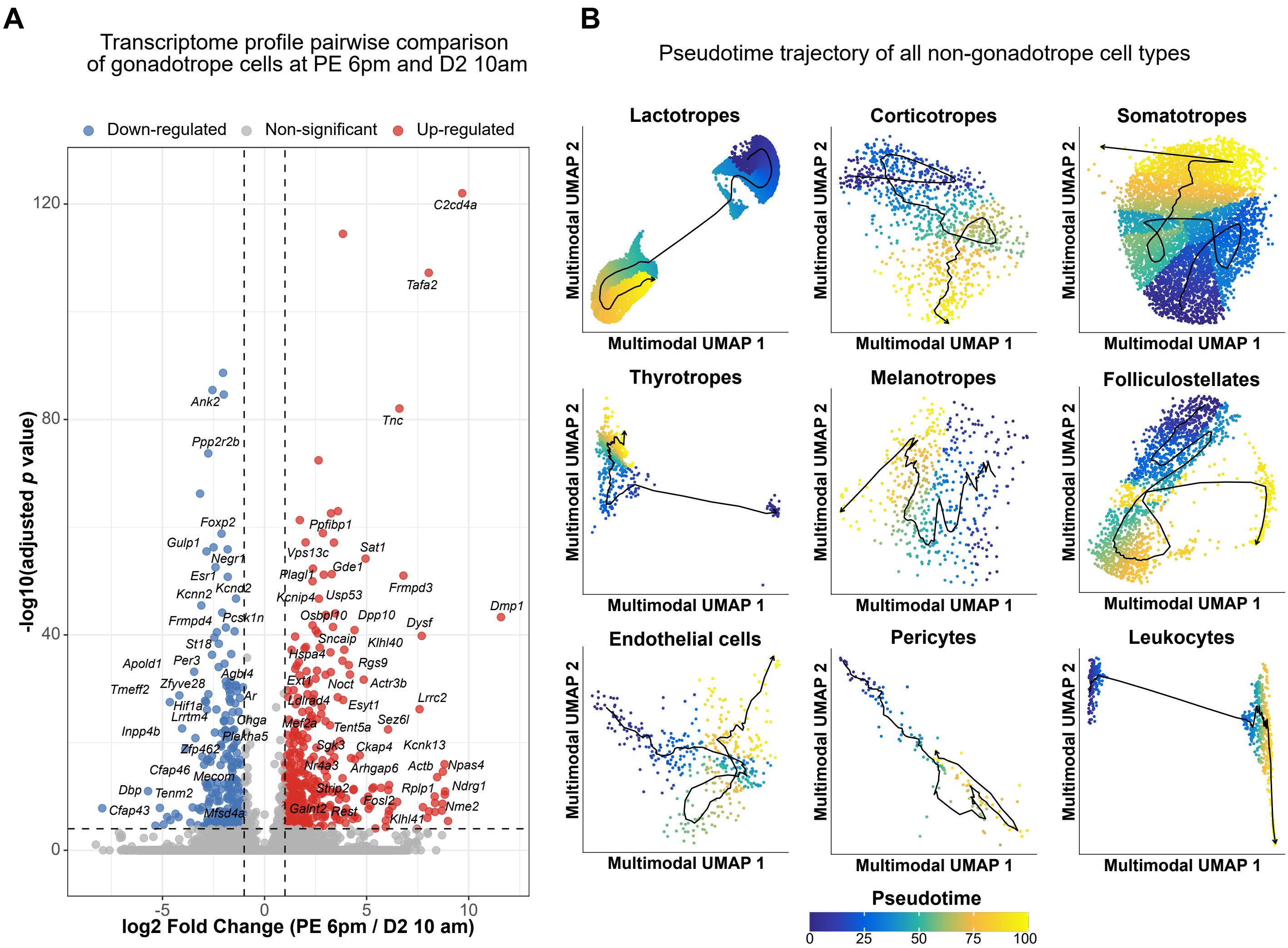

### Supplementary Figure S5

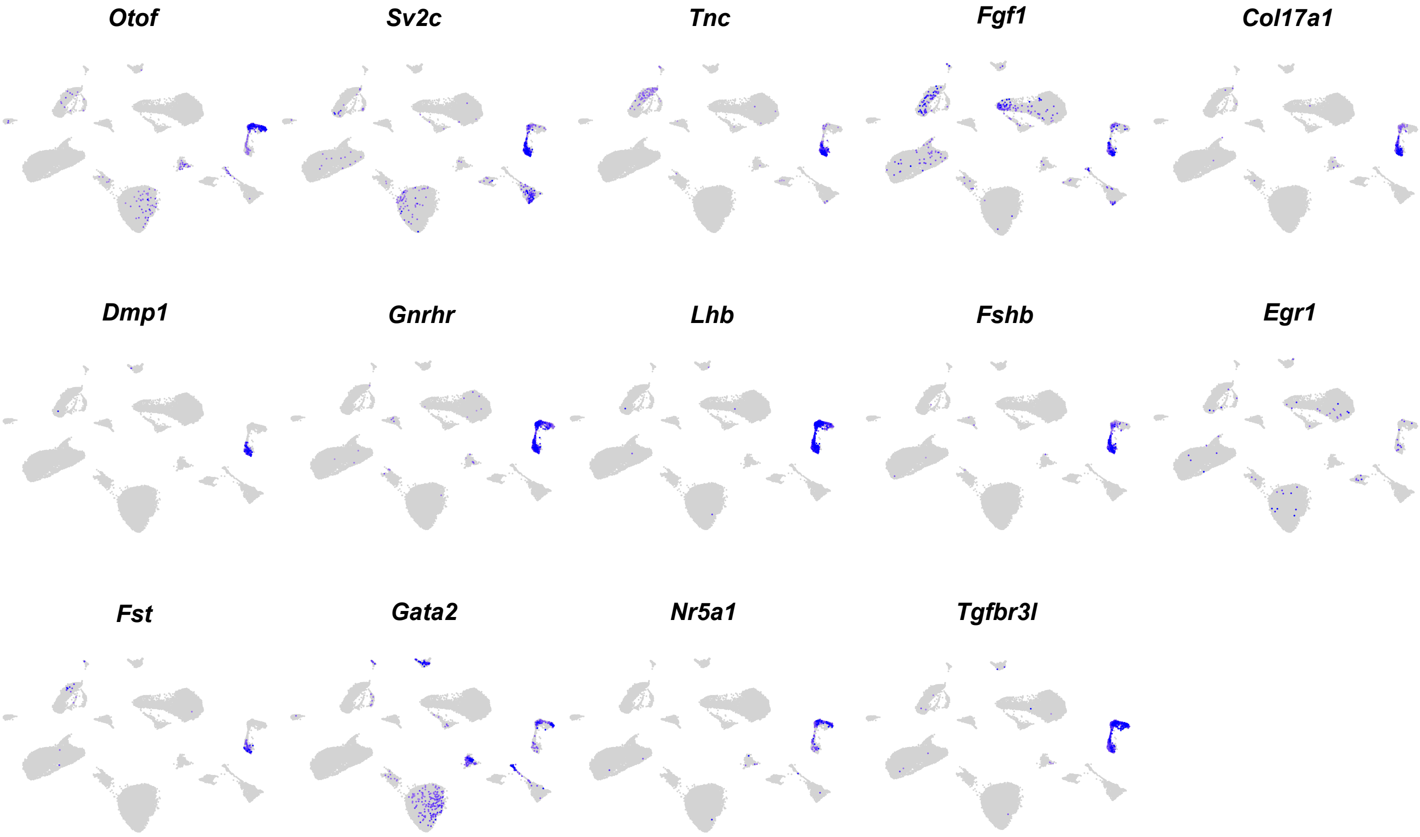

### Supplementary Figure S6

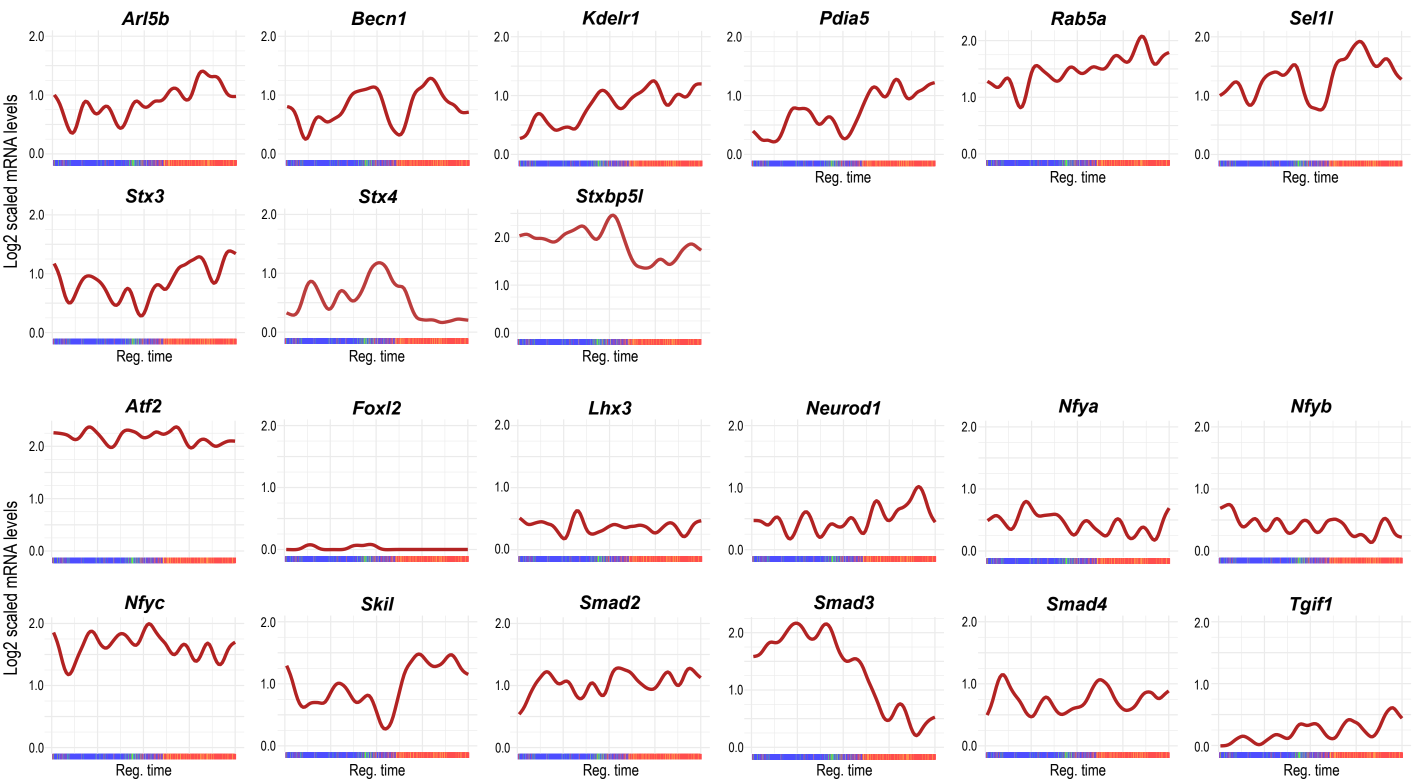

### Supplementary Figure S7.1

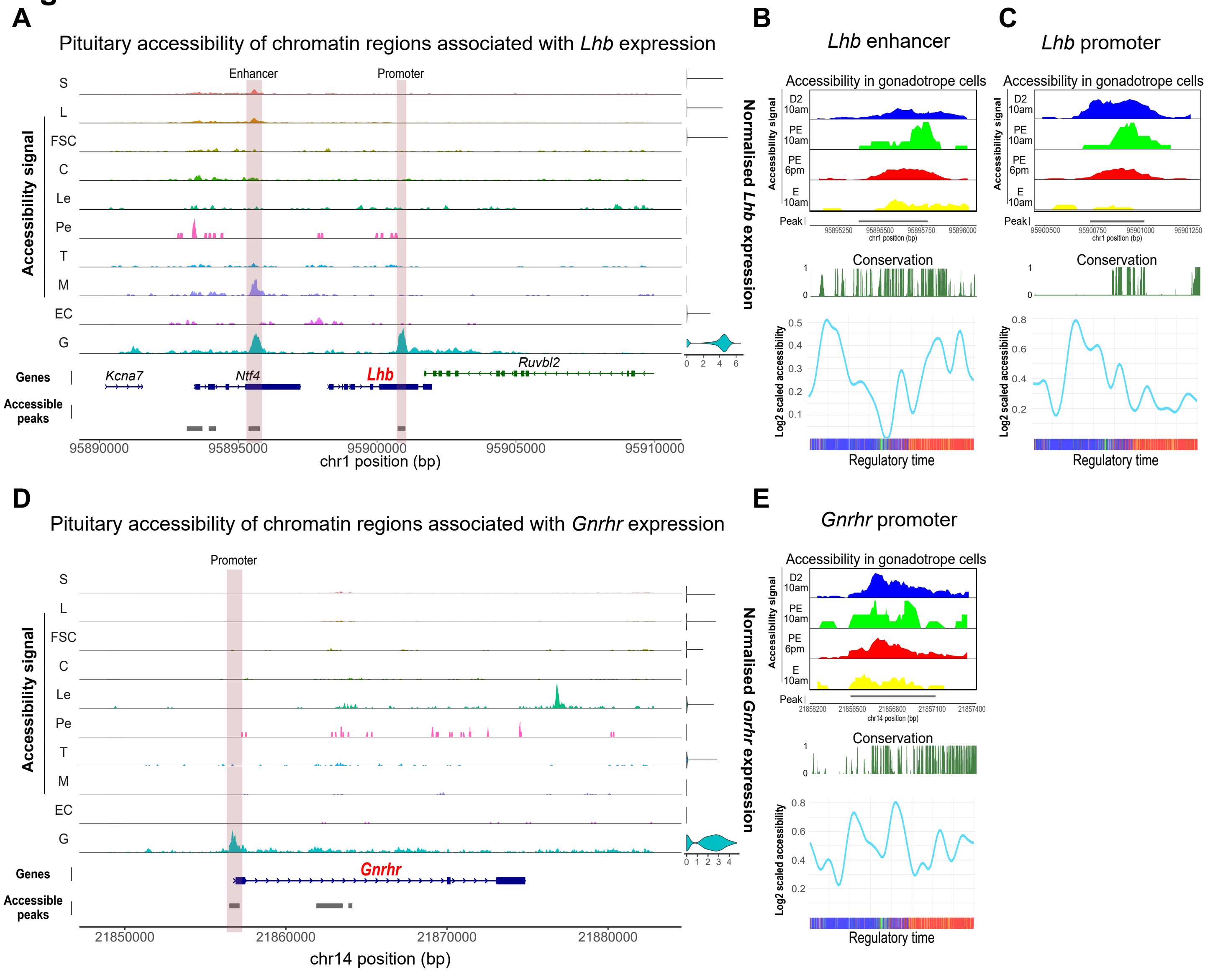

### Supplementary Figure S7.2

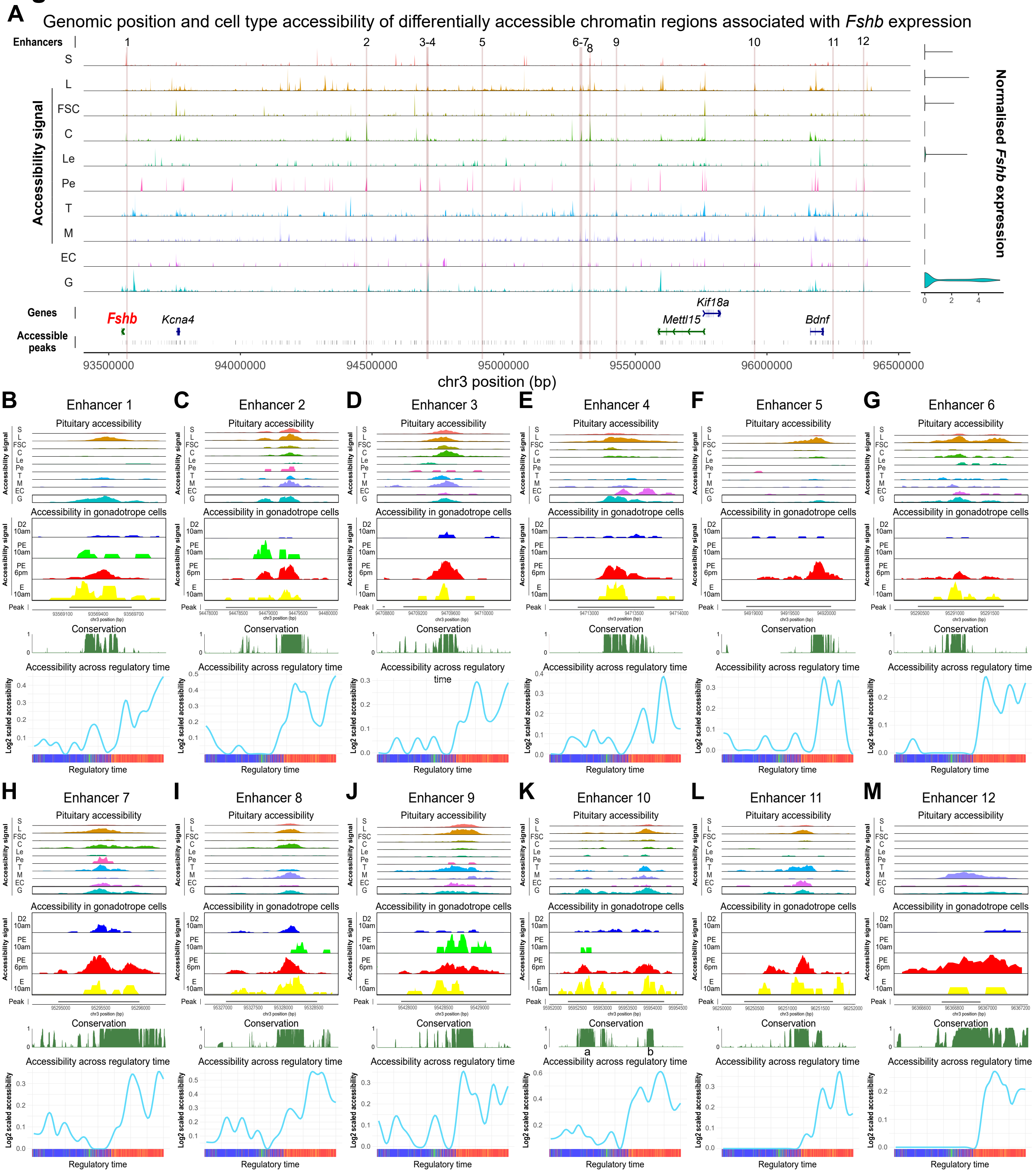

### Supplementary Figure S8

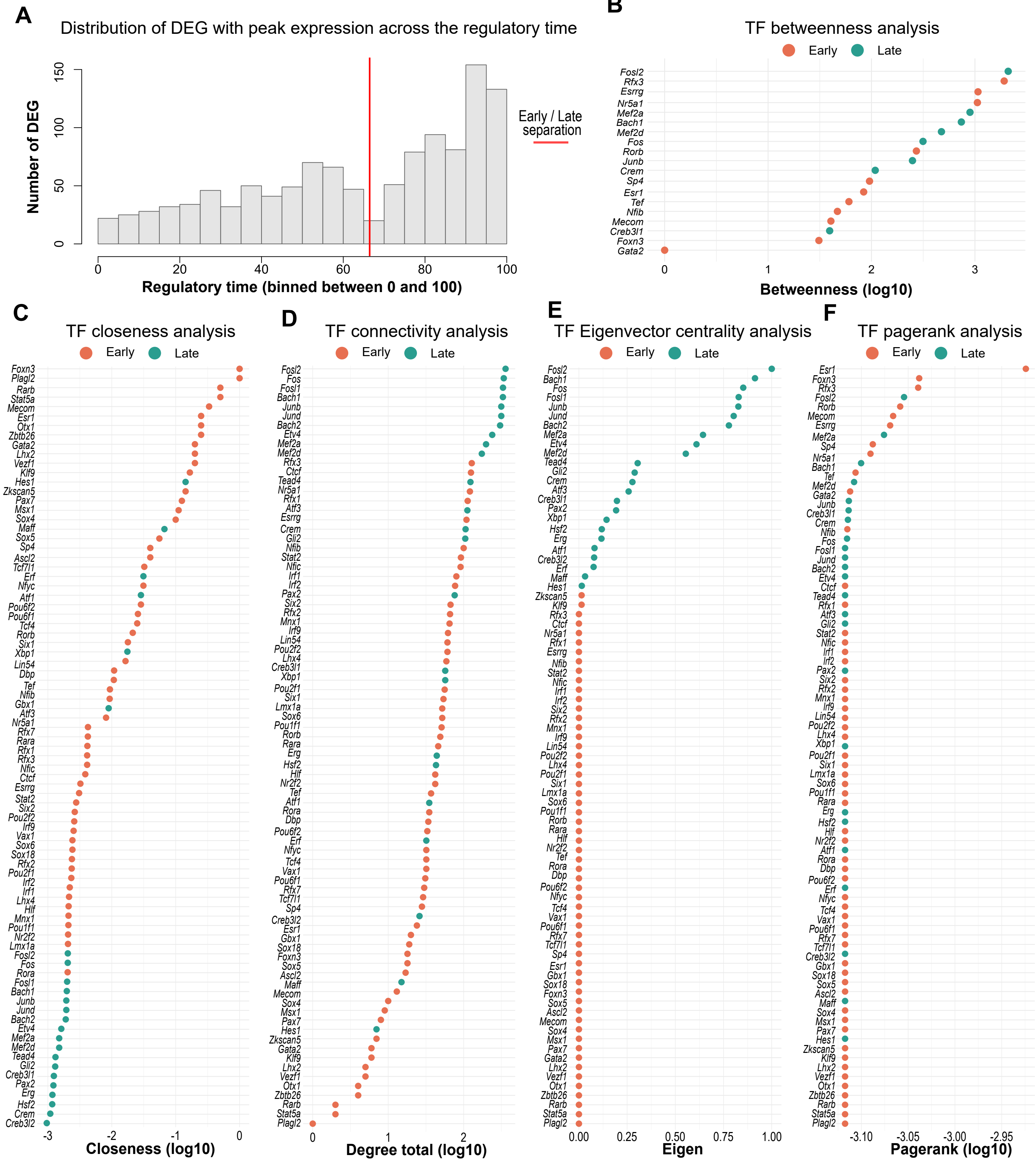

### Supplementary Figure S9

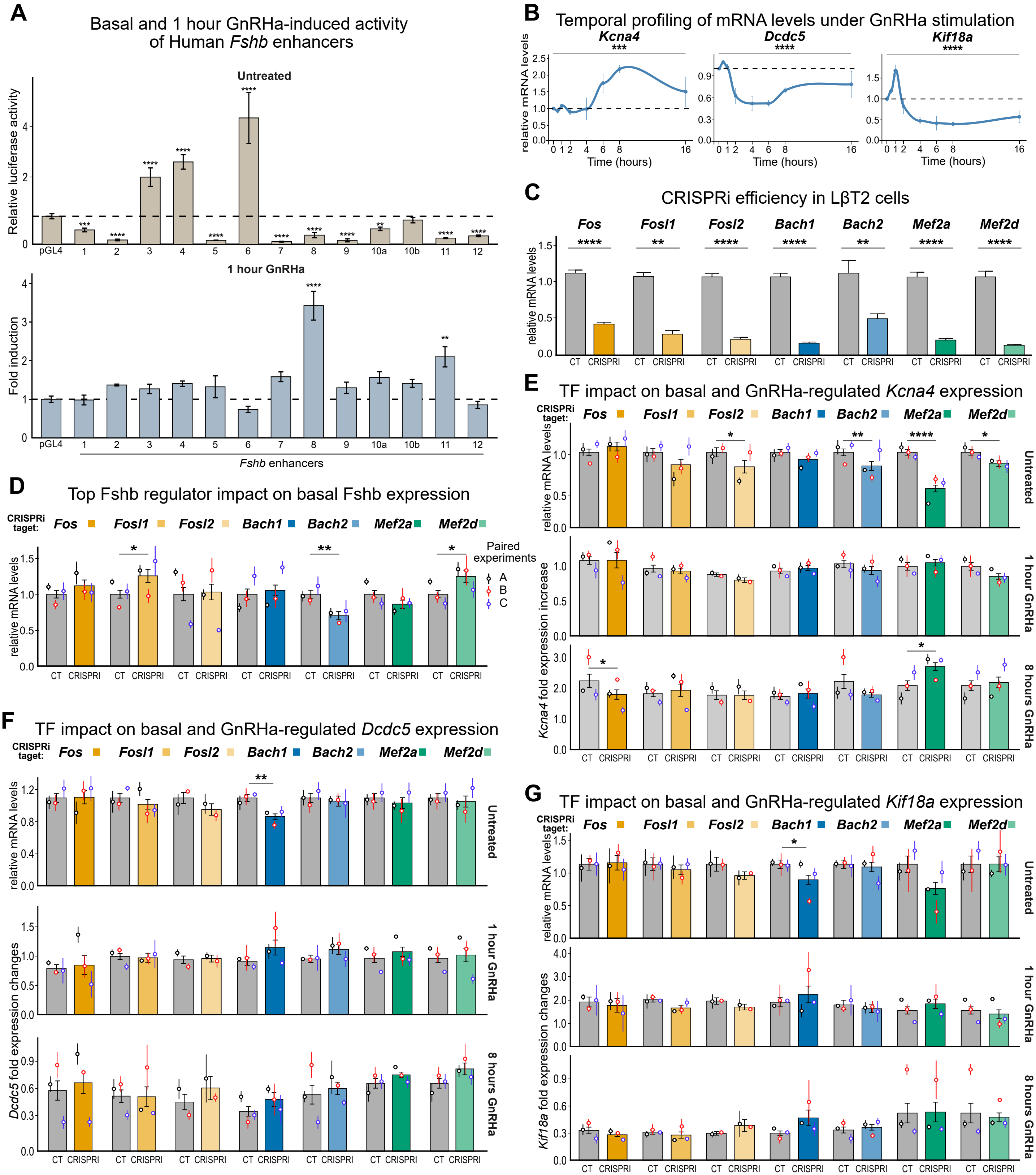
